# UB-MBX-46 is a potent and selective antagonist of the human P2X7 receptor developed by structure-based drug design

**DOI:** 10.1101/2025.02.12.637899

**Authors:** Adam C. Oken, Andreea L. Turcu, Eva Tzortzini, Kyriakos Georgiou, Jessica Nagel, Marta Barniol-Xicota, Ga-Ram Kim, So-Deok Lee, Annette Nicke, Yong-Chul Kim, Christa E. Müller, Antonios Kolocouris, Santiago Vázquez, Steven E. Mansoor

## Abstract

The P2X7 receptor is an ATP-gated ion channel that activates inflammatory pathways involved in diseases such as cancer, atherosclerosis, and neurodegeneration. However, despite the potential benefits of blocking overactive signaling, no P2X7 receptor antagonists have been approved for clinical use. Interspecies variation among existing antagonists has proven challenging, in part due to the dearth of molecular information on different receptor orthologs. Here, to identify distinct molecular features in the human receptor, we determine high-resolution cryo-EM structures of the full-length wild-type human P2X7 receptor in apo closed and ATP-bound open state conformations and draw comparisons with new and existing structures of other orthologs. We also report a cryo-EM structure of the human receptor in complex with an adamantane-based inhibitor, which we leverage, in conjunction with functional data and molecular dynamics simulations, to design a potent and selective antagonist with a unique polycyclic scaffold. Functional and structural analysis reveal how this optimized ligand, UB-MBX-46, interacts with the classical allosteric pocket of the human P2X7 receptor with picomolar potency and high selectivity, revealing its significant therapeutic potential.

## Introduction

When present in high concentrations, such as in pathological inflammatory states, extracellular ATP acts as a danger signal to cells by binding to and activating the P2X7 receptor (P2X7R)^1–5^. This spatiotemporal signaling stimulates the innate immune system by triggering assembly of the NLRP3 inflammasome and subsequent cytokine release, as well as activation of various less understood signaling cascades, and ultimately apoptosis^6–12^. Modulation of P2X7R-mediated cellular responses has the potential to treat diseases such as atherosclerosis in the cardiovascular system, Alzheimer’s disease in the central nervous system, as well as autoimmune diseases, infections, and select cancers in the immune system^4,13–21^. For these reasons, the P2X7R is a particularly important drug target.

While only few antagonists target the orthosteric binding site, there are numerous chemically distinct allosteric P2X7R antagonists, with wide-ranging potencies across orthologs. These ligands feature scaffolds containing adamantane or heterocycles such as tetrazole, pyridine or quinoline (Extended Data Fig. 1)^22–24^. But although several P2X7R antagonists have progressed to clinical trials and safety has been well established, the outcomes have been disappointing, and none have reached the market^25,26^. This could be partially attributed to the species-specific P2X7R expression patterns and variable pharmacological effects that each antagonist has on P2X7Rs from different species, complicating the predictability and translation of animal data to human efficacy. For instance, the P2X7R antagonist AZD9056, which reached phase II trials for rheumatoid arthritis (RA), ultimately showed limited efficacy, highlighting the challenges for extrapolating the data from the animal models to human outcomes^25^. Other examples of P2X7R antagonists that have shown variations in activity across orthologs include JNJ47965567, which has ∼10-fold higher affinity for the rP2X7R than the human P2X7R (hP2X7R) (Extended Data Fig. 1)^27^. In contrast, AZ11645373 is ∼100-fold less potent at the mouse P2X7R (mP2X7R), and >500-fold less effective against the rP2X7R, than the hP2X7R (Extended Data Fig. 1)^28,29^. There are P2X7R antagonists, such as A438079, that have equal inhibitory activity across all three orthologs but are generally less potent compared to more ortholog-specific antagonists (Extended Data Fig. 1)^5,30^.

As a result, there is a therapeutic need for small molecules that specifically and potently modulate hP2X7Rs while maintaining sufficient potency in rodents required for preclinical studies. Structure-based drug design could meet this need, but our poor molecular understanding of the pharmacological variability between P2X7R orthologs has delayed progress^25,31,32^. Structural investigations of the truncated panda P2X7R (pdP2X7R) and the full-length rP2X7R provided initial insight into the mechanism of allosteric antagonism of P2X7Rs^33,34^. P2X7R antagonists bind to either classical or extended allosteric ligand-binding sites that exist within the extracellular domain at the interface of two protomers (Extended Data Fig. 2)^33,34^. Such antagonists have been classified into shallow, deep, or starfish binders, depending on their functional properties and the residues within distinct allosteric ligand-binding sites with which they interact^34^. For example, JNJ47965567 is a deep-binding ligand that occupies the classical allosteric binding site whereas methyl blue is a starfish-binding ligand that occupies the extended allosteric binding site (Extended Data Fig. 1, 2)^34^. Although these data define the overarching principles of P2X7R antagonism, a comprehensive description of the ligand-binding sites and the associated molecular pharmacology that distinguishes different P2X7R orthologs remains enigmatic.

Here, to guide the development of hP2X7R ligands for therapeutic intervention, we use structural, biophysical, and electrophysiological methods to describe the molecular pharmacology of full-length wild-type rat, mouse, and human P2X7Rs. Our cryo-EM structures of the mP2X7R and the hP2X7R in the apo closed state, and the hP2X7R in the ATP-bound open state, identify ortholog-specific conformational differences and previously uncharacterized cholesterol binding sites. Further, through comparison with our previously published structure of the rP2X7R, we reveal differences in the classical allosteric ligand-binding site that underlie the pharmacological diversity between these orthologs^35,36^. We also determine the cryo-EM structure of the adamantane-based inhibitor UB-ALT-P30 bound to the hP2X7R, revealing the potential for larger scaffold ligands to bind within the classical allosteric pocket, and leverage structure-based drug design to develop a potent and specific scaffold for the hP2X7R. With this knowledge, we design and synthesize several antagonist scaffolds, and identify a promising compound, UB-MBX-46, for functional and cryo-EM analysis. UB-MBX-46 binds to the hP2X7R with picomolar potency and high selectivity, and thus defines a new ligand scaffold with the potential to treat P2X7R-associated diseases.

## Results

### P2X7R orthologs have distinct allosteric binding sites

To characterize the molecular differences between key mammalian P2X7R orthologs, we obtained cryo-EM structures of the full-length wild-type mP2X7 (2.5 Å) and hP2X7 (2.5 Å) receptors in the apo closed state conformation (Fig. 1A, Extended Data Fig. 3,4, Extended Data Table 1). The overall architecture of this P2X receptor (P2XR) subtype is consistent across human, mouse, and rat (PDB: 8TR5) orthologs (mean RMSD = 0.7 Å between Cα carbons)^35^. Conserved features include their trimeric architecture, the 3_10_-helices forming a closed gate, a partially hydrated sodium ion present in the closed pore, palmitoylation of residues on the C-cys anchor, and zinc and guanosine nucleotide-binding sites in the cytoplasmic ballast (Fig. 1, Extended Data Fig. 2A, 5)^35,36^. However, we also identified ortholog-specific features in the hP2X7R, including two cholesterol hemisuccinate (CHS) binding sites per protomer at the interface between transmembrane helix 1 (TM1) and transmembrane helix 2 (TM2) on the extracellular leaflet of the membrane bilayer (Fig. 1B-D). The two CHS molecules are stacked on top of each other such that the inner molecule is closer to the TM1/TM2 interface and the outer molecule is positioned between TM1 and the inner molecule (Fig. 1C,D). The inner CHS molecule is coordinated by hydrophobic interactions with residues F328 and L333 on TM2; F38, V41, C42, L45, and Y51 on TM1; and W265 on the lower body domain (Fig. 1C, Extended Data Fig. 2A). Two oxygen atoms on the hemisuccinate group of the inner CHS molecule also form hydrogen bonds with the sidechain of R264 and backbone nitrogen of W265 (distances of 2.5 Å and 2.9 Å, respectively) (Fig. 1C). The outer CHS molecule is coordinated by hydrophobic interactions with residues F38, C42, and V46 on TM1; residue F266 on the lower body domain; and the inner CHS molecule (Fig. 1D, Extended Data Fig. 2A). There are also hydrogen bonds between the hemisuccinate group of the outer CHS molecule and the sidechains of N261, H268, and K49 (distances of 2.8 Å, 3.4 Å, and 3.3 Å, respectively) (Fig. 1D). Although all three ortholog reconstructions are of similar resolution (∼2.5 Å), there are no densities in either the rP2X7R or the mP2X7R reconstructions to support modeling of CHS molecules, suggesting that these bound CHS molecules are specific to the hP2X7R.

**Figure 1:**
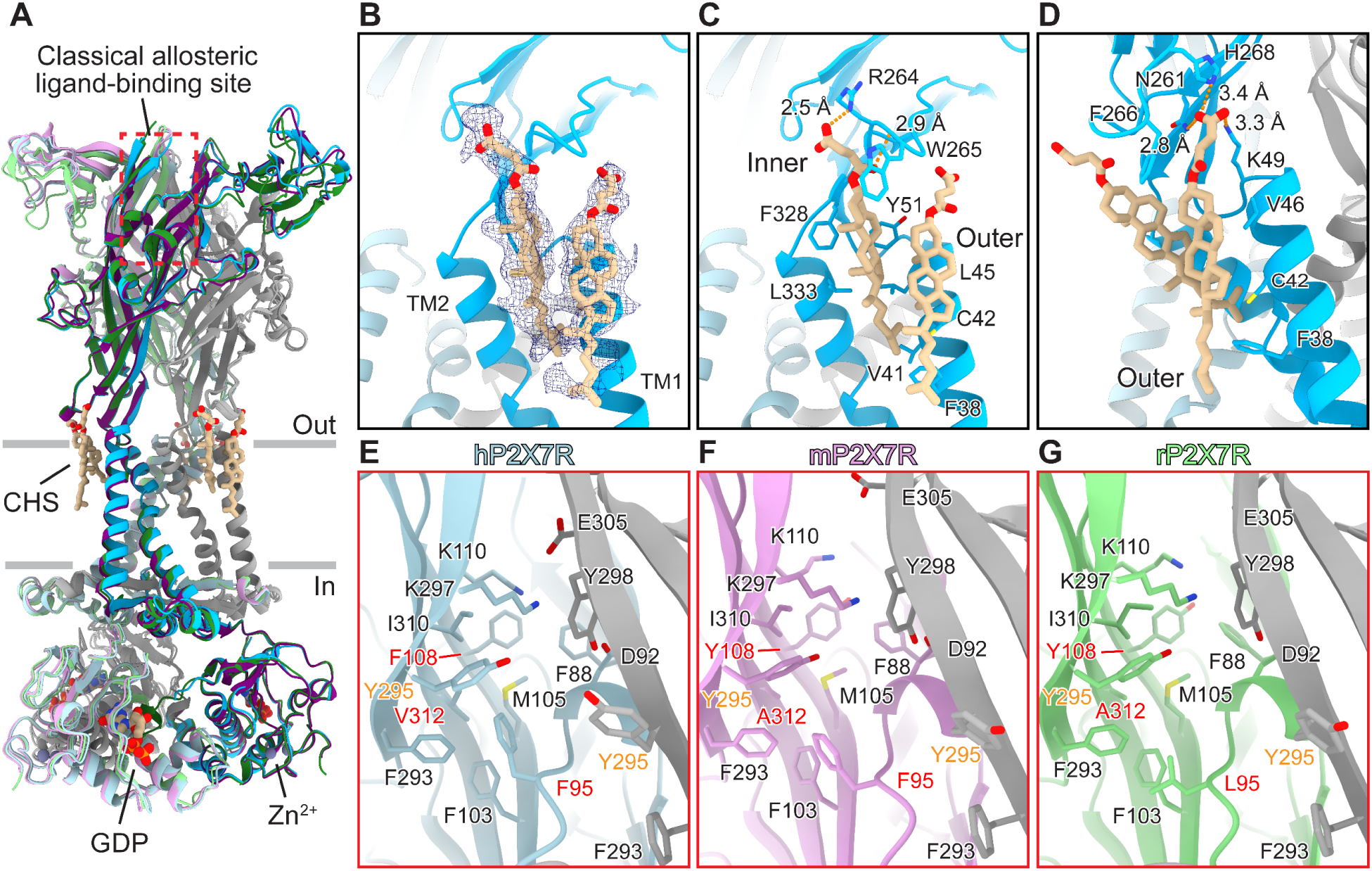
The apo closed state structures of human, mouse, and rat P2X7Rs reveal distinct ligand-binding sites. (**A**) Ribbon representation of the apo closed state structures of human (shades of blue and gray), mouse (shades of purple and gray), and rat (shades of green and gray) P2X7Rs (PDB code: 8TR5)^35^. The classical allosteric ligand-binding site is boxed in red. Two CHS molecules per protomer (inner and outer; both tan), located at the interface between TM1 and TM2 on the extracellular side of the membrane, are found only in the hP2X7R. GDP (tan) and Zn^2+^ ions (slate gray) are labeled and shown within the cytoplasmic ballast (Extended Data Fig. 1). (**B**) Magnified and 120° rotated view of **A** highlighting density for two molecules of CHS bound to one protomer of the hP2X7R. (**C**) Same view as **B** highlighting the hydrophobic and hydrogen bonding interactions between the inner CHS molecule and the hP2X7R. (**D**) 50° rotated view from **C** highlighting the hydrophobic and hydrogen bonding interactions between the outer CHS molecule and the hP2X7R. (**E-G**) Magnified view of the classical allosteric ligand-binding site from **A** highlighting the differences between human (light blue and gray), mouse (light purple and gray), and rat (light green and gray) P2X7R orthologs^35^. Residues 95, 108, and 312 (red) are the key residue differences in the classical allosteric pockets between the three orthologs. The larger V312 in hP2X7Rs (A312 in mP2X7Rs and rP2X7Rs) forces the neighboring residue Y295 (orange) to adopt an alternative rotameric conformation that slightly expands the classical allosteric pocket only in the human ortholog.

Comparison of these three structures also reveals molecular differences in the unoccupied classical allosteric ligand-binding sites (Fig. 1A,E-G)^35^. Across the three orthologs, the classical allosteric pocket comprises multiple conserved residues, including F88, D92, F103, M105, K110, F293, K297, Y295, K297, Y298, E305, and I310 (Fig. 1E-G, Extended Data Fig. 6). However, the identity of three residues within the classical allosteric pocket are different between human, mouse, and rat orthologs as seen in sequence alignments and visualized in their respective structures (Fig. 1E-G, Extended Data Fig. 6). First, F95 in hP2X7Rs and mP2XRs correlates to L95 in rP2X7Rs (Fig. 1E-G, Extended Data Fig. 6). This residue occupies the same general location and rotameric orientation in all three orthologs, but the phenylalanine found in the hP2X7R occupies much more space within the classical allosteric pocket^35^. Second, F108 in hP2X7Rs correlates to Y108 in both mP2X7Rs and rP2X7Rs (Fig. 1E-G, Extended Data Fig. 6). Again, this residue occupies the same position and rotameric conformation in all three structures, likely playing a similar role (Fig. 1E-G)^35^. Finally, V312 in hP2X7Rs correlates to A312 in both mP2X7Rs and rP2X7Rs (Fig. 1E-G, Extended Data Fig. 6). The larger valine sidechain in the human ortholog occupies more space within the classical allosteric pocket than the alanine in rat or mouse (Fig. 1E-G, Extended Data Fig. 6)^35^. Interestingly, although not apparent from sequence alignments, our structure of the hP2X7R shows that the larger sidechain of V312 forces the adjacent, conserved residue Y295 to adopt an alternative rotameric conformation, expanding the size of the pocket (Fig. 1E-G). Thus, the classical allosteric ligand-binding site in the human ortholog has a different shape which impacts the binding of antagonists.

### P2X7R orthologs have distinct orthosteric ATP-binding sites

Because pharmacological tools that activate human, mouse, and rat P2X7Rs are distinct, we examined the molecular determinants of agonism in the human ortholog to facilitate the development of higher-affinity agonists^30,35–37^. In agreement with previous measurements, two electrode voltage clamp (TEVC) recordings of human, mouse, and rat P2X7Rs revealed half maximal effective concentrations (EC_50_) for ATP of 89 ± 8.3 µM, 70 ± 17 µM, and 34 ± 8.4 µM, respectively (Fig. 2A)^35,36,38^. The modestly lower apparent affinity of ATP for the hP2X7R was borne out by direct measurements of kinetics and equilibrium binding affinities using bio-layer interferometry (BLI). The rate constant for association (k_a_) of 7.4 ± 1.3 ×10^4^ M^−1^s^−1^, and rate constant for dissociation (k_d_) of 4.8 ± 1.2 ×10^−2^ s^−1^, are ∼27% and ∼6% slower than those for ATP binding to the rP2X7R (Fig. 2B)^35^. This corresponds to ATP binding to the hP2X7R with an equilibrium dissociation constant (K_D_ = 650 ± 120 nM) that is approximately 20% lower than that of the rP2X7R (K_D_ = 540 ± 230 nM) (Fig. 2B)^35^.

**Figure 2:**
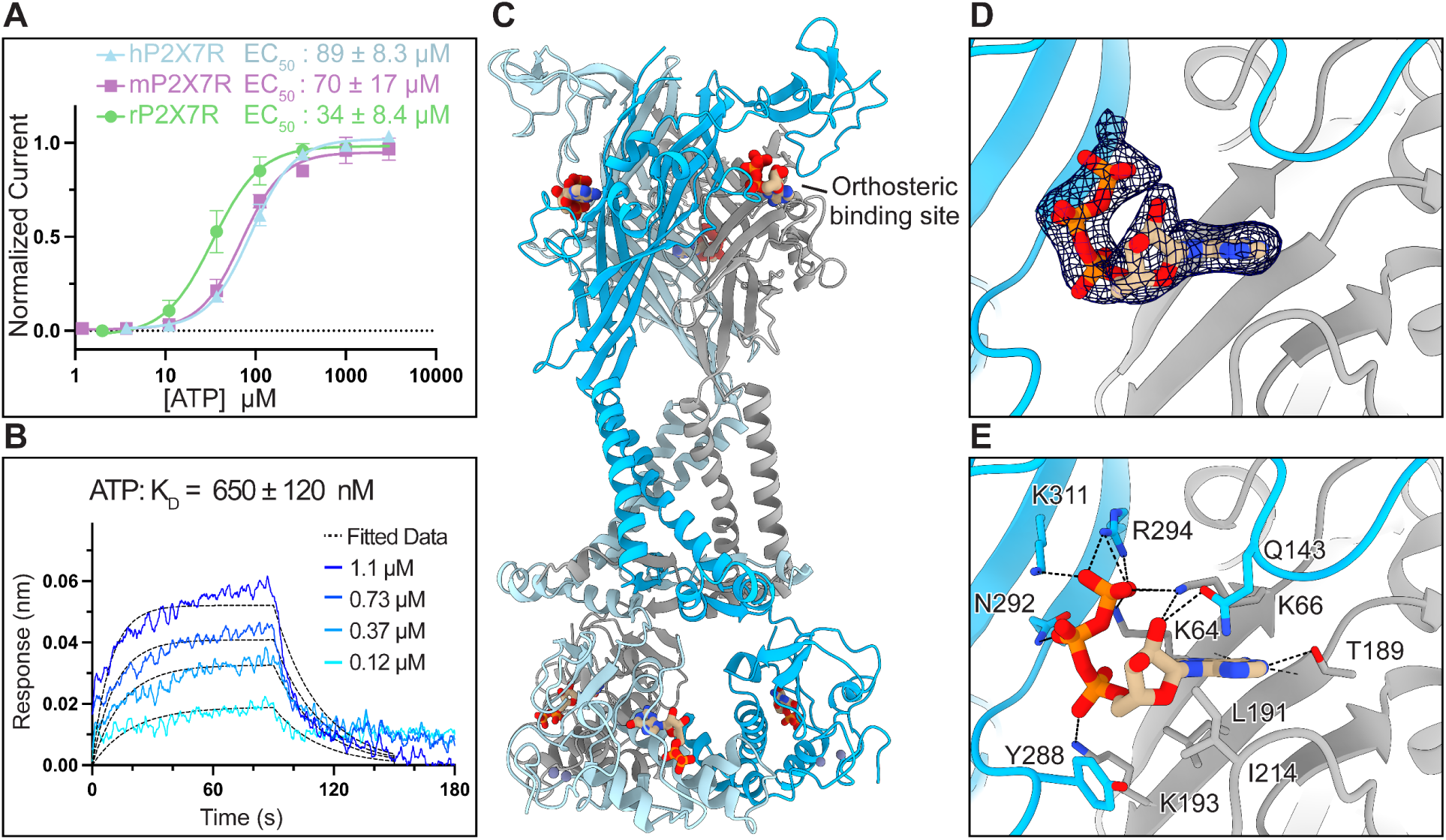
ATP binds to the orthosteric pocket of the hP2X7R. (**A**) Dose response curves from TEVC experiments measuring the activation of full-length wild-type human (blue), mouse (purple), and rat (green) P2X7Rs by ATP (EC_50_ = 89 ± 8.3 µM, 70 ± 17 µM, and 34 ± 8.4 μM, respectively). Data points and error bars represent the mean and standard deviation of normalized current, respectively, across triplicate experiments. (**B**) Representative BLI sensorgram for a dilution series of ATP (shades of blue) binding to biotinylated hP2X7R immobilized on streptavidin (SA) biosensors. Kinetic data were globally fit using a Langmuir 1:1 model to determine the equilibrium dissociation constant (K_D_) of ATP to the hP2X7R as K_D_ = 650 ± 120 nM, representing the mean and standard deviation across triplicate experiments. For kinetic analysis, a 90-second association time and a 60-second dissociation time were used for analysis. (**C**) Ribbon representation of the ATP-bound open state structure of the hP2X7R solved to 3.0 Å colored by protomer (blue, light-blue, and grey) highlighting the orthosteric ATP-binding site. (**D-E**) Magnified view of the orthosteric ATP-binding site from **A** highlighting the cryo-EM density for ATP (**D**) and residues interactions that coordinate ATP (**E**). ATP is coordinated by seven residues conserved across all subtypes (K64, K66, T189, K193, N292, R294, and K311) and four subtype-specific residues (Q143, L191, I214, and Y288).

To gain molecular insight into the binding of ATP to hP2X7Rs, we determined the cryo-EM structure of the full-length wild-type hP2X7R in the ATP-bound open state at 3.0 Å resolution (Fig. 2C-E, Extended Data Fig. 3,4, Extended Data Table 1). As expected, the ATP-bound structure of the hP2X7R has a global architecture similar to the rP2X7R in the ATP-bound (RMSD = 0.8 Å at Cα carbons, PDB: 6U9W) and BzATP-bound (RMSD = 1.0 Å at Cα carbons, PDB: 8TRJ) open state conformations (Extended Data Fig. 7)^35,36^. The minimum pore radius of the hP2X7R in the ATP-bound open state is 2.5 Å, the same as the rP2X7R in the ATP-bound open state, and large enough to pass partially hydrated sodium ions such as those present in the closed pores of human, mouse, and rat P2X7Rs^36,39^. Although there are structural similarities within the extracellular domains, the pore and cytoplasmic domains of the hP2X7R in the ATP-bound open state are rotated in comparison to the rP2X7R (Extended Data Fig. 7). Relative to the rP2X7R, TM1 of the hP2X7R is rotated up in-plane by ∼7° and TM2 is rotated up in-plane by ∼6° with the hinges at the extracellular ends of TM1 and TM2, respectively (Extended Data Fig. 7C). These differences result in lateral displacements of 5.2 Å (distance between Cα carbons of residue 24) at the start of TM1 and 4.8 Å (distance between Cα carbons of residue 358) at the end of TM2 (Extended Data Fig. 7C). The rotation of TM2 is further propagated into a global rotation of the cytoplasmic ballast by ∼10° in the hP2X7R relative to the rP2X7R (Extended Data Fig. 7C). This twist in the pore and cytoplasmic domain, much like the tightening of a spring, decreases the overall height of the hP2X7R by ∼4 Å compared to the rP2X7R (distance between Cα carbons of residues 521 and 514), likely representing an inherent flexibility of the transmembrane and cytoplasmic domains in the ATP-bound open state conformation.

The orthosteric binding site of the hP2X7R is clearly visualized at 3.0 Å resolution (Fig. 2C,D). In the ATP-bound open state of the hP2X7R, ATP is coordinated by the seven residues that are conserved in the ATP-binding sites of all P2XR subtypes: K64, K66, T189, K193, N292, R294, and K311 (Fig. 2E, Extended Data Fig. 6). In addition, the hP2X7R-specific residues, L191, I214, and Y288, form hydrophobic interactions with the ribose group and the sidechain of Q143 interacts with the 2’-hydroxy on the ribose group (3.3 Å) (Fig. 2E)^26^. Of these subtype-specific residues, I214 and Y288 in hP2X7Rs are not conserved across orthologs (Extended Data Fig. 6). Residue Y288 in hP2X7Rs corresponds to a valine in mP2X7Rs and a phenylalanine in rP2X7Rs, causing differences that might affect the pharmacology of ATP binding (Extended Data Fig. 6). Residue I214 in hP2X7Rs and rP2X7Rs corresponds to a glycine in mP2X7Rs, introducing flexibility in the mouse ortholog that could explain some pharmacological differences (Extended Data Fig. 6)^30^. Indeed, it has previously been shown that R125, Q143, and I214 in the rP2X7R are the critical determinants of efficacy, potency, and full agonism for ATP compared to BzATP^35^. In our structure of the hP2X7R, residues R125, Q143, and I214 are in similar positions and rotameric conformations to the rP2X7R, including the flexible sidechain of R125, which is solvent-facing and stubbed at the Cβ carbon. These molecular details of the hP2X7R in the ATP-bound open state will be crucial for structure-based drug design of high-affinity P2X7R agonists.

### The classical allosteric pocket of the hP2X7R can fit larger cage alkyls

We sought to understand how an existing P2X7R antagonist binds to the human ortholog to determine its suitability as a starting candidate for structure-based drug design. We initially selected the adamantane-containing compound UB-ALT-P30 because of its simplicity, ease of synthesis, and potential for further functionalization^40^. UB-ALT-P30 consists of adamantyl and 2-chlorophenyl moieties at either end connected by a hydrazide linker (Fig. 3A, Extended Data Fig. 1). In calcium influx assays, UB-ALT-P30 has a half maximal inhibitory concentration (IC_50_) of 18 ± 8.3 nM for the hP2X7R, 120 ± 50 nM for the rP2X7R, and 150 ± 27 nM for the mP2X7R, establishing its greater potency for the hP2X7R (Fig. 3B). Moreover, using a concentration at which we can expect 100% inhibition of the hP2X7R (10 μM), UB-ALT-P30 is more selective for the hP2X7R than the human P2X1 receptor (hP2X1R, 36 ± 8% inhibition), the human P2X2 receptor (hP2X2R, 29 ± 7%), the human P2X3 receptor (hP2X3R, 86 ± 9%), and the human P2X4 receptor (hP2X4R, 44 ± 14%) (Fig. 3C). This strong selectivity and high potency for the hP2X7R render UB-ALT-P30 a good initial compound for ligand development and optimization.

**Figure 3:**
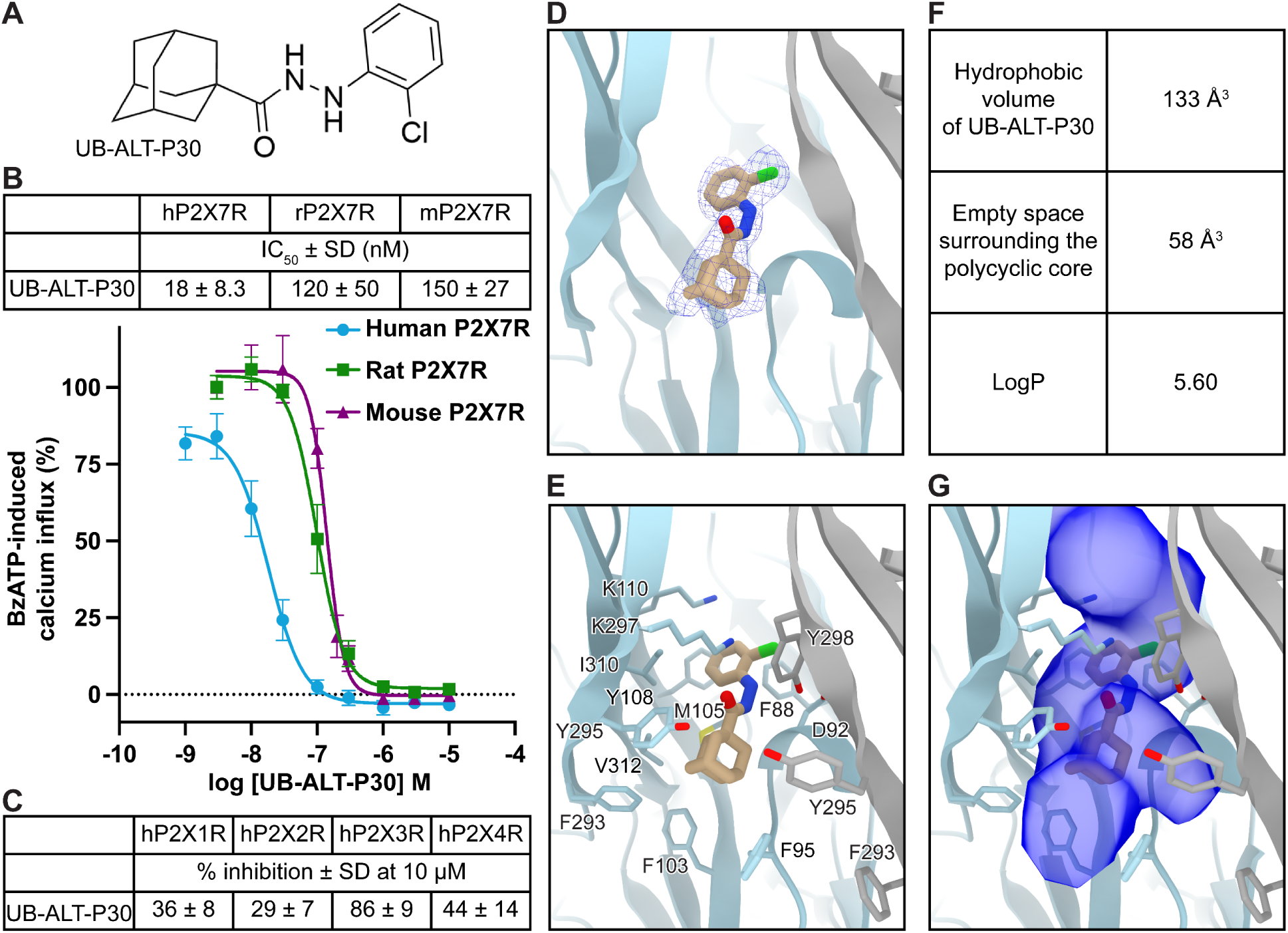
Cryo-EM analysis and molecular dynamics of UB-ALT-P30 bound to the hP2X7R show that larger caged alkyls fit the human classical allosteric pocket. (**A**) 2D chemical structure of UB-ALT-P30. (**B**) Concentration-dependent inhibition dose response curves (IC_50_) for UB-ALT-P30 on human, rat, and mouse P2X7Rs. Receptors were recombinantly expressed in 1321N1 astrocytoma cells and BzATP-induced calcium influx was normalized to an EC_80_. Data represent mean ± SD of at least three independent experiments performed in duplicates. (**C**) Calcium influx assays to measure the potency of UB-ALT-P30 for hP2X1Rs, hP2X2Rs, hP2X3Rs, and hP2X4Rs stably expressed in 1321N1 astrocytoma cells. Receptors were activated by ATP concentrations at their respective EC_80_ (hP2X1R, 100 nM; hP2X2R, 1,000 nM; hP2X3R, 100 nM; hP2X4R, 300 nM). Data represent mean ± SD from at least three independent experiments performed in duplicates. (**D**) Ribbon representation of the classical allosteric hP2X7R ligand-binding site, located at the interface of two protomers (gray and light blue) with one molecule of UB-ALT-P30 shown with corresponding electron density (blue mesh) at 2.8 Å. (**E**) Residues in the classical allosteric ligand-binding site that interact with UB-ALT-P30. The polycyclic group of UB-ALT-P30 forms hydrophobic interactions with the receptor. (**F**) Computational values that describe the binding properties of UB-ALT-P30 within the classical allosteric pocket of the hP2X7R. Hydrophobic volume is the hydrophobic surface of the adamantyl group calculated via Maestro; Schrödinger. Unoccupied space surrounding the adamantyl was calculated via POVME3.0^41^. LogP values calculated via Maestro; Schrödinger. (**G**) Same view as **E** showing the accessible volume of the classical allosteric pocket in the hP2X7R as calculated by Fpocket^70^. Several voids flanking UB-ALT-P30 as well as vacant space above and below the ligand can be occupied by further ligand development.

The high resolution cryo-EM structure of UB-ALT-P30 bound to the hP2X7R (2.8 Å) reveals receptor-ligand interactions and provides key insights to optimize ligand design (Fig. 3D,E, Extended Data Fig. 3, 4, Extended Data Table 1). UB-ALT-P30 binds to the classical allosteric ligand-binding site, at the interface of upper body domains from neighboring protomers, in a shallow binding pose (Fig. 1A, Extended Data Fig. 2)^34^. The binding of ligands to the classical allosteric pocket is thought to prevent receptor movements necessary for transition to the ATP-bound open state^33,34^. UB-ALT-P30 binds to the hP2X7R in a similar pose as other adamantane-containing antagonists, generating a pruned RMSD of 0.6 Å compared to the structure of AZD9056 bound to the rP2X7R (PDB code: 8TR8)^34^. However, important ortholog-specific interactions are also apparent.

The adamantyl moiety of UB-ALT-P30 is predominantly coordinated by hydrophobic interactions with the sidechains of residues F95, F103, M105, F293, Y295, and V312, which we confirmed with 500 ns molecular dynamics (MD) simulations (Fig. 3E, Extended Data Fig. 8A,C). Of these residues, F95 is specific to the hP2X7R ortholog and located on a dynamic loop in the upper body domain (residues 88-100) that is positioned differently in rat and human adamantane-based antagonist-bound structures (Fig. 3E, Extended Data Fig. 1, 2, 6)^34^. In the structure of AZD9056 bound to the rP2X7R, the corresponding L95 is rotated towards the ligand, making extensive hydrophobic interactions^34^. In contrast, F95 points away from UB-ALT-P30 in the hP2X7R, forming weak hydrophobic interactions and creating empty space below the molecule (Fig. 3E,F,G)^34^. Another ortholog-specific residue, V312, fills a hydrophobic cavity at one side of the adamantyl moiety in the hP2X7R (Fig. 3E, Extended Data Fig. 6). One additional hydrophobic cavity in the hP2X7R exists on the opposite side of the adamantyl moiety from V312, where three water molecules are coordinated by the backbone carbonyls of A296 (2.8 Å), R294 (3.2 Å), A91 (2.6 Å), and T94 (3.0 Å), as well as the backbone nitrogen of Y295 (3.3 Å). These cavities create ∼58 Å^3^ of empty hydrophobic space around the polycyclic core of UB-ALT-P30, presenting an opportunity to optimize this ligand by replacing the adamantane with more precisely fitting scaffolds (Fig. 3F,G).

Additional receptor-ligand interactions can be seen closer to the extracellular surface of the classical allosteric pocket in the hP2X7R. The hydrazide linker forms hydrogen bonds with the backbone carbonyl of D92 (2.8 Å) as well as the sidechain hydroxyl of Y298 (3.4 Å) (Fig. 3E, Extended Data Fig. 8A,C). The chlorophenyl moiety is predominantly coordinated by hydrophobic interactions with the sidechains of residues F88, M105, F108, and I310, as well as a cation-π interaction with the sidechain ammonium of K297 (Fig. 3E, Extended Data Fig. 8A,C). Residues F108 in the hP2X7R and Y108 in the rP2X7R appear to play the same role, forming edge-to-face interactions with ligands^34^. Together, the empirical knowledge gained from identifying ligand-receptor interactions and empty space around the adamantyl moiety of UB-ALT-P30 in the classical allosteric pocket of the hP2X7R can be used to design ligand scaffolds with higher potency and greater selectivity (Fig. 3F,G)^41^.

### UB-MBX-46 is a P2X7R antagonist with an improved scaffold

Given our detailed understanding of the molecular interactions between UB-ALT-P30 and the hP2X7R, we embarked on structure-based drug design to develop a more potent and selective antagonist. We replaced the adamantyl scaffold of UB-ALT-P30 with six alternative polycyclic cores of different sizes and shapes (Table 1). Although no other singular polycyclic hydrocarbon has the success rate of adamantane in medicinal chemistry, certain polycycles have outperformed it for specific targets, including some incorporated into P2X7R antagonists in clinical trials^26,42–46^. We therefore synthesized new compounds from the corresponding polycyclic carboxylic acids shown in Supplementary Fig. 1, following procedures reported for the synthesis of UB-ALT-P30^40,45,46^. All new compounds were characterized by their spectroscopic data, melting point, exact mass, and elemental analysis or HPLC/UV (Supplementary Figs. 1-5 and Supplementary Methods).

**Table 1:** Structure and characterization of P2X7R antagonists. IC_50_ for each compound on the hP2X7R was measured by ethidium bromide accumulation assays. Each value represents the mean and standard deviation of normalized responses across triplicate experiments. IC_50_ values of UB-MBX-P1 and UB-MBX-P2 have been reported previously^45^. RBFE values calculated with TI/MD method (ΔΔG_b,TI/MD_) for each perturbative transformation of UB-ALT-P30 to the analogs indicated in Table 1 represent mean and standard deviation for the three symmetry-related ligands bound to the receptor. Positive values indicate less favorable energetics and negative values indicate favorable energetics.

| Compound | Structure | IC<sub>50</sub> (nM) | $\Delta\Delta G_{exp}^a$<br>(kcal mol<sup>-1</sup>) | $\Delta\Delta G_{b,TI/MD}^b$<br>(kcal mol<sup>-1</sup>) | $\vert \Delta\Delta G_{b,TI/MD} - \Delta\Delta G_{exp}\vert ^c$<br>(kcal mol<sup>-1</sup>) |
| --- | --- | --- | --- | --- | --- |
| UB-MBX-46 |  | 1.0 ± 0.1 | -2.37 ± 0.32 | -2.77 ± 0.06 | 0.40 ± 0.16 |
| UB-ALT-P30 |  | 47 ± 4.2 | 0 | 0 | 0 |
| UB-ALT-P36 |  | 477 ± 170 | 1.43 ± 1.93 | 1.33 ± 0.07 | 0.10 ± 0.97 |
| UB-MBX-P2 |  | 18 ± 2 | -0.59 ± 0.37 | -1.56 ± 0.05 | 0.97 ± 0.07 |
| UB-ALT-P37 |  | > 1000 | > 1.88 | 2.66 ± 0.06 | < 0.78 |
| UB-MBX-P1 |  | 933 ± 156 | 1.84 ± 1.89 | 3.02 ± 0.07 | 1.18 ± 1.79 |
| UB-ALT-P38 |  | 430 ± 64 | 1.36 ± 1.50 | 4.32 ± 0.00 | 2.96 ± 1.13 |

<sup>a</sup> Experimental relative binding free energies ( $\Delta\Delta G_{b,exp}$ ) computed using the experimental pIC<sub>50</sub> values and equation (1), as described in Supplementary Methods; <sup>b</sup> correlation between $\Delta\Delta G_{b,TI/MD}$ and $\Delta\Delta G_{b,exp}$ values was 0.91; <sup>c</sup> $\mu_{ue} = 1.05$ kcal/mol (mean unsigned error computed the mean of $|\Delta\Delta G_{b,TI/MD} - \Delta\Delta G_{b,exp}|$ values).

The relative binding free energies (RBFEs) for each UB-ALT-P30 analog on the hP2X7R were determined *in silico* based on the corresponding perturbative transformation calculated using thermodynamic integration coupled with MD simulations (TI/MD) in phospholipid bilayers via Amber22 (Table 1)^47–50^. Of all our compounds, only UB-MBX-46 and UB-MBX-P2 resulted in negative RBFEs (ΔΔG_b,TI/MD_), indicating that the perturbative transformations of these two compounds were energetically favorable (Table 1). The inhibitory potency of each compound was experimentally tested using ethidium bromide accumulation assays (Table 1) and generally correlated with the computational predictions (UB-MBX-46 being the most potent compound, followed by UB-MBX-P2). Similarly, compounds with positive ΔΔG_b TI/MD_ values were less potent, including UB-MBX-P1 and UB-ALT-P37 (Table 1). These data indicate that UB-MBX-46 is the most promising compound for further validation. The polycyclic tetracyclo[4.4.0.0^3,9^.0^4,8^]decane scaffold of UB-MBX-46 features two cyclopentane rings in a “frozen” envelope conformation and two unique cyclohexane rings in boat conformation. This scaffold is larger than adamantane and cubane, yet smaller than the pentacyclic moiety of UB-MBX-P1 and is unique since it has scarcely been used in medicinal chemistry^46^.

### UB-MBX-46 is a potent and selective antagonist for the hP2X7R

To pharmacologically characterize the improved scaffold of UB-MBX-46, the compound was tested on different P2X7R orthologs and P2XR subtypes (Fig. 4A, Extended Data Fig. 1). In calcium influx assays, UB-MBX-46 has an inhibitory potency of 0.51 ± 0.04 nM for the hP2X7R and is ∼35x more potent on the hP2X7R, ∼3x more potent on the rP2X7R, and ∼33x more potent on the mP2X7R than the starting compound UB-ALT-P30 (Fig. 4B). Furthermore, UB-MBX-46 is more potent at the hP2X7R than either the rP2X7R (∼80-fold less potent) or the mP2X7R (∼8.9-fold less potent) (Fig. 4B). The selectivity of UB-MBX-46 for other P2XR subtypes was also tested using calcium influx assays. Using a concentration at which we can expect 100% inhibition of the hP2X7R (10 µM), UB-MBX-46 is more selective for the hP2X7R than the hP2X1R (13 ± 10% inhibition), the hP2X2R (15 ± 9%), the hP2X3R (29 ± 16%), or the hP2X4R (41 ± 19%) (Fig. 4C). Finally, UB-MBX-46 is even more selective for the hP2X7R compared to other P2XR subtypes than UB-ALT-P30 (Fig. 3C, 4C).

**Figure 4:**
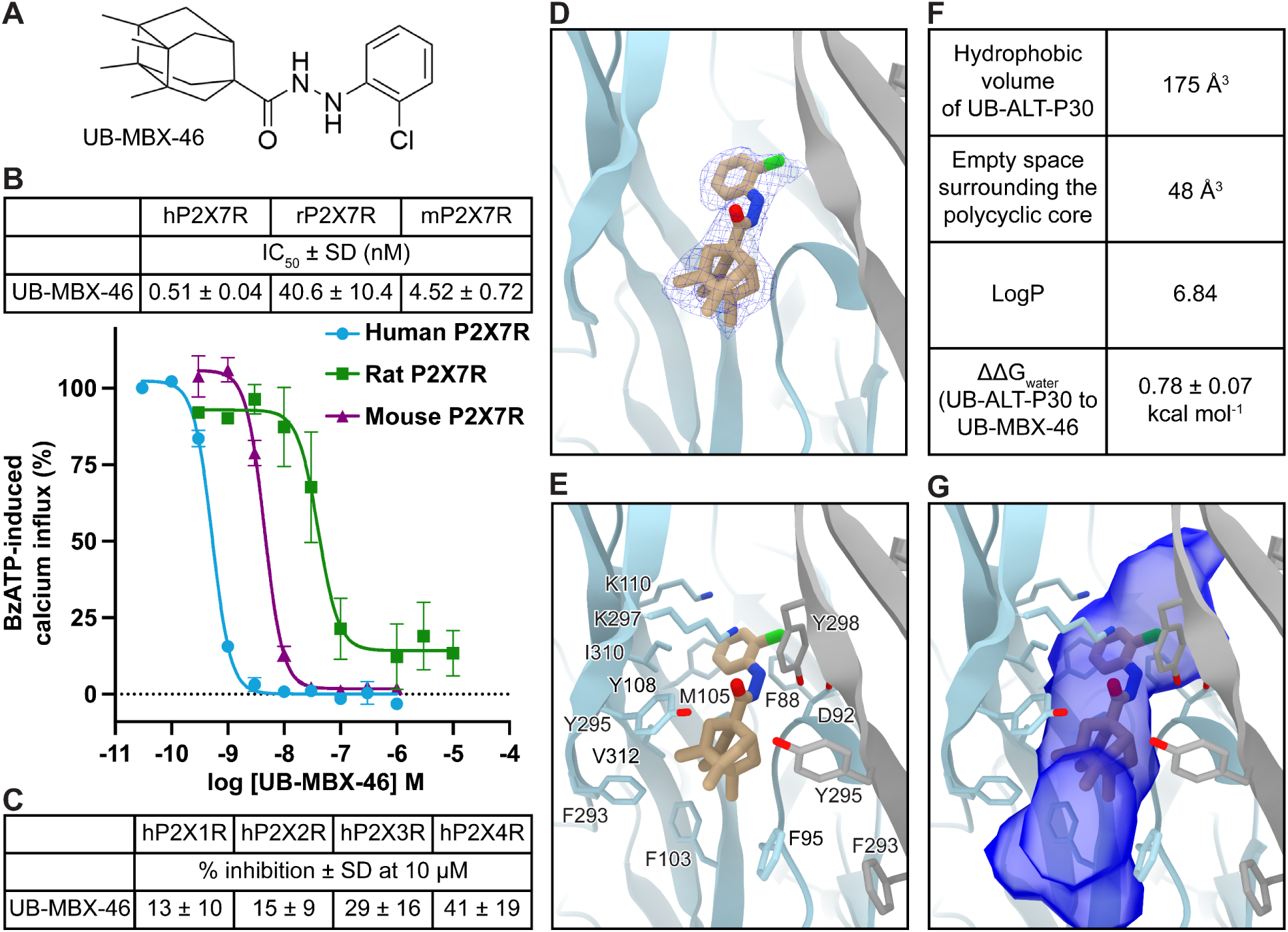
UB-MBX-46 is a potent and selective antagonist for the hP2X7R. (**A**) 2D chemical structure of UB-MBX-46. (**B**) Concentration-dependent inhibition dose response curves (IC_50_) for UB-MBX-46 on human, rat, and mouse P2X7Rs. Receptors were recombinantly expressed in 1321N1 astrocytoma cells and BzATP-induced calcium influx was normalized to an EC_80_. Data represent mean ± SD of at least three independent experiments performed in duplicates. (**C**) Calcium influx assays to measure the potency of UB-MBX-46 for hP2X1Rs, hP2X2Rs, hP2X3Rs, and hP2X4Rs stably expressed in 1321N1 astrocytoma cells. Receptors were activated by ATP concentrations at their respective EC_80_ (hP2X1R, 100 nM; hP2X2R, 1,000 nM; hP2X3R, 100 nM; hP2X4R, 300 nM). Data represent mean ± SD from at least three independent experiments performed in duplicates. (**D**) Ribbon representation of the classical allosteric hP2X7R ligand-binding site, located at the interface of two protomers (gray and light blue) with one molecule of UB-MBX-46 shown with its corresponding electron density (blue mesh) at 2.5 Å. (**E**) Residues in the classical allosteric ligand-binding site of the hP2X7R that interact with UB-MBX-46. The polycyclic group of UB-MBX-46 forms hydrophobic interactions with the receptor and the backbone carbonyl from D92 as well as the sidechain hydroxyl of Y298 form hydrogen bonding interactions with the hydrazide linker. (**F**) Computational values that describe the binding properties of UB-MBX-46 within the classical allosteric pocket of the hP2X7R. Hydrophobic volume is the hydrophobic surface of the polycyclic group calculated via Maestro, Schrödinger. Unoccupied space surrounding the polycyclic core was calculated via POVME3.0^41^. LogP values calculated via Maestro, Schrödinger. (**G**) Same view as **E** showing the accessible volume of the classical allosteric pocket in the hP2X7R calculated by Fpocket^70^. UB-MBX-46 tightly fits the center of the classical allosteric pocket of the hP2X7R, although unoccupied voids remain above and below the ligand.

We next investigated the molecular basis of the improved functional characteristics of UB-MBX-46 by determining the high resolution cryo-EM structure of UB-MBX-46 bound to the hP2X7R at 2.5 Å resolution (Fig. 4D,E, Extended Data Fig. 3,4, Extended Data Table 1). Similar to the adamantane derivative, UB-MBX-46 binds to the classical allosteric ligand-binding site in a shallow binding pose (Fig. 1A, Extended Data Fig. 2)^34^. The two ligand-bound structures are similar, with a pruned RMSD of 0.5 Å, however the longer and larger scaffold of UB-MBX-46 relative to UB-ALT-P30 (175 Å^3^ vs 133 Å^3^) binds deeper into the classical allosteric pocket and extends closer to the extracellular surface of the receptor (Fig. 3E,F, 4E,F). The hydrazide linker and chlorophenyl moieties of UB-MBX-46 are ∼0.9 Å closer to the extracellular surface than UB-ALT-P30 (average measurements between equivalent chlorine atoms, nitrogen atoms on the hydrazide linker, and closest carbon atoms to the linker of the polycyclic moieties). Yet, due to the longer length of UB-MBX-46 and its larger polycyclic moiety, the ligand also extends deeper into the classical allosteric pocket by ∼1.3 Å. As a result, there is only ∼48 Å^3^ of empty hydrophobic space surrounding the polycyclic core of UB-MBX-46; 10 Å^3^ less than around UB-ALT-P30 (Fig. 3F,G, 4F,G)^41^.

The cryo-EM structures and MD simulations show that many of the receptor-ligand interactions associated with UB-ALT-P30 in the classical allosteric pocket of the hP2X7R are present with UB-MBX-46. Similar to the adamantyl moiety in UB-ALT-P30, the polycyclic core of UB-MBX-46 is predominantly coordinated by hydrophobic interactions with the sidechains of residues F95, F103, M105, F293, Y295, and V312 (Fig. 4E, Extended Data Fig. 8B,D). However, due to the ligand’s depth in the pocket, the larger polycyclic core of UB-MBX-46 is more able to form hydrophobic interactions with the human-specific residue F95. The hydrazide linker forms hydrogen bonding interactions with the backbone carbonyl of D92 (2.7 Å) as well as the sidechain hydroxyl of Y298 (3.4 Å and 3.1 Å), creating one more hydrogen bond than UB-ALT-P30 (Fig. 3E, 4E, Extended Data Fig. 8). Finally, the chlorophenyl moiety is predominantly coordinated by hydrophobic interactions with the sidechains of residues F88, M105, F108, W167, and I310 as well as a cation-π interaction with the sidechain ammonium of K297 (Fig. 4E, Extended Data Fig. 8B,D). UB-MBX-46 appears to occupy the classical allosteric pocket with these improved ligand-receptor interactions compared to UB-ALT-P30. Further, from TI/MD calculations, UB-MBX-46 also has a lower desolvation penalty to bind to the receptor, indicating a preference for the hydrophobic pocket of the receptor over the solvent phase (Fig. 4F). Thus, more favored hydrophobic interactions surrounding the polycyclic core, more favorable hydrogen bonding interactions, and a lower desolvation penalty contribute to the higher potency of UB-MBX-46.

## Discussion

The P2X7R is a promising therapeutic target for numerous pathological diseases but pharmacological differences between receptor orthologs coupled with a lack of structural information have hampered efforts to develop a selective allosteric antagonist against the hP2X7R using structure-based drug design. The structural and functional data that we present here provide insights into the molecular differences between full-length wild-type rat, mouse, and human P2X7Rs – the three most relevant orthologs for drug development – in apo closed and ATP-bound open state conformations. The structure of the hP2X7R in complex with UB-ALT-P30 also reveals ligand-receptor interactions within the classical allosteric pocket. We leveraged these data to optimize an antagonist scaffold tailored to the classical allosteric ligand-binding site in the human ortholog using structure-based drug design. After synthesizing and characterizing six antagonists with different polycyclic cores, we identified one (UB-MBX-46) that was highly potent and selective for the hP2X7R. High-resolution structures solved using cryo-EM confirmed that it binds to the classical allosteric binding site with optimized ligand-receptor interactions. Due to its picomolar potency and high selectivity, UB-MBX-46 is a promising compound with significant therapeutic potential for treating P2X7R-associated diseases.

CHS-binding sites were only evident in the apo closed state structure of the hP2X7R but not in the rP2X7R or the mP2X7R structures, nor in the ATP-bound open state structure of the hP2X7R. The CHS-binding sites are located on the extracellular leaflet of the membrane, similar to the apo closed state structure of the hP2X1R, and likely represent cholesterol binding sites *in vivo* (Extended Data Fig. 9A,B)^51^. While characterizing the hP2X7R, we have observed that varying levels of CHS in the solubilization and purification process affect the stability of the cytoplasmic domain (Extended Data Fig. 2A, 9C,D). Insufficient levels of CHS throughout detergent solubilization and purification led to cryo-EM reconstructions with weak density for the cytoplasmic domains, in contrast to our experience with the mouse and rat orthologs (Extended Data Fig. 9C,D). Upon 3D-classification of the hP2X7R, data can be split into two distinct reconstructions with nearly identical extracellular and transmembrane domains but with the absence or presence of density for residues before TM1 or after TM2 (Extended Data Fig. 2A, 9C,D). These include residues that form the cytoplasmic cap, the C-cys anchor, and the cytoplasmic ballast (Extended Fig. 2A). We speculate that CHS molecules rigidify the detergent micelle to stabilize the palmitoyl groups on the C-cys anchor, which are known to permanently anchor the cytoplasmic cap^36^. Interestingly, the CHS molecules do not appear to interact with the palmitoyl groups within the model of the hP2X7R, suggesting other potential functional roles for cholesterol. Indeed, it is known that P2X7R activation and signaling is sensitive to cholesterol, serving as a negative regulator of large-pore formation^52,53^. Thus, although we visualize the CHS-binding sites on the extracellular side of the lipid bilayer and have evidence that CHS stabilizes cytoplasmic domains on the intracellular side of the lipid bilayer, the functional effects of CHS on the hP2X7R need more investigation.

The identities of three residues and the rotameric conformation of a fourth residue within the classical allosteric pockets of human, mouse, and rat P2X7Rs confer ortholog-specific properties that explain why small molecules targeting P2X7Rs have significant interspecies variation. Among the three orthologs, residues F95, F108, and V312 in hP2X7Rs form the largest steric combination of amino acids. The larger sidechain of V312 forces an alternative rotameric conformation of Y295 in the hP2X7R compared to its conformation in the mP2X7R or the rP2X7R. As a result of these residue differences, the classical allosteric pocket of the hP2X7R is smaller than in the rP2X7R or the mP2X7R in the apo closed state (Fig. 5A-C). In the presence of an antagonist, each of these residues play an important role in coordinating ligand binding to either the rP2X7R or the hP2X7R (Extended Data Fig. 8C,D)^34^. Although no studies have investigated whether residues 95, 108, 295, and 312 affect ortholog-specific inhibition in functional assays, it is clear from the structures of all three orthologs that each residue impacts ligand coordination in the classical allosteric pocket. Moreover, the identities of residues 95, 108, 295, and 312 are known to affect the inhibitory potency of other ligands and are therefore important considerations for the structure-based design of antagonists to target the hP2X7R^54,55^.

**Figure 5:**
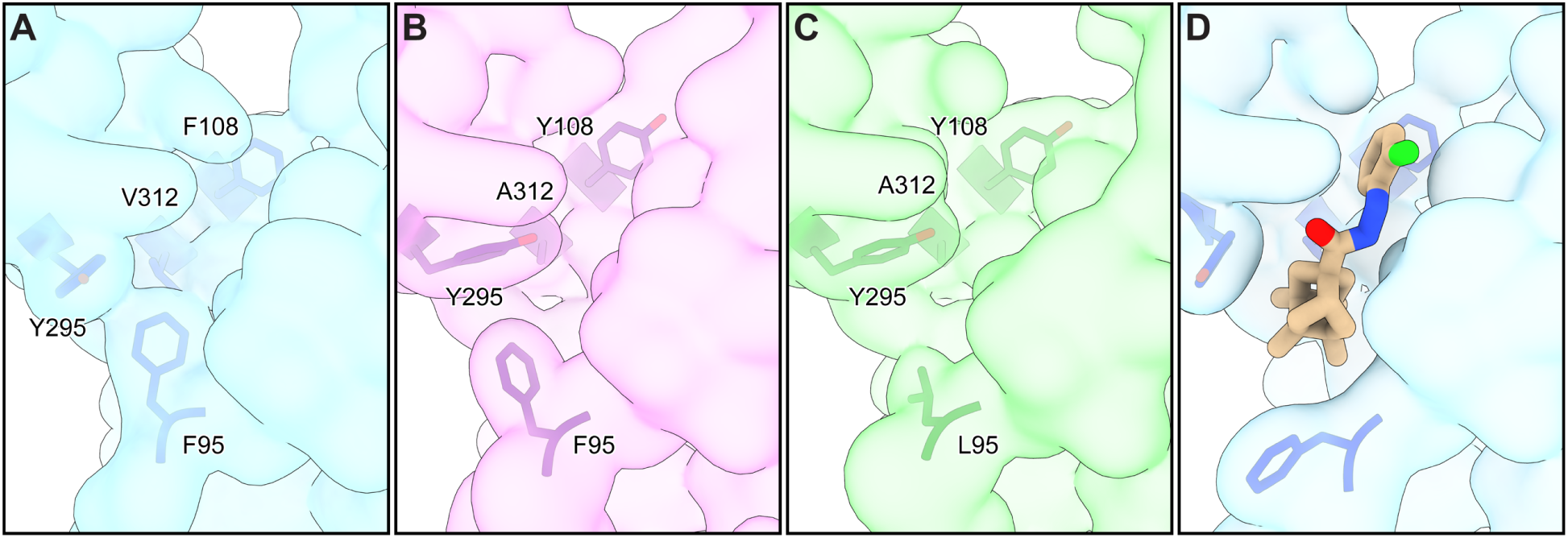
Potent and selective scaffolds for the allosteric modulation of the hP2X7R. (**A-D**) Schematic representation of the unoccupied classical allosteric pocket in three P2X7R orthologs and in the presence of UB-MBX-46 for the hP2X7R. Surface rendering for one P2X7 protomer is shown to highlight the size and identity of ortholog-specific residues comprising the classical allosteric pocket. (**A-C**) The classical allosteric pockets of the hP2X7R (**A**), the mP2X7R (**B**), and the rP2X7R (**C**) in the apo closed state conformation. The larger residues in the allosteric site of the hP2X7R appear to condense the pocket compared to the mP2X7R or the rP2X7R. Specifically, V312, F95, and the sterically rotated Y295 occupy more volume deep within the pocket. (**D**) The classical allosteric pocket of the hP2X7R in the presence of the potent and selective antagonist UB-MBX-46. The ligand is well tailored to the size and shape of the classical allosteric pocket in the human ortholog.

To examine the stereoelectronic requirements for ligand-binding in the classical allosteric pocket of the hP2X7R, we determined the cryo-EM structure of an adamantane-based inhibitor, UB-ALT-P30, bound to the hP2X7R. In agreement with MD simulations, we observed a poor fit of the molecule in the lipophilic cavity within the pocket and directed our efforts towards optimizing the polycyclic core of UB-ALT-P30, while retaining the hydrazide linker and chlorophenyl moiety, which appeared to have adequate interactions with the receptor. This motivation was triggered by recent research on polycyclic hydrocarbon structures in commercial drugs that occupy hydrophobic binding pockets and optimize drug potency^56^. Synthesis and testing of six different polycyclic moieties of different sizes and shapes revealed that a hydrophobic core that is either too big or too small results in poor inhibitory potency (Table 1). Initial efforts focused on adding methyl groups to the adamantane to enhance hydrophobic interactions while also filling the binding pocket with minimal steric hindrance. However, this approach resulted in a significant reduction of antagonistic activity. We also tested both smaller and larger alternative polycyclic structures. The larger of these permitted extensive surface interactions across the binding pocket but the much less spherical shape introduced steric clashes that reduced its overall inhibitory activity. Another compound containing a bisnoradamantyl scaffold with a smaller polycyclic system of two methyl groups provided efficient hydrophobic interactions and a two-fold increase in potency compared to UB-ALT-P30. Finally, the compound with the smallest polycyclic core lacks stabilizing hydrophobic interactions needed for effective binding within the classical allosteric pocket, rendering it a poor inhibitor.

Our most successful compound, UB-MBX-46, has an intermediate-sized scaffold. For this compound, we modified its tetracyclo[4.4.0.0^3,9^.0^4,8^]decane core with four strategically placed methyl groups, generating a tetracyclic unit with a well-balanced size and hydrophobic profile (Fig. 5D). Consequently, UB-MBX-46 has picomolar potency (IC_50_ = 510 ± 40 pM) and high selectivity for this P2XR subtype. The cryo-EM structure of the hP2X7R in complex with UB-MBX-46 and the associated MD simulations define the optimized ligand-receptor interactions that underlie its improved pharmacology (Fig. 5D). While unoccupied voids remain above and below the ligand suggesting further optimization is possible, the flanking regions of UB-MBX-46 tightly fit the lateral constrictions within the classical allosteric pocket of the hP2X7R. Together, these results show why it is crucial to perform structure-based drug design on human orthologs. Moreover, as a result of this effort, we have provided a picomolar P2X7R antagonist with an improved ligand scaffold that can serve as a basis to develop even more potent molecules with significant therapeutic potential.

## Methods

### Ethical statement

Unfertilized Xenopus laevis oocytes were purchased through Ecocyte Bioscience and kept at 18°C until injection. This research complies with all relevant ethical regulations. All surgical procedures for isolation of Xenopus laevis oocytes were done in accordance with animal welfare laws, followed national and institutional guidelines for humane animal treatment and complied with relevant legislation. Ecocyte Bioscience protocols help reduce the stress and harm on the laboratory animals, and appropriate aftercare such as pain management is employed to further minimize the impact of surgeries on the animals.

### Cell Lines

SF9 cells were cultured in SF-900 III SFM (Fisher Scientific) at 27°C. Cells of female origin were used for the expression of baculovirus.

HEK293 GNTI-and TSA201 cells were cultured using Gibco Freestyle 293 Expression Medium (Fisher Scientific) at 37°C supplemented with 2% v/v fetal bovine serum (FBS). HEK293 cells of female origin were used to express P2X7Rs.

Non-transfected human 1321N1 astrocytoma cells were purchased from Sigma-Aldrich (Taufkirchen, Germany). All stable cell lines were cultured at 37 °C and 5-10% CO_2_ in Dulbecco’s Modified Eagle Medium (Thermo Fisher Scientific, Dreieich, Germany) supplemented with 10% fetal calf serum, penicillin (100 U/mL), streptomycin (100 µg/mL), and the selection antibiotic geneticin G418 (800 µg/mL). All supplements were purchased from PAN Biotech (Aidenbach, Germany).

### Receptor Constructs

The full-length wild-type hP2X7R and mP2X7R constructs used for structure determination contain a C-terminal GFP, a 3C protease site, and a histidine affinity tag for purification. No mutations or truncations were made to the receptor for structure determination. For electrophysiology experiments, the hP2X7R-WT, mP2X7R-WT, rP2X7R-WT constructs are unmodified, full-length wild-type receptors with no GFP, protease sites, or affinity tags present.

For calcium influx assays, a chimeric hP2X1R was employed, and for the hP2X3R, the S15V-mutant was used to prevent fast receptor desensitization.

### Receptor Expression and Purification

The full-length wild-type hP2X7R and mP2X7R constructs were expressed by baculovirus mediated gene transfection (BacMam) using similar protocols as previously outlined for the rP2X7R^35,36^. Briefly, HEK293 GNTI^−^ or HEK293 cells were grown in suspension to a sufficient density and infected with P2 BacMam virus. Specifically, the ATP-bound and UB-ALT-P30 bound samples were prepared using HEK293 cells while both apo and UB-MBX-46 samples were prepared using HEK293 GNTI^−^ cells. After overnight growth at 37 °C, sodium butyrate was added (final concentration of 10 mM) and the HEK293 GNTI^−^ cells shifted to 30 °C for an additional 48 hours. For the HEK293 cells, no sodium butyrate was added and after overnight growth at 37 °C, the cells shifted to 30 °C for an additional 48 hours. The cells were then harvested by centrifugation, washed with PBS buffer (137 mM NaCl, 2.7 mM KCl, 8 mM Na_2_HPO_4_, 2 mM KH_2_PO_4_), suspended in TBS (50 mM Tris pH 8.0, 150 mM NaCl) containing protease inhibitors (1 mM PMSF, 0.05 mg/mL aprotinin, 2 µg/mL Pepstatin A, 2 µg/mL leupeptin), and broken via sonication. Intact cells and cellular debris were removed by centrifugation and then membranes isolated by ultracentrifugation. Membranes were snap frozen and stored at −80 °C until use.

When ready, membranes were thawed, resuspended in TBS buffer containing 15% glycerol, dounce homogenized, and then solubilized. For the mP2X7R, the homogenized membranes were solubilized in 40 mM dodecyl-β-D-maltopyranoside (DDM or C12M) and 8 mM cholesterol hemisuccinate tris salt (CHS) while the hP2X7R was solubilized in 40 mM DDM and ∼17 mM CHS. After ultracentrifugation, the soluble fraction was incubated with TALON resin in the presence of 10 mM imidazole at 4 °C for 1–2 h. After packing into an XK-16 column, the purification column was washed with 2 column volumes of buffer (TBS plus 5% glycerol, 1 mM C12M, 0.2 mM CHS at pH 8.0) containing 20 mM imidazole, 10 column volumes containing 30 mM imidazole, and eluted with buffer containing 250 mM imidazole. Peak fractions containing the protein were concentrated and digested with HRV 3-C protease (1:25, w/w) at 4 °C overnight. The digested protein was then ultracentrifuged and injected onto a Superose 6 10/300 GL column for size exclusion chromatography (SEC) that was equilibrated with 20 mM HEPES, pH 7.0, 100 mM NaCl, and 0.5 mM C12M. Fractions were analyzed by SDS-PAGE and fluorescence size exclusion chromatography (FSEC), pooled accordingly, and concentrated for cryo-EM grid preparation.

### Electron Microscopy Sample Preparation

To prepare ligand-bound samples, the purified receptor was incubated with ligand at 3-4x the molar concentration of the monomer for antagonists or 900 µM ATP. After a one-hour incubation shaking at 4 °C, cryo-EM grids were prepared for each sample. For all samples, 2.5 μL of solution was applied to glow-discharged (15 mA, 1 min) Quantifoil R1.2/1.3 300 mesh gold holey carbon grids and then blotted for 1.5 s under 100% humidity at 6 °C. The grids were flash frozen in liquid ethane using a FEI Vitrobot Mark IV and stored under liquid nitrogen until screening and large-scale data acquisition.

### Electron Microscopy Data Acquisition

Cryo-EM data for all receptor orthologs and complexes were collected on Titan Krios microscopes (FEI) operated at 300 kV at the Pacific Northwest Center for Cryo-EM (PNCC). Datasets were acquired on Gatan K3 direct-electron detectors in super-resolution mode using defocus values that ranged between −0.8 and −1.5 µm. Movies were collected with between 44 and 50 frames and a total dose ranging between 42 to 45 e^−^/ Å^2^. Datasets for both apo and antagonist-bound samples used an energy filter and nominal magnification of 130,000x, corresponding to a physical pixel size of ∼0.648 Å/pixel. On the other hand, the ATP-bound dataset did not use an energy filter and was collected at a nominal magnification of 37,000x, corresponding to a physical pixel size of ∼0.623 Å/pixel. Each dataset utilized ‘multi-shot’ and ‘multi-hole’ collection schemes driven by serialEM^57^.

### Electron Microscopy Data Processing

Cryo-EM movies were imported into cryoSPARC and patch motion correction was completed, outputting all micrographs at the physical pixel size (∼0.648 Å/pixel) (Extended Data Fig. 3, Extended Data Table 1)^58^. Following estimating of the contrast transfer function (CTF) parameters, micrographs were curated and particles picked using 2D templates generated from a 3D volume. Particles were inspected, extracted, and sent directly to 3D classification, skipping 2D classification (Extended Data Fig. 3, Extended Data Table 1). To remove bad particles, ab initio jobs generated references volumes that were used for iterative heterogeneous classifications, ultimately yielding the final particle stacks. The final homogeneous particle stacks were used for non-uniform refinements with global and local CTF refinements to yield the consensus cryo-EM reconstructions (Extended Data Fig. 3, 4, Extended Data Table 1)^59^.

### Model Building and Structure Determination

Homology models for apo closed state structures of the hP2X7R and the mP2X7R were generated from the apo closed state structure of the rP2X7R (PDB code: 8TR5) while the hP2X7R in the ATP-bound open state model was generated from the ATP-bound open state of the rP2X7R (PDB code: 6U9W) using SWISS-MODEL^35,36,60^. Each initial model was then docked into the corresponding cryo-EM map using ChimeraX^61,62^. Model building involved iterations of manual correction in COOT followed by refinements in PHENIX^61,62^. All ligands were built and refined using eLBOW with protonation states corresponding to approximately pH 7^63^. Limited glycosylation, acylation, and palmitoylation were included in models when justified by the corresponding electron density. Model quality was evaluated by MolProbity (extended Data Table 1)^64^.

### Two-electrode voltage clamping

#### Preparation of oocytes expressing P2X7Rs

Defolliculated *Xenopus Laevis* oocytes were purchased from Ecocyte Biosciences and resuspended in Modified Barth’s Solution (88 mM NaCl, 1 mM KCl, 0.82 mM MgSO_4_, 0.33 mM Ca(NO_3_)_2_·4H_2_O, 0.41 mM CaCl_2_·2H_2_O, 2.4 mM NaHCO_3_, 5 mM HEPES) supplemented with amikacin 250 mg/L and gentamycin 150 mg/L. Oocytes were then injected with 50 nL of either hP2X7R (40 ng/μL), mP2X7R (20 ng/μL), or rP2X7R (20 ng/μL) mRNA made from linearized full-length wild-type pcDNA 3.1x according to the protocol provided in the mMessage mMachine kit (Invitrogen). Injected oocytes were allowed to express for ∼20 hours before recording was performed.

#### TEVC recordings

Data acquisition was performed using an Oocyte Clamp OC-725C amplifier and pClamp 8.2 software. Buffers were applied using a gravity fed RSC-200 Rapid Solution Changer flowing at ∼5 mL/min. All experiments use Sutter filamented glass 10 cm in length with an inner diameter of 0.69 mm and an outer diameter of 1.2 mm to impale oocytes and clamp the holding voltage at −60 mV. Experiments were recorded in buffer containing 100 mM NaCl, 2.5 mM KCl, 0.1 mM EDTA, 0.1 mM flufenamic acid, and 5 mM HEPES at pH 7.4. All oocytes expressing P2X7R were facilitated with multiple applications of 100 μM ATP before any data was recorded.

#### Dose response (EC_50_) experiments

Excitatory responses to dilution series of ATP, ranging from 3 mM to ∼1 µM, were performed to determine the EC_50_ value. Each evoked response was normalized to the signal evoked by the largest concentration of ATP and the data was fitted in Prism 10 using the nonlinear regression named “EC_50_, x is concentration” to afford EC_50_ values. This value is then averaged amongst each singular condition and reported as the mean ± standard deviation.

### Calcium influx assays

The inhibitory potency of the antagonists UB-MBX-46 and UB-ALT-P30 was determined in calcium influx assays, as previously described^65,66^. For 1321N1 astrocytoma cells stably expressing the human, rat, or mouse P2X7Rs, or hP2X2Rs or hP2X4Rs, the fluorescent Ca^2+^-chelating dye Fluo-4 acetoxymethyl ester (Fluo-4 AM, Thermo Fisher Scientific, Dreieich, Germany) was used. For 1321N1 astrocytoma cell lines recombinantly expressing the hP2X1Rs or hP2X3Rs, the FLIPR^®^ Calcium 5 Assay Kit (Molecular Devices, San José, CA, USA) was employed. Human, rat, and mouse P2X7Rs as well as hP2X1Rs, hP2X2Rs, hP2X3Rs, and hP2X4Rs were recombinantly expressed in 1321N1 astrocytoma cells using a retroviral expression system^65–67^.

On the first day, 45,000 cells per well (30,000 cells for the human P2X3 receptor-expressing cells) were seeded into a 96-well polystyrene microplate with a black, flat bottom in a final volume of 200 µL. After overnight incubation at 37 °C and 10% CO_2_ (5% CO_2_ for the human P2X3 receptor-expressing cell line), the medium was removed by inverting the plate, and the cells were loaded with dye solution. Hanks’ balanced salt solution (HBSS, Thermo Fisher Scientific, Dreieich, Germany) was used as the assay buffer for human P2X1, P2X2, P2X3, and P2X4 receptors. For the P2X7Rs, a different assay buffer was used: 150 mM sodium-glutamate, 5 mM KCl, 0.5 mM CaCl_2_, 0.1 mM MgCl_2_, 10 mM D-glucose, 25 mM 4-(2-hydroxyethyl)-1-piperazineethanesulfonic acid (HEPES), pH 7.4. The dye solutions were freshly prepared. The Fluo-4 AM solution contained 3 µM of Fluo-4 AM in assay buffer and 1% of the non-ionic detergent Pluronic F-127^®^ (Sigma-Aldrich, Taufkirchen, Germany). The Calcium 5 dye, used for hP2X1Rs and hP2X3Rs, was prepared in HBSS buffer. Cells were loaded with Fluo-4 AM and incubated for 1h at room temperature with gentle shaking (100 rpm), while those loaded with the Calcium 5 dye were incubated for 1h at 37 °C. After incubation, the dye solution was carefully removed from the plate, and buffer was added to the cells for the evaluation of agonists. For determining EC_80_ values for ATP (Roth, Karlsruhe, Germany), and BzATP (Jena Bioscience, Jena, Germany), respectively, dilution series were prepared in transparent 96-well plates (Boettger, Bodenmais, Germany) using the respective assay buffer. To determine IC_50_ values of the antagonists, the dye solution was removed, and the dilution series of the antagonist dissolved in DMSO was added (final DMSO concentration: ≤1% for hP2X2Rs, hP2X4Rs, and hP2X7Rs, and ≤0.5% for hP2X1Rs and hP2X3Rs). The agonist solution was prepared in a 96-well transparent plate obtaining the respective EC_80_ values. As a control, assay buffer without test compound was used. After 30 min of incubation, the plates were measured using a fluorescence imaging plate reader NOVOstar (BMG Labtech GmbH, Offenburg, Germany) at an excitation wavelength of 485 nm and an emission wavelength of 520 nm, for 30 s at 0.4 s intervals. Calcium influx measurements were initiated by automatic addition of 20 µL of agonist solution by the plate reader’s pipetting device. Data were analyzed using Microsoft Excel and GraphPad Prism (Version 8.0, San Diego, CA, USA). EC_80_ and IC_50_ values were calculated using nonlinear regression with a sigmoidal dose-response equation.

### Measurement of ethidium bromide accumulation in hP2X7R-expressing HEK293 cells

Human P2X7R-expressing HEK293 cells were cultured in a humidified atmosphere of 5% CO_2_ at 37 ℃ in Dulbecco’s Modification of Eagle’s Medium (DMEM; Corning) supplemented with 10% (v/v) fetal bovine serum (Corning) and 1% (v/v) antibiotic–antimycotic (Gibco)^68^. To perform the assay, the DMEM was removed, and the HEK293 cells were washed with 3 mL of 1X DPBS (Dulbecco’s Phosphate-Buffered Saline; Corning). After the removal of DPBS solution, the cells were detached from the dish with 1 mL of trypsin/EDTA (Gibco), and the collected cells were subsided by centrifugation at 1000 rpm for 2 minutes. The cells counted with 2.25 × 10^7^ were re-suspended in 4-(2-hydroxyethyl)-1-piperazineethanesulfonic acid (HEPES)-buffered salt solution consisting of 140 mM potassium chloride, 1 mM ethylene diamine tetraacetic acid (EDTA), 5 mM glucose, 20 mM HEPES and 0.1 mM ethidium bromide (pH 7.4). The vehicles or appropriate range of concentrations of compounds (10 μL, pre-diluted in 10% (v/v) DMSO in DW from 10 mM stock) were added to each well of 96-well black plate (Corning), and 80 μL of the cell suspension was subsequently added to each well. BzATP (10 μL, pre-diluted in DW from 10 mM stock) was then added, and the final assay volume was maintained as 100 μL. After the incubation in 5% CO_2_ at 37 ℃ for 2 hours, the ethidium dye uptake was detected by measuring the fluorescence (excitation wavelength of 530 nm and emission wavelength of 590 nm) of each well using a CHAMELEON™ Multi-Technology Plate Reader. The antagonistic activities of compounds are expressed as the percentages relative to the maximum accumulation of ethidium bromide observed in the control group with the stimulation of the hP2X7R by an agonist, BzATP. The IC_50_ values of compounds as antagonisms were calculated using nonlinear regression analysis using OriginPro 9.1 software.

### Synthesis of P2X7R antagonists

Compounds were synthesized as depicted in Supplementary Fig. 1. A series of new acyl hydrazides was synthesized starting from the corresponding carboxylic acids, following similar procedures to the previously reported for the synthesis of UB-ALT-P30^40,46^. The final products were purified by crystallization. Identity and purity were confirmed by ^1^H and ^13^C nuclear magnetic resonance (NMR), high-resolution mass spectrometry, elemental analysis, infrared, melting point and HPLC/UV (Supplementary Fig. 2-5). See supplementary materials for detailed synthetic procedures, analytical data, NMR spectra and, when appropriate, HPLC traces to demonstrate identity and purity for all products.

### Molecular Dynamics simulations

The structures of f-hP2X7R in the apo closed state and f-hP2X7R in complex with UB-ALT-P30 or the UB-MBX-46 (with bound cholesterols) were utilized as starting models for 1 μs or 500 ns MD simulations at 310 K, respectively, with the Amber22 software^49^. Each protein structure after suitable preparation was inserted in a pre-equilibrated hydrated POPC bilayer expanding 30 Å from the furthermost vertex of the protein to the edge of the simulation orthorhombic box in all axes. The resulting lipid buffer contained ∼479,000 atoms, consisting of 584 POPC lipids and ∼123,500 water molecules. The dimensions of the simulation box were 150×148×226 Ǻ^3^. Periodic boundary conditions were applied. We used the ff19sb to model the protein, the lipid21 force field to model the POPC lipids, GAFF2 to model the ligand, and the TIP3P model for waters and ions. After equilibration phase including restrained energy minimization, NVT MD simulation steps with Langevin thermostat (dynamics) and NPT MD simulation steps with Berendsen barostat were performed. Bonds involving hydrogen atoms were constrained by the SHAKE algorithm and a time step of 1 fs and time of 2 fs with the leapfrog Verlet integrator was applied for unrestrained production runs at 310 K. Long range electrostatics were calculated using Particle-mesh Ewald summation (PME), with a 1 Å grid, and short-range non-bonding interactions were truncated at 12 Å. A known MD simulation protocol was applied^69^.

For the TI/MD calculations performed with Amber22 software, minimized geometries of the complexes were then used for simulations at all *λ* values^47,49^. Eleven *λ* values were applied, equally spaced between 0.0 to 1.0. Each MD simulation was heated to 310 K for 500 ps using the Langevin thermostat for temperature control. The Berendsen barostat was used to adjust the density. The 500 ps of NVT equilibration was followed by 2 ns NVT production simulation without restraints. Production simulations recalculated the potential energy at each *λ* value every 1 ps for later analysis with MBAR. A known MD simulation protocol was applied^69^. For each alchemical calculation of UB-ALT-P30 in complex with the full-length hP2X7R was applied dual topology and the 1-step protocol was performed which includes disappearing one ligand and appearing the other ligand simultaneously, and the electrostatic and van der Waals interactions are scaled simultaneously using softcore potentials from real atoms that are transformed into dummy atoms. Two repeats were performed for the 2 ns-TI/MD calculation for each alchemical transformation shown in Table 1. The alchemical calculation UB-ALT-P30 → UB-MBX-46 was performed also in the water and gas phase to account for the desolvation penalty. Particle Mesh Ewald Molecular Dynamics (pmemd) and energy minimization step was performed using the Central Processing Unit (CPU) of workstations. The rest of the equilibration steps including the unrestraint production were run with AMBER22 software on RTX 4090 GPUs in lab workstations using pmemd.CUDA algorithm^49^.

See supplementary materials for detailed molecular dynamics simulations methodology.

## Data Availability

All cryo-EM density maps for the full-length wild-type mP2X7R in the apo closed state and the full-length wild-type hP2X7R in the apo closed, ATP-bound open, and antagonist-bound inhibited states have been deposited in the Electron Microscopy Data Bank (EMDB) under accession codes: EMD-47490 (apo closed hP2X7R), EMD-47491 (ATP-bound open hP2X7R), EMD-47492 (UB-ALT-P30-bound inhibited hP2X7R), EMD-47493 (UB-MBX-46-bound inhibited hP2X7R), and EMD-47494 (apo closed mP2X7R). The maps within these depositions include both half maps, sharpened/unsharpened maps, refinement masks, and any local refinements or locally sharpened maps that helped with model building. The corresponding coordinates for the structures have been deposited in Protein Data Bank under the PDB accession codes 9E3M (apo closed hP2X7R), 9E3N (ATP-bound open hP2X7R), 9E3O (UB-ALT-P30-bound inhibited hP2X7R), 9E3P (UB-MBX-46-bound inhibited hP2X7R), and 9E3Q (apo closed mP2X7R). All active compounds are available from the authors on reasonable request.

## Acknowledgements

We thank O. Davulcu, C. Yoshioka, and C. López at PNCC for access and microscopy assistance. Electron microscopy grid screening was performed at the Multiscale Microscopy Core within Oregon Health & Science University (OHSU). BLI analysis was performed at the Biophysics Core within OHSU. We thank L. Anson for comments on the manuscript. A portion of this research was supported by NIH grant U24GM129547 and performed at the PNCC at OHSU and accessed through EMSL (grid.436923.9), a DOE Office of Science User Facility sponsored by the Office of Biological and Environmental Research. This research was supported by the National Heart, Lung and Blood Institute (R00HL138129, SEM), the National Institute of General Medical Sciences (DP2GM149551, SEM), and the American Heart Association (24PRE1195450, ACO). Part of this work was funded by the Spanish *Ministerio de Ciencia, Innovación y Universidades*, MICIU/AEI/10.13039/501100011033: grant PID2023-147004OB-I00 (to S.V.). The Kolocouris lab thanks Chiesi Hellas for funding (SERG No. 10354). C.E.M is grateful to the German Federal Ministery of Education and Research (BMBF) for support (Biopharma Neuroallianz, 0315606B), the Deutsche Forschungsgemeinschaft (SFB 1328) for supporting this work, and the COST Action CA21130 “P2X receptors as a therapeutic opportunity (PRESTO)”. AN is supported by the Deutsche Forschungsgemeinschaft (DFG, German Research Foundation) Project-ID 335447717 - SFB 1328 The Mansoor Lab would like to thank Barbara Allen, Jim Batzer, and Randy and Barbara Lovre for their generous support.

## Author Contributions

A.C.O., A.L.T, S.V., A.K., and S.E.M. designed the project. A.C.O. performed the cryo-EM sample preparation, data collection, data processing, and built the models. A.C.O. performed and analyzed the BLI experiments. A.C.O. performed and analyzed the TEVC electrophysiological experiments. J.N. and C.E.M. prepared stable cell lines expressing human, rat and mouse P2X7Rs, and performed the calcium influx assays, and analyzed the data. G.K., S.L., and Y.K. performed and analyzed the ethidium bromide accumulation experiments. A.L.T., M.B.X., and S.V. designed, synthesized, and purified the P2X7R antagonists. E.T., K.G., and A.K. performed and analyzed the MD simulations as well as additional calculations; E.T. and K.G. contributed equally. A.N. provided intellectual contributions. A.C.O. and S.E.M wrote the manuscript. A.C.O., A.L.T, A.N., J.N., C.E.M., A.K., S.V., and S.E.M. edited the manuscript. All authors approved the manuscript.

## Extended Data Figures

**Extended Data Fig. 1:**
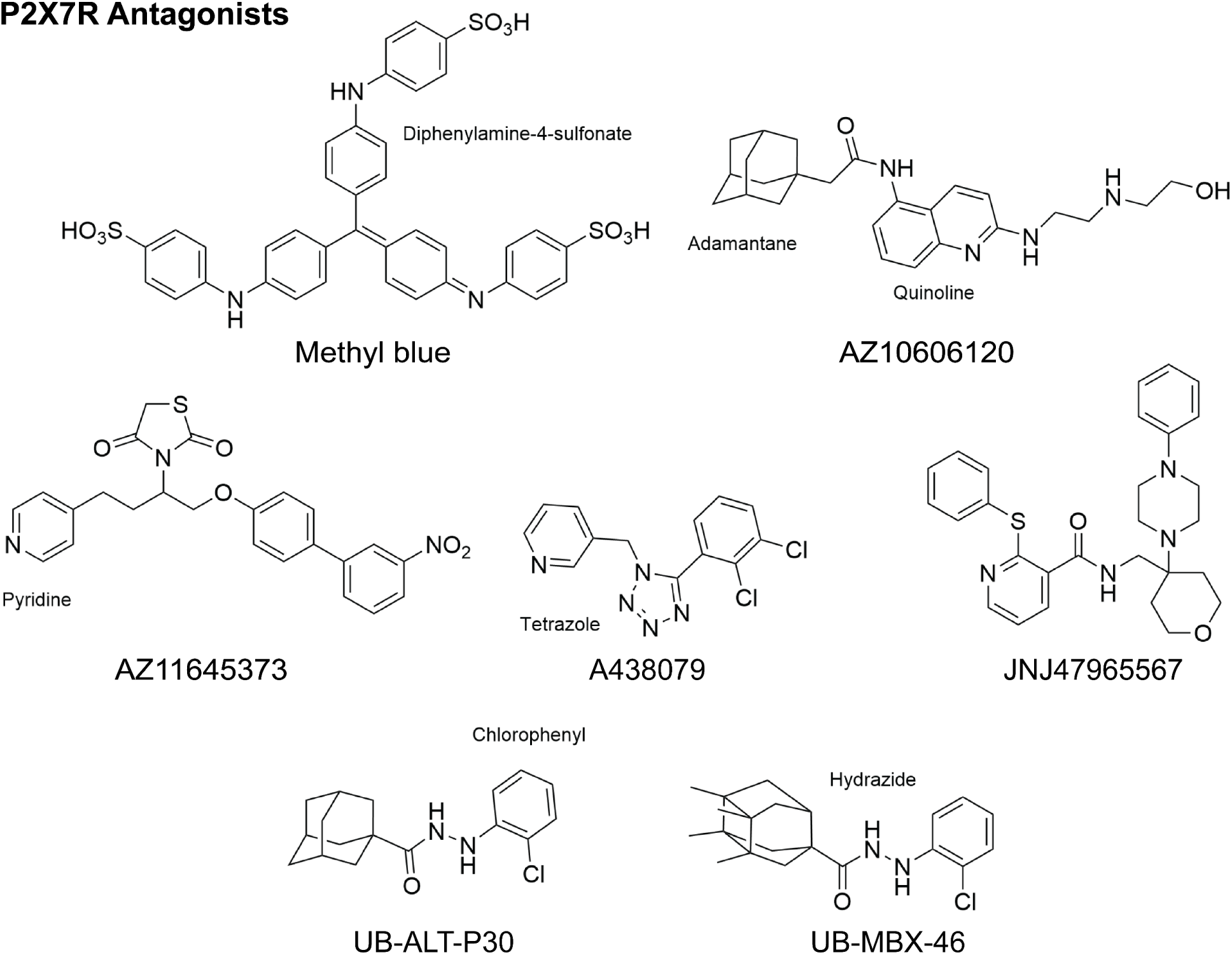
2D chemical structures of P2X7R antagonists. The structural basis of antagonism for AZ10606120, A438079, JNJ47965567, and methyl blue has been previously described^33,34^. The synthesis and activity of UB-ALT-P30 was originally disclosed by Abbott^40^. In contrast, UB-MBX-46 is a new ligand scaffold with high potency and selectivity.

**Extended Data Fig. 2:**
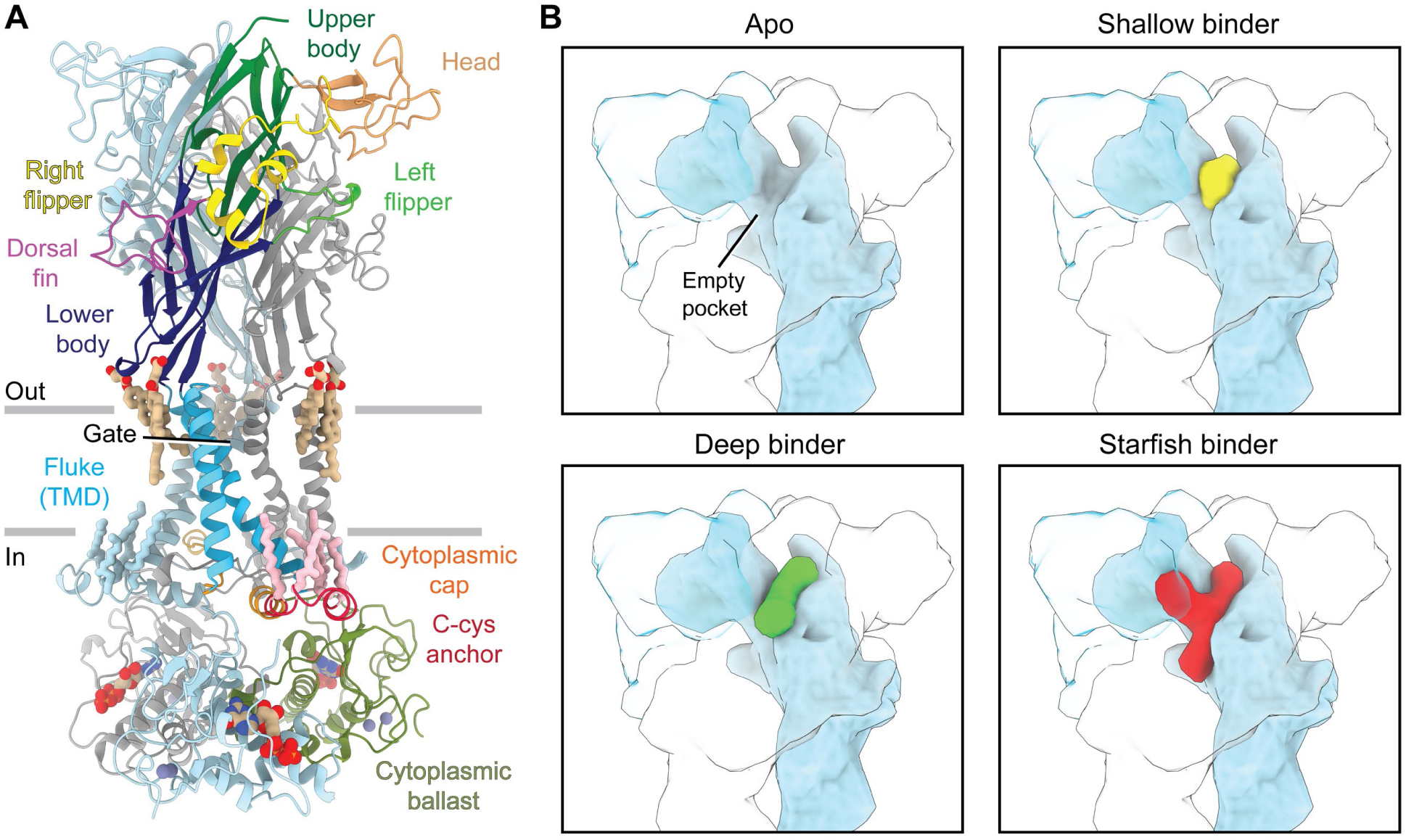
Domain architecture and allosteric binding modes of the P2X7R. (**A**) Ribbon representation of the hP2X7R in the apo closed state with one protomer colored by domain nomenclature and the other two colored in light blue and grey^36,71^. (**B**) Cartoon representation of the receptor highlighting three antagonist binding modes defined from structures of antagonists bound to the rP2X7R ortholog, with one protomer shown in light blue and the other two in transparent blue and transparent light grey^34^. The apo receptor with an empty pocket contrasts shallow binding ligands (yellow), deep binding ligands (green), and starfish binding ligands (red)^34^. Each mode of binding has distinct functional properties^34^.

**Extended Data Fig. 3:**
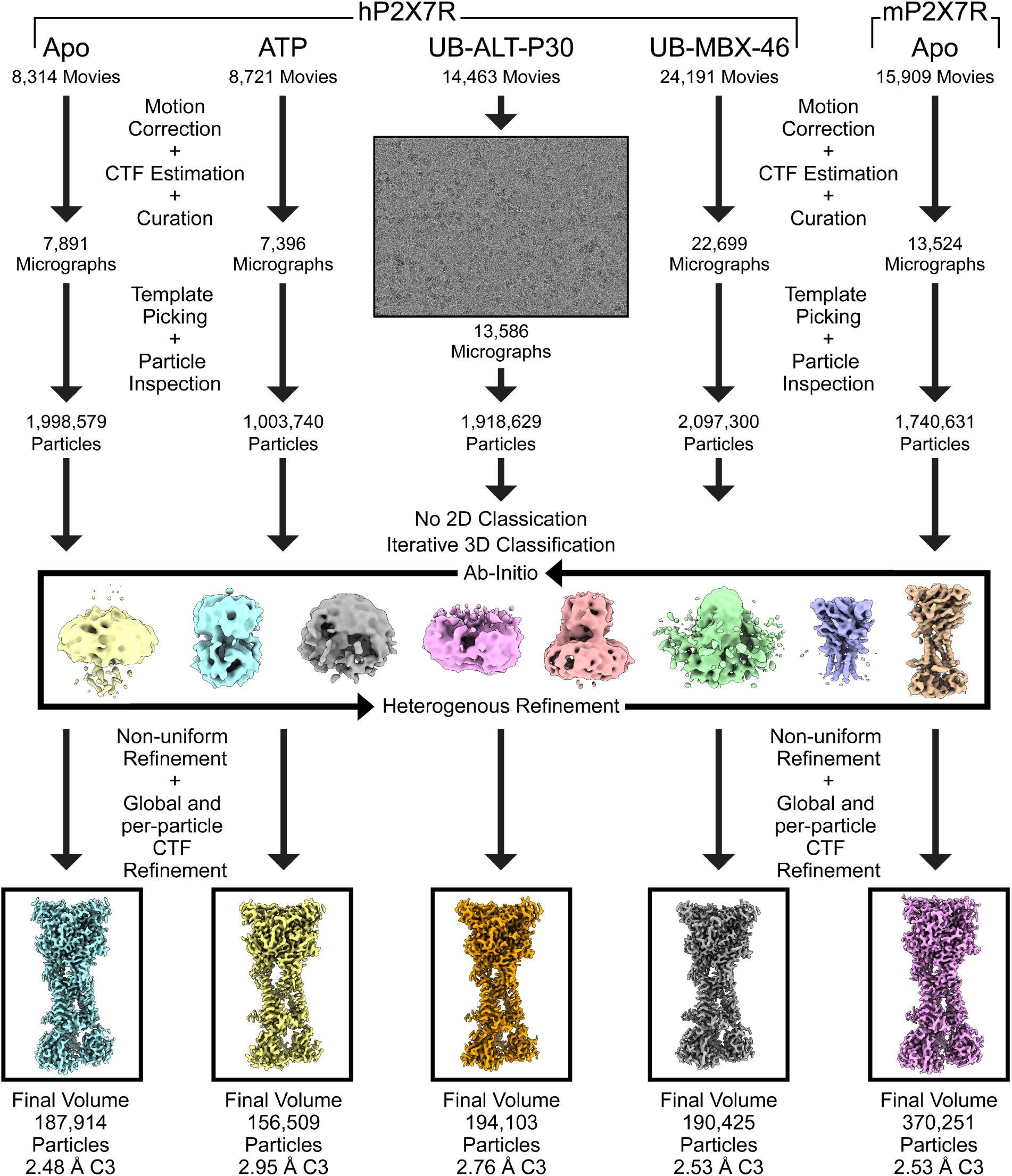
Cryo-EM processing pipeline for P2X7R reconstructions. Following data acquisition, movies were binned to the physical pixel size during motion correction and patch CTF correction was performed in cryoSPARC^58^. Next, micrographs were culled, and particles were picked using 2D templates generated from a low-resolution 3D reconstruction. Initial particle picks were inspected, extracted, and sent directly to iterative 3D classification in cryoSPARC using ab initio and heterogeneous refinement jobs. After classification into a culled particle stack, the final particles were re-extracted at the physical pixel size for non-uniform refinement performed with global and per-particle CTF refinements^59^.

**Extended Data Fig. 4:**
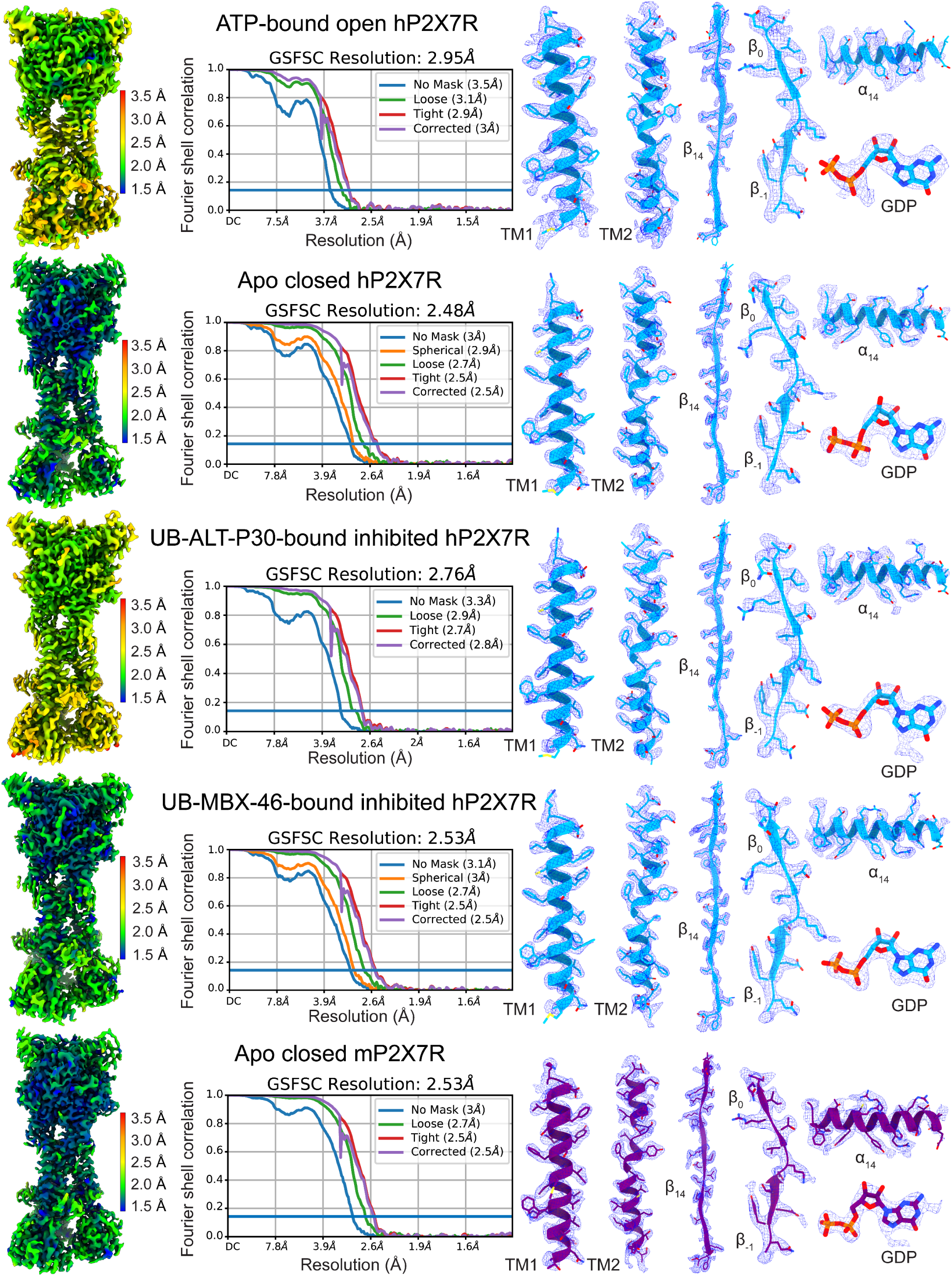
FSC and local resolution plots for P2X7R reconstructions. Resolution stated is at an FSC = 0.143. All local resolution plots range between 1.5 Å (blue) and 3.5 Å (red). Representative regions (TM1, TM2, β_-1_, β_0_, β_14_, α_14_, and GDP) from the ATP-bound open hP2X7R, apo closed hP2X7R, UB-ALT-P30-bound inhibited hP2X7R, UB-MBX-46-bound inhibited hP2X7R, and apo closed mP2X7R structures shown within their respective electron densities (blue mesh) highlighting the strong map-to-model fit. Models for the hP2X7R are colored blue and the model for the mP2X7R is colored purple.

**Extended Data Fig. 5:**
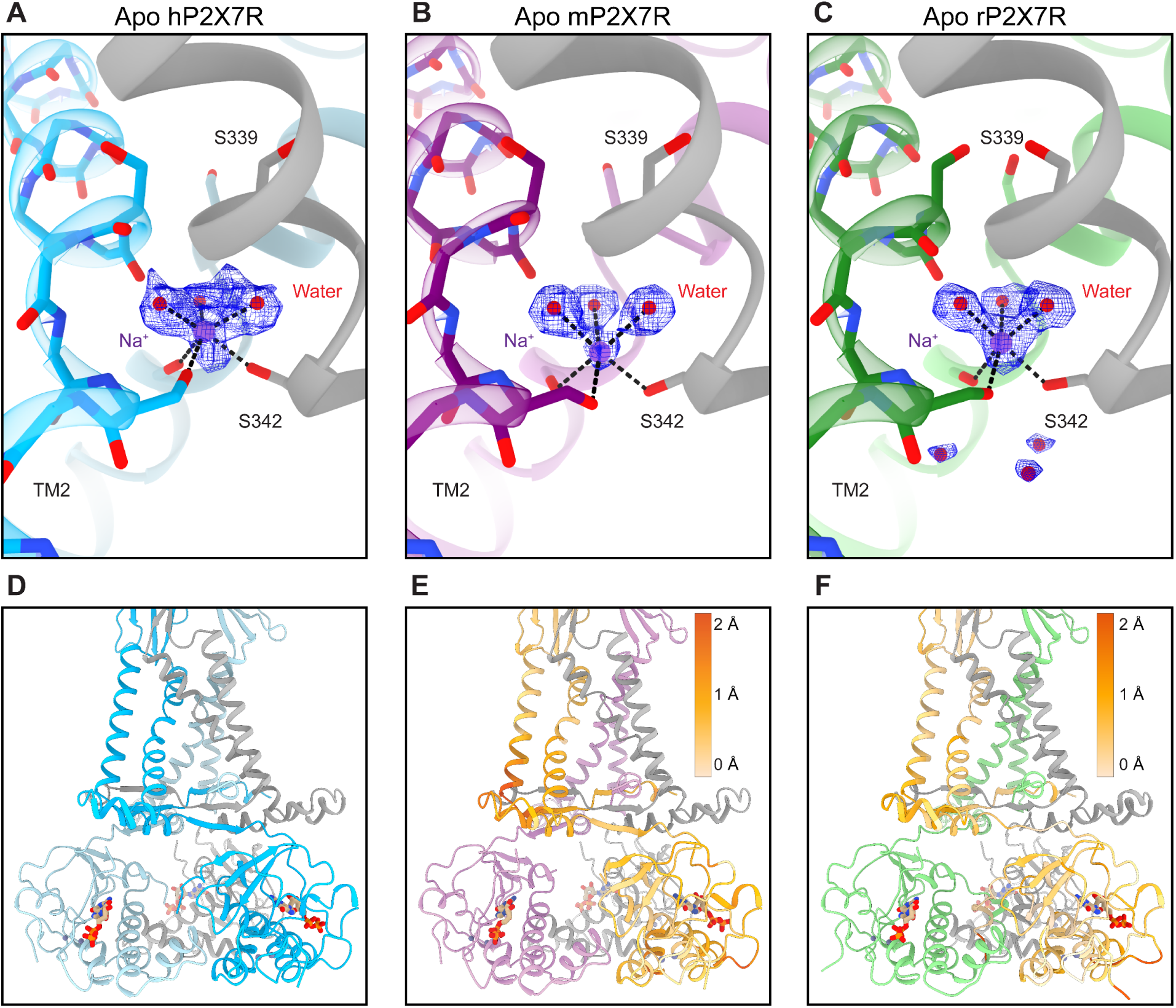
The pore architecture and cytoplasmic domain of P2X7Rs are conserved across human, mouse, and rat orthologs. (**A-C**) Ribbon representation of the hP2X7R (shades of blue and grey), the mP2X7R (shades of purple and grey), and the rP2X7R (shades of green and grey, PDB code 8TR5) in the apo closed state highlighting the coordination of a partially hydrated Na^+^ ion (purple sphere) directly above the closed gate with the electron density shown in blue mesh^35^. Across all three ortholog structures, the partially hydrated Na^+^ ion is coordinated by three symmetry-related waters above (red spheres) and the sidechain hydroxyl of S342 from each protomer below^35^. The identity of the partially hydrated Na^+^ ion was previously confirmed and the same coordination is now observed in human and mouse orthologs for the first time^34,35^. (**D-F**) Ribbon representation of the transmembrane and cytoplasmic domains of human, mouse, and rat P2X7Rs in the apo closed state are similar and maintain the same ligands (one GDP and two zinc ions per protomer). (**D-F**) View highlighting the cytoplasmic domains of the hP2X7R (shades of blue and grey), the mP2X7R (shades of purple and grey), and the rP2X7R (shades of green and grey, PDB code 8TR5)^35^. (**E and F**) Same view as **D** with one protomer of the mP2X7R (**E**) or the rP2X7R (**F**) colored in shades of orange by RMSD as compared to the human ortholog, highlighting the similarity between ortholog structures. A light tan color represents an RMSD of 0 Å while a dark orange indicates a 2 Å or greater change.

**Extended Data Fig. 6:**
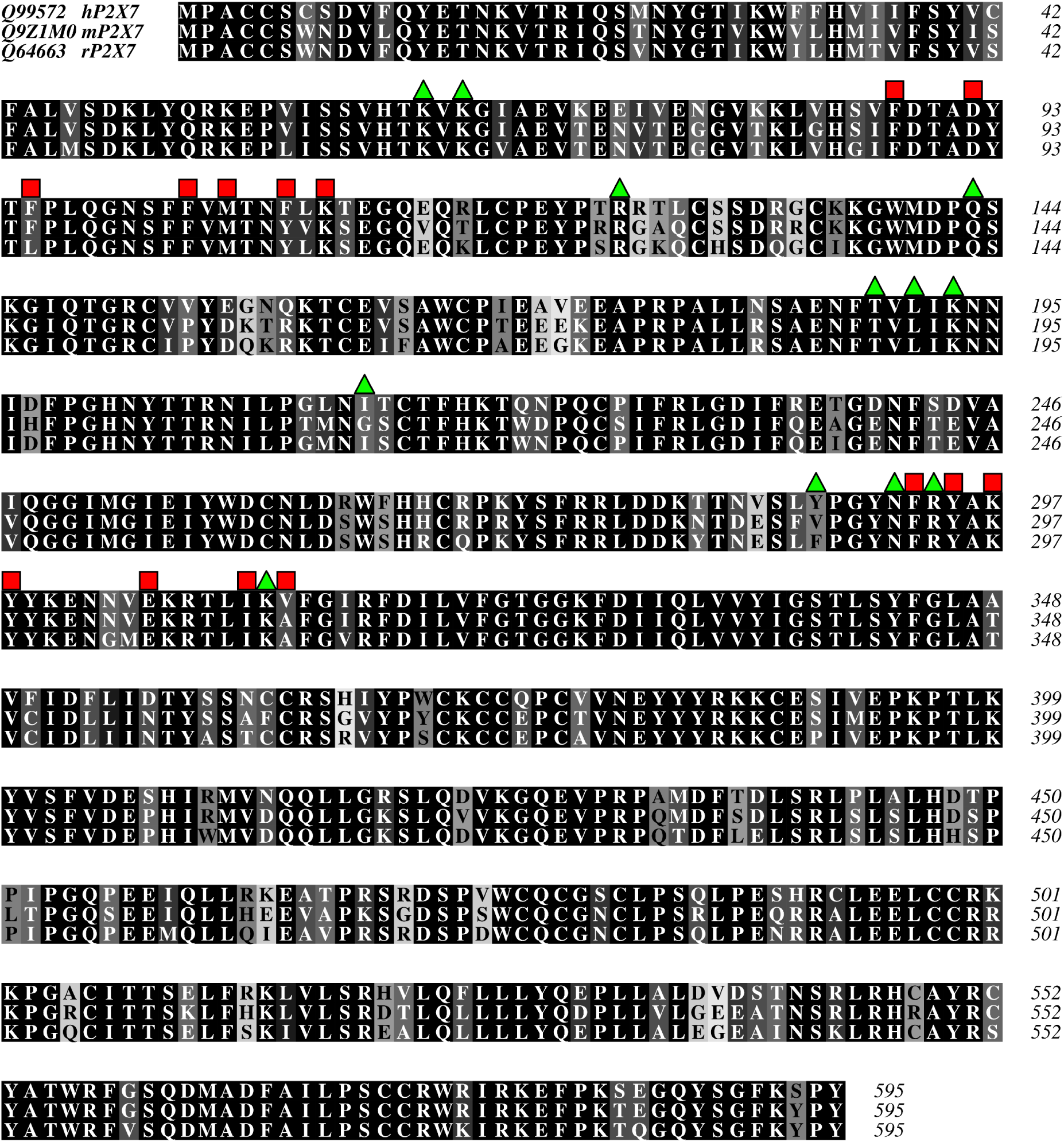
Sequence alignment of human, mouse, and rat P2X7Rs. Protein sequence alignment of human, mouse, and rat P2X7Rs colored by level of sequence conservation as calculated by Alscript with black denoting full sequence conservation and white denoting no sequence conservation^72^. Green triangles above the alignment indicate the residues that coordinate ATP. Red squares above the alignment indicate the residues that are located within the classical allosteric ligand-binding site. The sequence alignment was generated in Clustal Omega and visualized in Aline^73,74^. Sequences were obtained from UniProt with accession numbers Q99572, Q9Z1M0, and Q64663 for the hP2X7R, the mP2X7R, and the rP2X7R, respectively.

**Extended Data Fig. 7:**
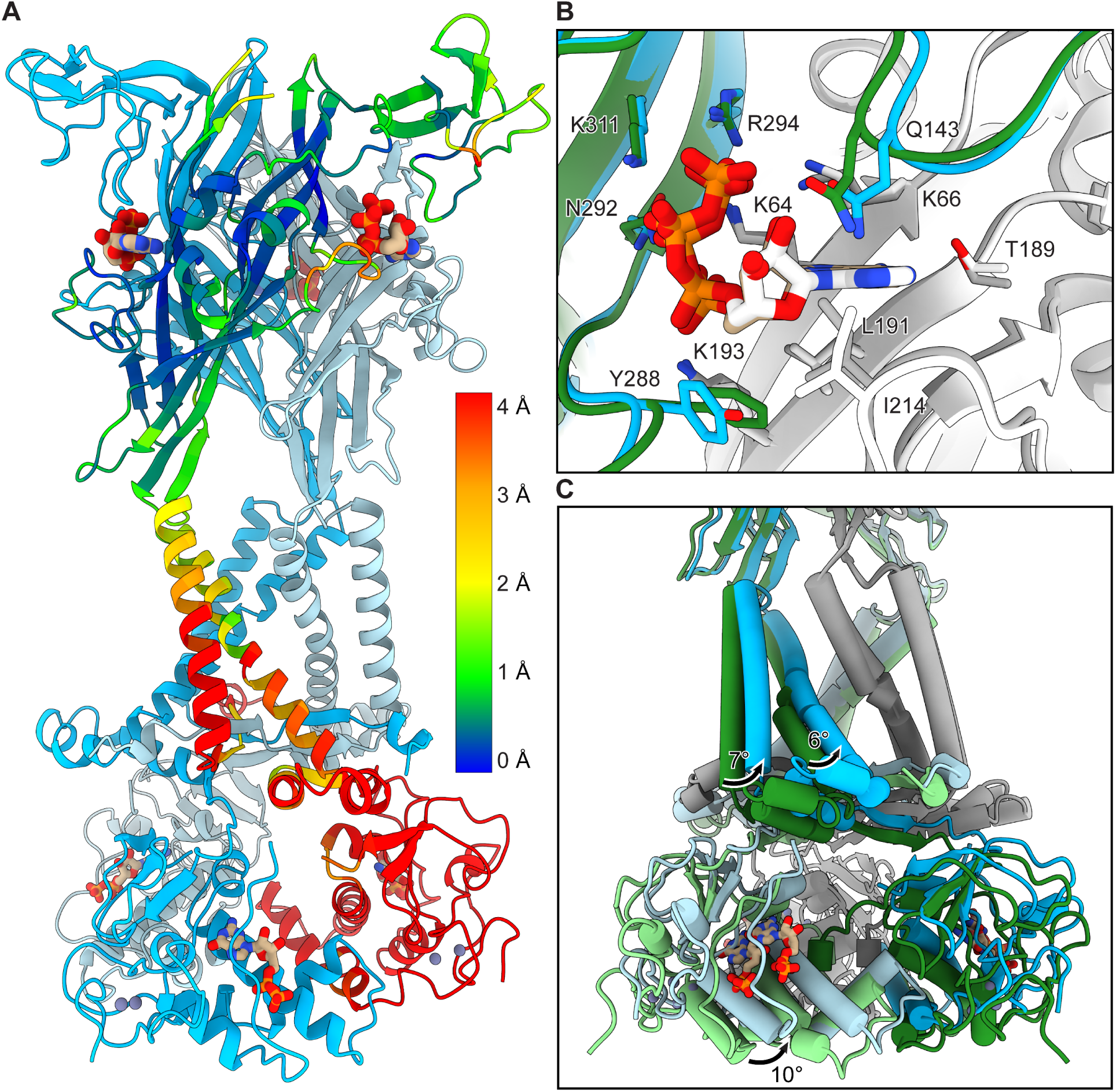
ATP-bound open state conformational differences between rat and human P2X7Rs. (**A**) Ribbon representation of the hP2X7R in the ATP-bound open state (shades of blue) with one protomer colored by RMSD according to the structural differences between human and rat ortholog structures (PDB code: 8TR5)^35^. Residues colored in blue signify little change with an RMSD of 0 Å while residues colored in red signify a larger change with an RMSD of >4 Å. Overall, the extracellular domains of ATP-bound P2X7R structures are similar with small RMSD’s while the transmembrane and cytoplasmic domains are different with larger RMSD’s. (**B**) Same view as Fig. 2E highlighting the structural differences between the ATP-bound open states of the rP2X7R (green and white with the carbon atoms of ATP in white) and the hP2X7R (blue and grey with the carbon atoms of ATP in tan). (**C**) Magnified view of the transmembrane and cytoplasmic domains of the rP2X7R and the hP2X7R in the ATP-bound open state highlighting 7° and 6° rotations of TM1 and TM2, respectively, ultimately resulting in a 10° rotation of the cytoplasmic ballast between rat and human receptor orthologs.

**Extended Data Fig. 8:**
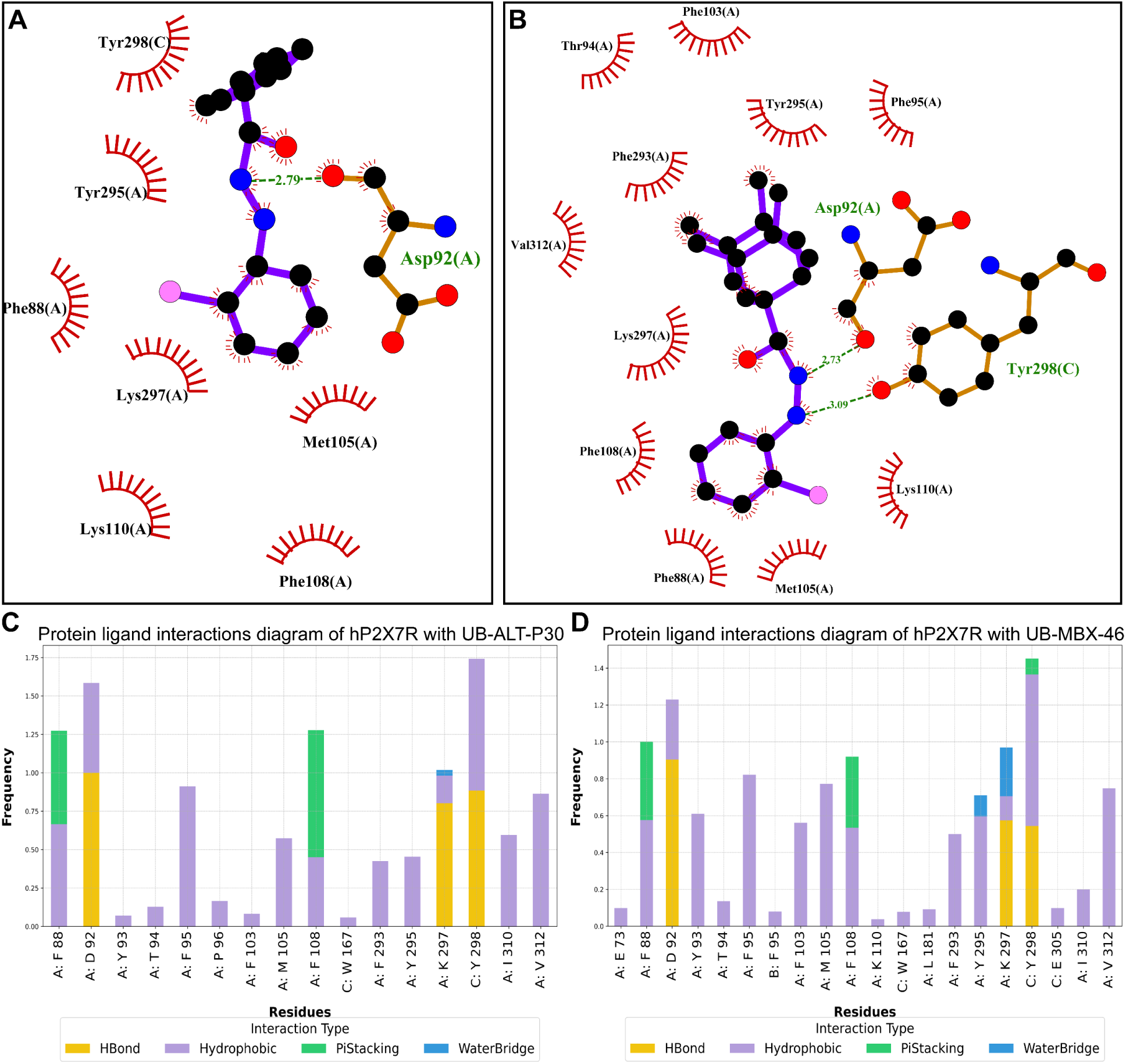
Receptor-ligand interactions for UB-ALT-P30 and UB-MBX-46 bound to the classical allosteric pocket of the hP2X7R. (**A and B**) LigPlot diagrams for UB-ALT-P30 (**A**) and UB-MBX-46 (**B**) bound to the classical allosteric pocket of the hP2X7R from cryo-EM structures^75^. These diagrams highlight the interactions between the ligand and the receptor. (**C and D**) Results from 500 ns or 1 μs MD simulations of f-hP2X7R in complex with UB-ALT-P30 (**C**) or UB-MBX-46 (**D**), respectively, embedded in POPC bilayers. Protein-ligand frequency interactions are depicted with bar plots. Each color within the graph corresponds to a different interaction: hydrogen bonding interactions are shown with yellow bars; hydrophobic interactions are shown with magenta bars; cation-π interactions are shown with green bars; water bridges with blue bars. Bars are plotted only for residues with interaction frequencies ≥ 0.2.

**Extended Data Fig. 9:**
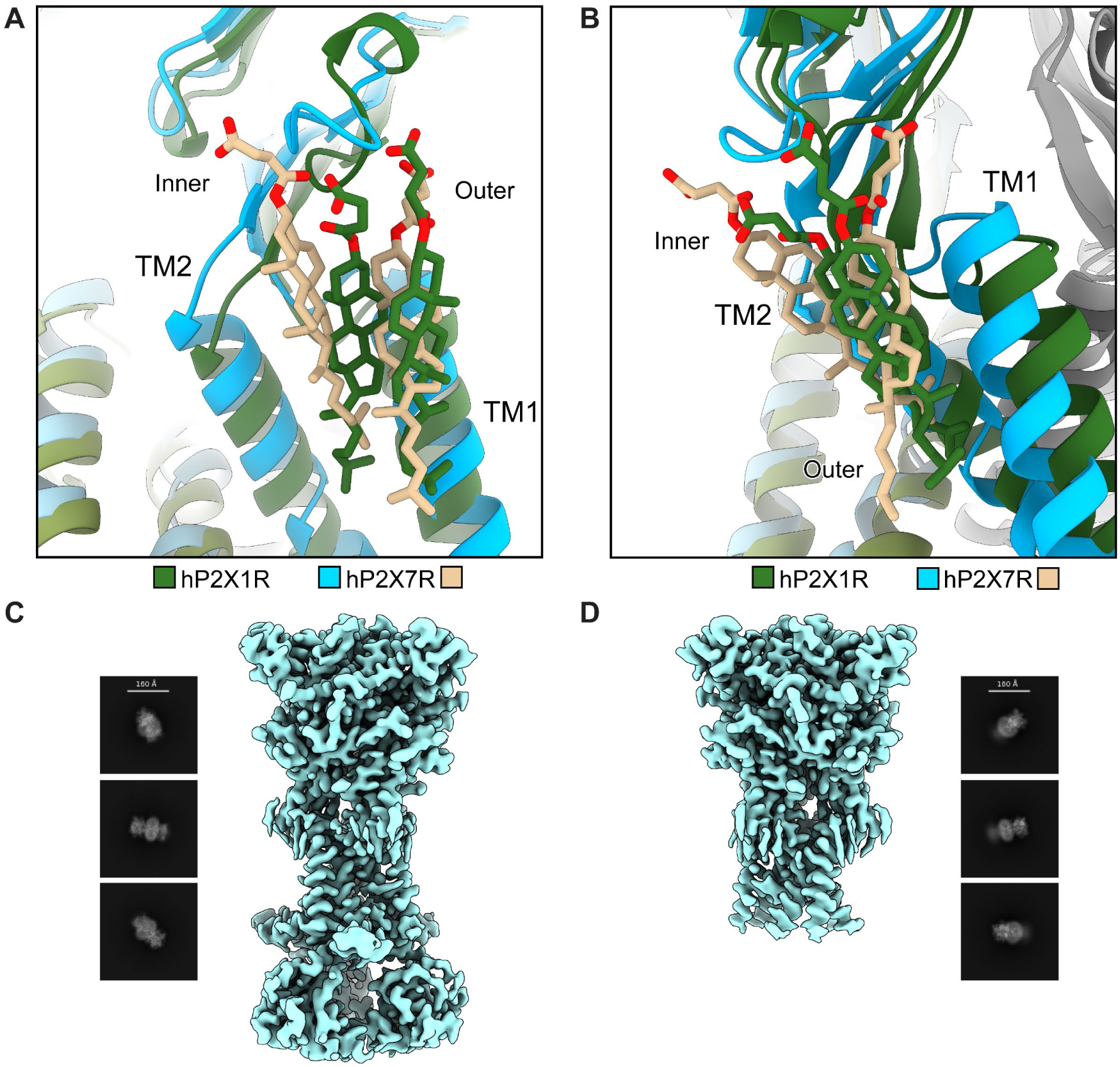
The cytoplasmic cap of the hP2X7R is stabilized by cholesterol. (**A and B**) Aligned structures of the hP2X1R and the hP2X7R highlighting the different positions of CHS molecules within their respective transmembrane domains. (**A**) Same view as Fig. 1C highlighting the position of the inner CHS molecules in the hP2X1R (receptor in green and CHS molecules in green) and the hP2X7R (receptor in blue and CHS molecules in tan). (**B**) Same view as Fig. 1D highlighting the position of the outer CHS molecules in the hP2X1R and the hP2X7R. Structures were aligned within ChimeraX^76^. (**C and D**) Cryo-EM processing of the hP2X7R in the apo closed and antagonist-bound inhibited states separated out reconstructions with (**C**) and without (**D**) the cytoplasmic domain which includes the cytoplasmic cap, C-cys anchor, and cytoplasmic ballast. 2D projections from each reconstruction are shown next to the map to highlight a lack of ordered density for the ballast-less reconstruction. Reconstructions with or without the cytoplasmic domain contain densities for CHS molecules.

**Extended Data Table 1:** Cryo-EM collection, refinement, and validation statistics.

|  | Apo<br>hP2X7R<br>(EMD-47490)<br>(PDB: 9E3M) | ATP<br>hP2X7R<br>(EMD-47491)<br>(PDB: 9E3N) | UB-ALT-P30<br>hP2X7R<br>(EMD-47492)<br>(PDB: 9E3O) | UB-MBX-46<br>hP2X7R<br>(EMDB-47493)<br>(PDB: 9E3P) | Apo<br>mP2X7R<br>(EMD-47494)<br>(PDB: 9E3Q) |
| --- | --- | --- | --- | --- | --- |
| **Data collection and processing** |  |  |  |  |  |
| Magnification (kx) | 130 | 37 | 130 | 130 | 130 |
| Voltage (kV) | 300 | 300 | 300 | 300 | 300 |
| Electron exposure (e-/Å) | 42 | 45 | 43 | 42 | 45 |
| Movie frames | 50 | 48 | 48 | 44 | 50 |
| Defocus range (µm) | -0.8 to -1.4 | -1.0 to -1.5 | -0.9 to -1.5 | -0.9 to -1.4 | -0.9 to -1.5 |
| Pixel size (Å/pixel) | 0.648 (0.324<br>super-res) | 0.623 (0.3115<br>super-res) | 0.6507 (0.32535<br>super-res) | 0.6483 (0.3242<br>super-res) | 0.647 (0.3235<br>super-res) |
| Symmetry imposed | C3 | C3 | C3 | C3 | C3 |
| Initial micrographs (no.) | 8,314 | 8,721 | 14,463 | 24,191 | 15,909 |
| Final micrographs used (no.) | 7,891 | 7,396 | 13,586 | 22,699 | 13,524 |
| Initial particle images (no.) | 1,998,579 | 1,003,740 | 1,918,629 | 2,097,300 | 1,740,631 |
| Final particle images (no.) | 187,914 | 156,509 | 194,103 | 190,425 | 370,251 |
| Map resolution (Å) | 2.48 | 2.95 | 2.76 | 2.53 | 2.53 |
| FSC threshold | (0.143) | (0.143) | (0.143) | (0.143) | (0.143) |
| Map resolution range (Å) | 1.4 to 7.0 | 1.8 to 36 | 1.4 to 43 | 2.2 to 37 | 1.5 to 8.1 |
| **Refinement** |  |  |  |  |  |
| Initial model used (PDB code) | 8TR5 | 6U9W | 8TR5 | 8TR5 | 8TR5 |
| Model resolution (Å) | 2.45 | 2.91 | 2.69 | 2.50 | 2.49 |
| FSC threshold | (0.143) | (0.143) | (0.143) | (0.143) | (0.143) |
| Map sharpening *B* factor (Å<sup>2</sup>) | 76.9 | 97.8 | 81.9 | 75.3 | 84.7 |
| **Model composition** |  |  |  |  |  |
| Non-hydrogen atoms | 14,153 | 13,616 | 14,013 | 14,353 | 13,622 |
| Protein Residues | 1,635 | 1,623 | 1,623 | 1,632 | 1,620 |
| Ligands | 34 | 27 | 30 | 40 | 16 |
| Waters | 121 | 68 | 177 | 273 | 193 |
| ***B* factors (Å<sup>2</sup>)** |  |  |  |  |  |
| Protein | 17.8/118/49.8 | 8.03/106/58.7 | 6.0/63.8/31.4 | 9.80/68.4/33.0 | 12.5/90.4/39.4 |
| Ligand | 30.0/113/41.2 | 27.6/131/65.2 | 14.2/72.6/24.1 | 13.4/74.4/39.7 | 23.4/88.8/35. |
| Nucleotide | 66.9/66.9/66.9 | 90.0/91.8/90.6 | 47.3/47.3/47.3 | 39.9/41.1/40.7 | 37.5/37.5/37.5 |
| Water | 15.5/64.1/26.5 | 17.4/51.1/35/7 | 5.5/46.8/14.7 | 8.44/33.8/19.2 | 11.6/44.2/23.0 |
| **R.m.s. deviations** |  |  |  |  |  |
| Bond lengths (Å) | 0.008 (0) | 0.007 (0) | 0.009 (0) | 0.007 (0) | 0.004 (0) |
| Bond angles (°) | 0.762 (0) | 0.921 (0) | 0.746 (0) | 0.782 (0) | 0.600 (0) |
| **Validation** |  |  |  |  |  |
| MolProbity score | 1.14 | 1.26 | 1.22 | 1.05 | 1.27 |
| Clash score | 3.50 | 4.90 | 4.37 | 2.66 | 5.09 |
| Poor rotamers (%) | 0.00 | 0 | 0 | 0.00 | 0.00 |
| **Ramachandran plot** |  |  |  |  |  |
| Favored (%) | 98.29 | 98.09 | 98.27 | 98.85 | 98.85 |
| Allowed (%) | 1.71 | 1.91 | 1.73 | 1.15 | 1.15 |
| Disallowed (%) | 0 | 0 | 0 | 0 | 0 |

## References

1. Di Virgilio, F., Vultaggio-Poma, V., Falzoni, S. & Giuliani, A.L. Extracellular ATP: A powerful inflammatory mediator in the central nervous system. Neuropharmacology 224, 109333 (2023).

2. Valera, S. et al. A new class of ligand-gated ion channel defined by P2x receptor for extracellular ATP. Nature 371, 516–9 (1994).

3. Surprenant, A., Rassendren, F., Kawashima, E., North, R.A. & Buell, G. The cytolytic P2Z receptor for extracellular ATP identified as a P2X receptor (P2X7). Science 272, 735–8 (1996).

4. North, R.A. Molecular physiology of P2X receptors. Physiological Reviews 82, 1013–67 (2002).

5. Illes, P. et al. Update of P2X receptor properties and their pharmacology: IUPHAR Review 30. British Journal of Pharmacology 178, 489–514 (2021).

6. Cheewatrakoolpong, B., Gilchrest, H., Anthes, J.C. & Greenfeder, S. Identification and characterization of splice variants of the human P2X7 ATP channel. Biochemical and Biophysical Research Communications 332, 17–27 (2005).

7. Kopp, R., Krautloher, A., Ramirez-Fernandez, A. & Nicke, A. P2X7 interactions and signaling - making head or tail of It. Frontiers in Molecular Neuroscience 12, 183 (2019).

8. Costa-Junior, H.M., Sarmento Vieira, F. & Coutinho-Silva, R. C terminus of the P2X7 receptor: treasure hunting. Purinergic Signal 7, 7–19 (2011).

9. Pelegrin, P. P2X7 receptor and the NLRP3 inflammasome: Partners in crime. Biochemical Pharmacology 187, 114385 (2021).

10. Kong, H., Zhao, H., Chen, T., Song, Y. & Cui, Y. Targeted P2X7/NLRP3 signaling pathway against inflammation, apoptosis, and pyroptosis of retinal endothelial cells in diabetic retinopathy. Cell Death & Disease 13, 336 (2022).

11. Franceschini, A. et al. The P2X7 receptor directly interacts with the NLRP3 inflammasome scaffold protein. FASEB Journal 29, 2450–61 (2015).

12. Wang, W., Xiao, J., Adachi, M., Liu, Z. & Zhou, J. 4-aminopyridine induces apoptosis of human acute myeloid leukemia cells via increasing [Ca2+]i through P2X7 receptor pathway. Cellular Physiology and Biochemistry 28, 199–208 (2011).

13. Burnstock, G. & Kennedy, C. P2X receptors in health and disease. Advances in Pharmacology 61, 333–72 (2011).

14. Burnstock, G. & Knight, G.E. The potential of P2X7 receptors as a therapeutic target, including inflammation and tumour progression. Purinergic Signal 14, 1–18 (2018).

15. Furlan-Freguia, C., Marchese, P., Gruber, A., Ruggeri, Z.M. & Ruf, W. P2X7 receptor signaling contributes to tissue factor-dependent thrombosis in mice. The Journal of Clinical Investigation 121, 2932–44 (2011).

16. Francistiova, L. et al. The Role of P2X7 Receptor in Alzheimer’s Disease. Frontiers in Molecular Neuroscience 13, 94 (2020).

17. Pegoraro, A., De Marchi, E. & Adinolfi, E. P2X7 Variants in Oncogenesis. Cells 10(2021).

18. Chen, X. et al. Brilliant Blue G improves cognition in an animal model of Alzheimer’s disease and inhibits amyloid-β-induced loss of filopodia and dendrite spines in hippocampal neurons. Neuroscience 279, 94–101 (2014).

19. Lara, R. et al. P2X7 in cancer: from molecular mechanisms to therapeutics. Frontiers in Pharmacology 11, 793 (2020).

20. Shokoples, B.G., Paradis, P. & Schiffrin, E.L. P2X7 receptors: an untapped target for the management of cardiovascular disease. Arteriosclerosis, Thrombosis, and Vascular Biology 41, 186–199 (2021).

21. Stachon, P. et al. P2X7 deficiency blocks lesional inflammasome activity and ameliorates atherosclerosis in mice. Circulation 135, 2524–2533 (2017).

22. Zhang, R. et al. From lead to clinic: A review of the structural design of P2X7R antagonists. European Journal of Medicinal Chemistry 251, 115234 (2023).

23. Mafra, J.C.M., Boechat, N., Teixeira, G.P. & Faria, R.X. Synthetic molecules as P2X7 receptor antagonists: A medicinal chemistry update focusing the therapy of inflammatory diseases. European Journal of Pharmacology 957, 175999 (2023).

24. Muller, C.E. & Namasivayam, V. Recommended tool compounds and drugs for blocking P2X and P2Y receptors. Purinergic Signal 17, 633–648 (2021).

25. Liu, X. et al. Unlocking the therapeutic potential of P2X7 receptor: a comprehensive review of its role in neurodegenerative disorders. Frontiers in Pharmacology 15(2024).

26. Oken, A.C. et al. Molecular Pharmacology of P2X Receptors: Exploring Druggable Domains Revealed by Structural Biology. Frontiers in Pharmacology 13(2022).

27. Bhattacharya, A. et al. Pharmacological characterization of a novel centrally permeable P2X7 receptor antagonist: JNJ-47965567. British Journal of Pharmacology 170, 624–640 (2013).

28. Stokes, L. et al. Characterization of a selective and potent antagonist of human P2X(7) receptors, AZ11645373. British Journal of Pharmacology 149, 880–7 (2006).

29. Michel, A.D. et al. Mechanism of action of species-selective P2X(7) receptor antagonists. British Journal of Pharmacology 156, 1312–25 (2009).

30. Donnelly-Roberts, D.L., Namovic, M.T., Han, P. & Jarvis, M.F. Mammalian P2X7 receptor pharmacology: comparison of recombinant mouse, rat and human P2X7 receptors. Br J Pharmacol 157, 1203–14 (2009).

31. Lionta, E., Spyrou, G., Vassilatis, D.K. & Cournia, Z. Structure-based virtual screening for drug discovery: principles, applications and recent advances. Curr Top Med Chem 14, 1923–38 (2014).

32. Lees, J.A., Dias, J.M. & Han, S. Applications of Cryo-EM in small molecule and biologics drug design. Biochemical Society Transactions 49, 2627–2638 (2021).

33. Karasawa, A. & Kawate, T. Structural basis for subtype-specific inhibition of the P2X7 receptor. Elife 5(2016).

34. Oken, A.C. et al. P2X7 receptors exhibit at least three modes of allosteric antagonism. Science Advances 10, eado5084 (2024).

35. Oken, A.C. et al. High-affinity agonism at the P2X7 receptor is mediated by three residues outside the orthosteric pocket. Nature Communications 15, 6662 (2024).

36. McCarthy, A.E., Yoshioka, C. & Mansoor, S.E. Full-Length P2X(7) Structures Reveal How Palmitoylation Prevents Channel Desensitization. Cell 179, 659–670 e13 (2019).

37. Roger, S. et al. Understanding the roles of the P2X7 receptor in solid tumour progression and therapeutic perspectives. Biochimica et Biophysica Acta (BBA) - Biomembranes 1848, 2584–2602 (2015).

38. Chessell, I.P. et al. Cloning and functional characterisation of the mouse P2X7 receptor. FEBS Letters 439, 26–30 (1998).

39. Degrève, L., Vechi, S.M. & Junior, C.Q. The hydration structure of the Na+ and K+ ions and the selectivity of their ionic channels. Biochimica et Biophysica Acta (BBA) - Bioenergetics 1274, 149–156 (1996).

40. Nelson, D.W. et al. Structure−Activity Relationship Studies on N′-Aryl Carbohydrazide P2X7 Antagonists. Journal of Medicinal Chemistry 51, 3030–3034 (2008).

41. Wagner, J.R. et al. POVME 3.0: Software for Mapping Binding Pocket Flexibility. Journal of Chemical Theory and Computation 13, 4584–4592 (2017).

42. Wanka, L., Iqbal, K. & Schreiner, P.R. The lipophilic bullet hits the targets: medicinal chemistry of adamantane derivatives. Chemical Reviews 113, 3516–3604 (2013).

43. Rey-Carrizo, M. et al. Easily accessible polycyclic amines that inhibit the wild-type and amantadine-resistant mutants of the M2 channel of influenza A virus. Journal of Medicinal Chemistry 57, 5738–5747 (2014).

44. Codony, S., et al. Synthesis, In Vitro Profiling, and In Vivo Evaluation of Benzohomoadamantane-Based Ureas for Visceral Pain: A New Indication for Soluble Epoxide Hydrolase Inhibitors. Journal of Medicinal Chemistry 65, 13660–13680 (2022).

45. Barniol-Xicota, M. et al. Escape from adamantane: Scaffold optimization of novel P2X7 antagonists featuring complex polycycles. Bioorganic & Medicinal Chemistry Letters 27, 759–763 (2017).

46. Leiva, R. et al. Pharmacological and Electrophysiological Characterization of Novel NMDA Receptor Antagonists. ACS Chemical Neuroscience 9, 2722–2730 (2018).

47. Song, L.F., Lee, T.-S., Zhu, C., York, D.M. & Merz, K.M., Jr. Using AMBER18 for relative free energy calculations. Journal of Chemical Information and Modeling 59, 3128–3135 (2019).

48. He, X. et al. Fast, accurate, and reliable protocols for routine calculations of protein–ligand binding affinities in drug design projects using AMBER GPU-TI with ff14SB/GAFF. ACS Omega 5, 4611–4619 (2020).

49. Case, D.A. et al. AmberTools. Journal of Chemical Information and Modeling 63, 6183–6191 (2023).

50. Tian, C. et al. ff19SB: Amino-Acid-Specific Protein Backbone Parameters Trained against Quantum Mechanics Energy Surfaces in Solution. Journal of Chemical Theory and Computation 16, 528–552 (2020).

51. Oken, A.C. et al. Cryo-EM structures of the human P2X1 receptor reveal subtype-specific architecture and antagonism by supramolecular ligand-binding. Nature Communications 15, 8490 (2024).

52. Robinson, L.E., Shridar, M., Smith, P. & Murrell-Lagnado, R.D. Plasma membrane cholesterol as a regulator of human and rodent P2X7 receptor activation and sensitization. Journal of Biological Chemistry 289, 31983–31994 (2014).

53. Lordén, G. et al. Lipin-2 regulates NLRP3 inflammasome by affecting P2X7 receptor activation. Journal of Experimental Medicine 214, 511–528 (2016).

54. Bin Dayel, A., Evans, R.J. & Schmid, R. Mapping the site of action of human P2X7 receptor antagonists AZ11645373, Brilliant Blue G, KN-62, Calmidazolium, and ZINC58368839 to the intersubunit allosteric pocket. Molecular Pharmacology 96, 355 (2019).

55. Jiang, L.-H., Caseley, E.A., Muench, S.P. & Roger, S. Structural basis for the functional properties of the P2X7 receptor for extracellular ATP. Purinergic Signalling 17, 331–344 (2021).

56. Tsien, J., Hu, C., Merchant, R.R. & Qin, T. Three-dimensional saturated C(sp3)-rich bioisosteres for benzene. Nature Reviews Chemistry 8, 605–627 (2024).

57. Mastronarde, D.N. Automated electron microscope tomography using robust prediction of specimen movements. Journal of Structural Biology 152, 36–51 (2005).

58. Punjani, A., Rubinstein, J.L., Fleet, D.J. & Brubaker, M.A. cryoSPARC: algorithms for rapid unsupervised cryo-EM structure determination. Nature Methods 14, 290–296 (2017).

59. Punjani, A., Zhang, H. & Fleet, D.J. Non-uniform refinement: adaptive regularization improves single-particle cryo-EM reconstruction. Nature Methods 17, 1214–1221 (2020).

60. Waterhouse, A. et al. SWISS-MODEL: homology modelling of protein structures and complexes. Nucleic Acids Research 46, W296–W303 (2018).

61. Emsley, P., Lohkamp B Fau - Scott, W.G., Scott Wg Fau - Cowtan, K. & Cowtan, K. Features and development of Coot. Acta Crystallographica. Section D, Biological Crystallography 66(Pt 4), 486–501 (2010).

62. Liebschner, D. et al. Macromolecular structure determination using X-rays, neutrons and electrons: recent developments in Phenix. Acta Crystallographica Section D 75, 861–877 (2019).

63. Moriarty, N.W., Grosse-Kunstleve, R.W. & Adams, P.D. electronic Ligand Builder and Optimization Workbench (eLBOW): a tool for ligand coordinate and restraint generation. Acta Crystallographica Section D 65(Pt 10), 1074–1080 (2009).

64. Williams, C.J. et al. MolProbity: More and better reference data for improved all-atom structure validation. Protein Science 27, 293–315 (2018).

65. Weinhausen, S. et al. Extracellular binding sites of positive and negative allosteric P2X4 receptor modulators. Life Sciences 311, 121143 (2022).

66. Abdelrahman, A. et al. Characterization of P2X4 receptor agonists and antagonists by calcium influx and radioligand binding studies. Biochemical Pharmacology 125, 41–54 (2017).

67. Markowitz, D., Hesdorffer, C., Ward, M., Goff, S. & Bank, A. Retroviral Gene Transfer Using Safe and Efficient Packaging Cell Lines. Annals of the New York Academy of Sciences 612, 407–414 (1990).

68. Lee, S.-Y. et al. Establishment of an Assay for P2X7 Receptor-Mediated Cell Death. Molecules and Cells 22, 198–202 (2006).

69. Shirts, M.R. & Pande, V.S. Erratum: “Comparison of efficiency and bias of free energies computed by exponential averaging, the Bennett acceptance ratio, and thermodynamic integration” [J. Chem. Phys. 122, 144107 (2005)]. The Journal of Chemical Physics 129, 229901 (2008).

70. Le Guilloux, V., Schmidtke, P. & Tuffery, P. Fpocket: An open source platform for ligand pocket detection. BMC Bioinformatics 10, 168 (2009).

71. Kawate, T., Michel, J.C., Birdsong, W.T. & Gouaux, E. Crystal structure of the ATP-gated P2X(4) ion channel in the closed state. Nature 460, 592–8 (2009).

72. Barton, G.J. ALSCRIPT: a tool to format multiple sequence alignments. Protein Engineering, Design and Selection 6, 37–40 (1993).

73. Sievers, F. & Higgins, D.G. Clustal Omega for making accurate alignments of many protein sequences. Protein Science 27, 135–145 (2018).

74. Bond, C.S. & Schuttelkopf, A.W. ALINE: a WYSIWYG protein-sequence alignment editor for publication-quality alignments. Acta Crystallographica Section D, Biological Crystallography 65, 510–2 (2009).

75. Laskowski, R.A. & Swindells, M.B. LigPlot+: Multiple Ligand–Protein Interaction Diagrams for Drug Discovery. Journal of Chemical Information and Modeling 51, 2778–2786 (2011).

76. Meng, E.C. et al. UCSF ChimeraX: Tools for structure building and analysis. Protein Science 32, e4792 (2023).

