## Supplementary information for "UB-MBX-46 is a potent and selective antagonist of the human P2X7 receptor developed by structure-based drug design"

a. Department of Chemical Physiology & Biochemistry, Oregon Health & Science University, Portland, Oregon 97239, USA.
b. Laboratori de Química Farmacèutica, Facultat de Farmàcia i Ciències de l'Alimentació, Universitat de Barcelona, Av. Joan XXIII, 27-31, 08028 Barcelona, Spain.
c. Institute of Biomedicine of the University of Barcelona, IBUB, 08028 Barcelona, Spain.
d. Laboratory of Medicinal Chemistry, Section of Pharmaceutical Chemistry, Department of Pharmacy, National and Kapodistrian University of Athens, Panepistimiopolis-Zografou, 15771, Greece.
e. PharmaCenter Bonn & Pharmaceutical Institute, Pharmaceutical & Medicinal Chemistry, University of Bonn, 53121 Bonn, Germany.
f. Present address: Department of Medicine and Life Sciences, Biomedical Research Park (PRBB), Universitat Pompeu Fabra, 08003 Barcelona, Spain
g. School of Life Sciences, Gwangju Institute of Science and Technology, 123 Cheomdangwagi-ro, Buk-gu, Gwangju 61005, Republic of Korea.
h. Walther Straub Institute of Pharmacology and Toxicology, Faculty of Medicine, Ludwig-Maximilians-Universität München, Munich, Germany
i. Division of Cardiovascular Medicine, Knight Cardiovascular Institute, Oregon Health & Science University, Portland, Oregon 97239, USA.


**This PDF file includes:**

Supplementary Figures 1 to 6
Synthesis and characterization methods
Detailed molecular dynamics simulations methods

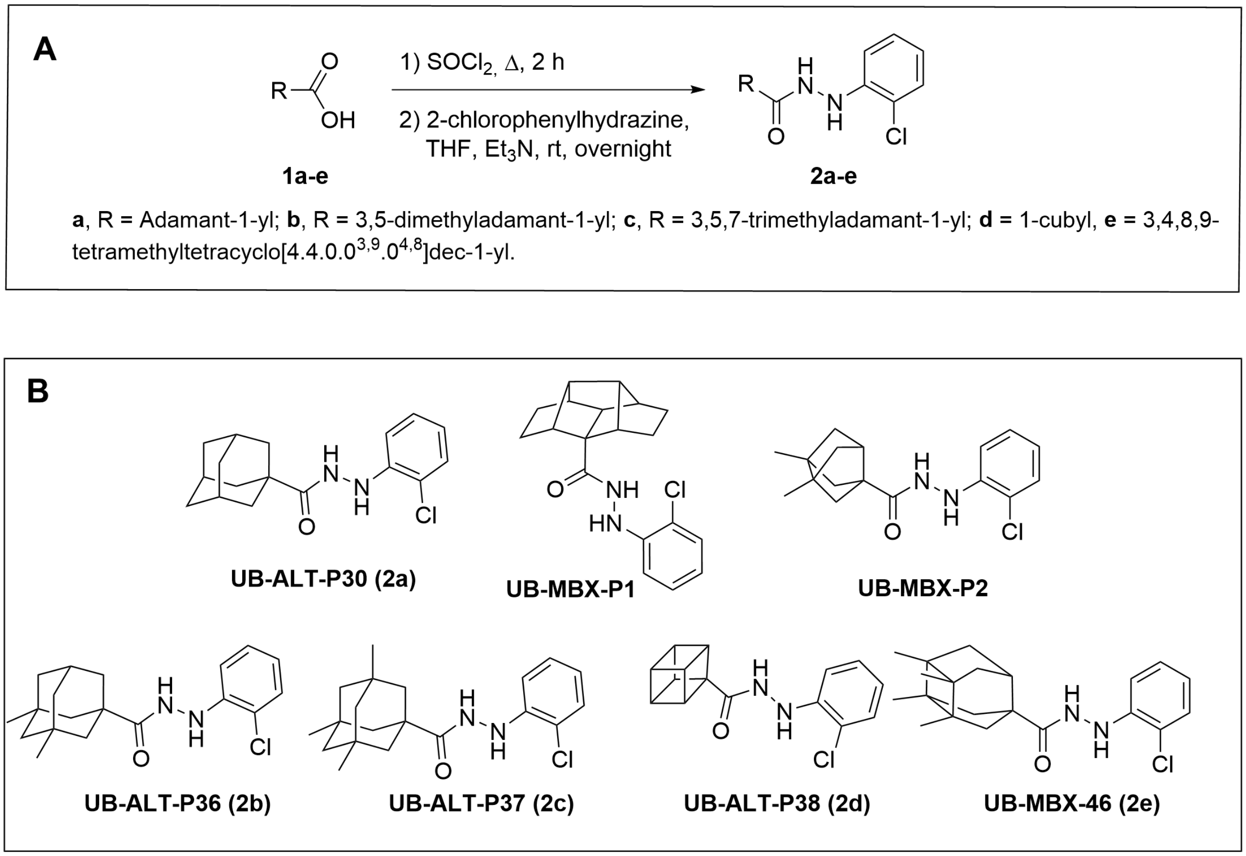


**Supplementary Figure 1:** (A) Synthetic route for the new acyl hydrazines. (B) Chemical structures of known P2X7R antagonists (UB-ALT-P30, UB-MBX-P1, UB-MBX-P2) and previously uncharacterized polycyclic analogs (UB-ALT-P36, UB-ALT-P37, UB-ALT-P38 and UB-MBX-46)[1,2].

44 [1]H NMR (400 MHz, $CDCl_3$)

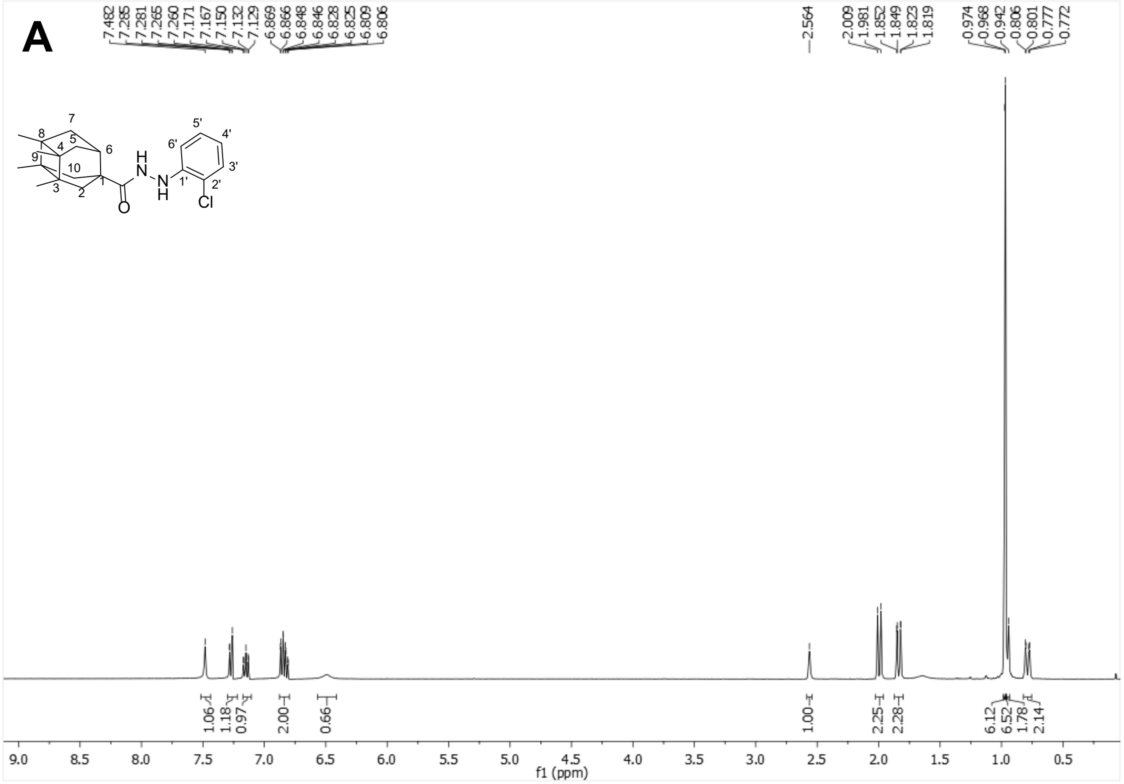


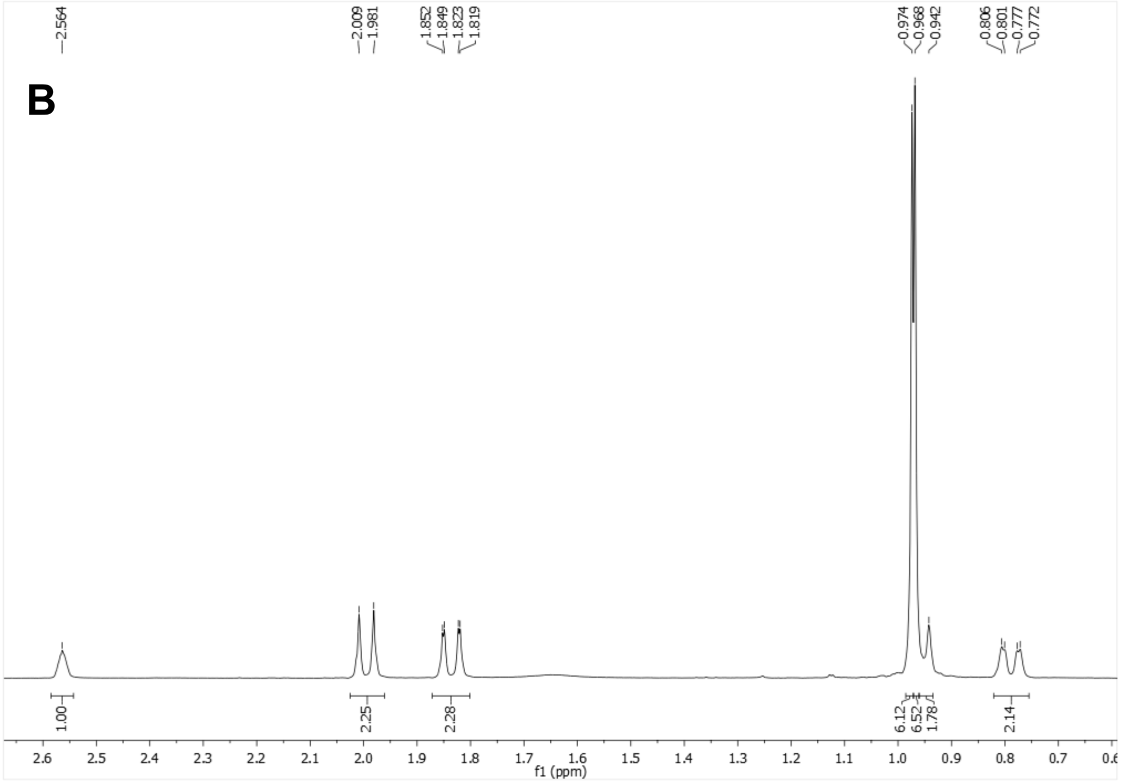

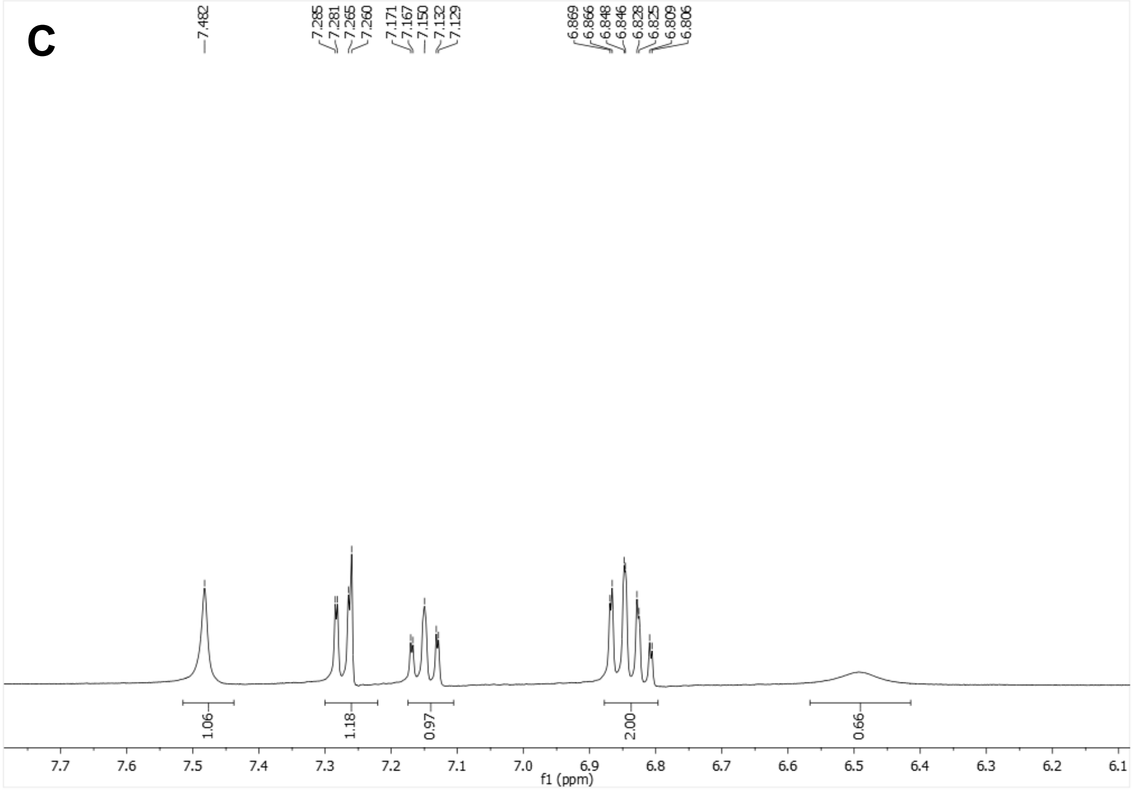


$^{13}$C-NMR (100.6 MHz, $CDCl_3$)

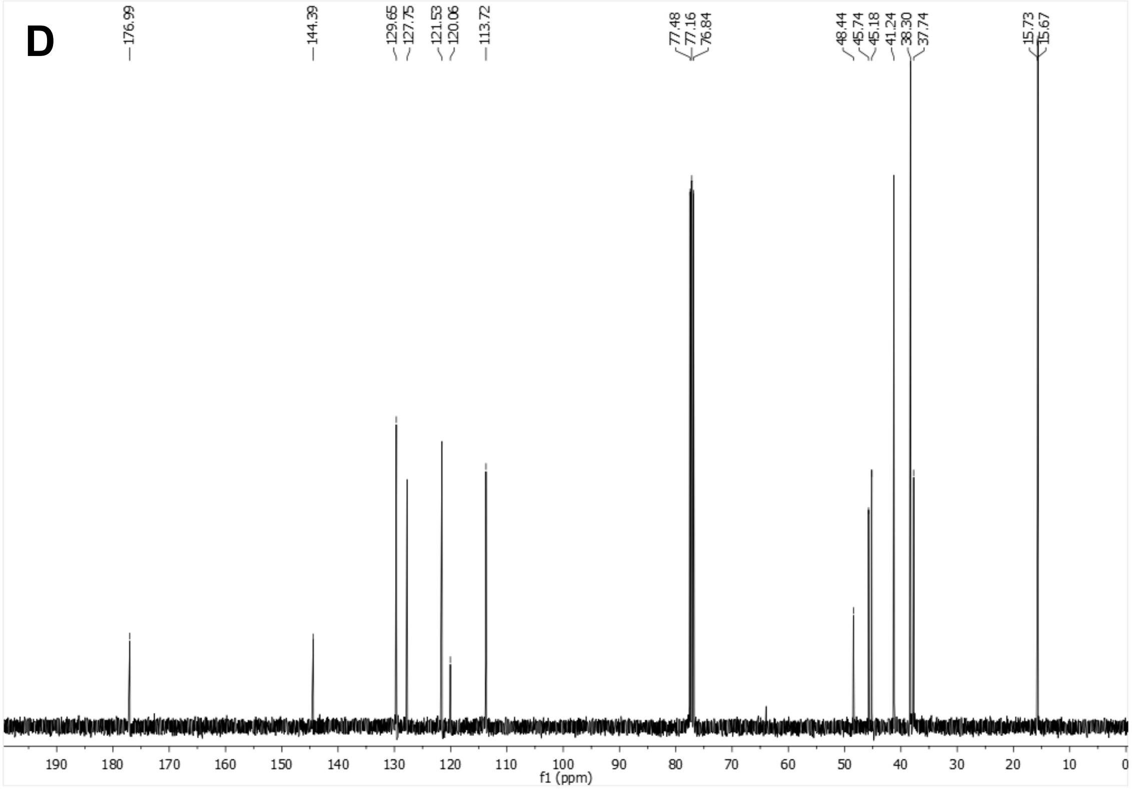

E

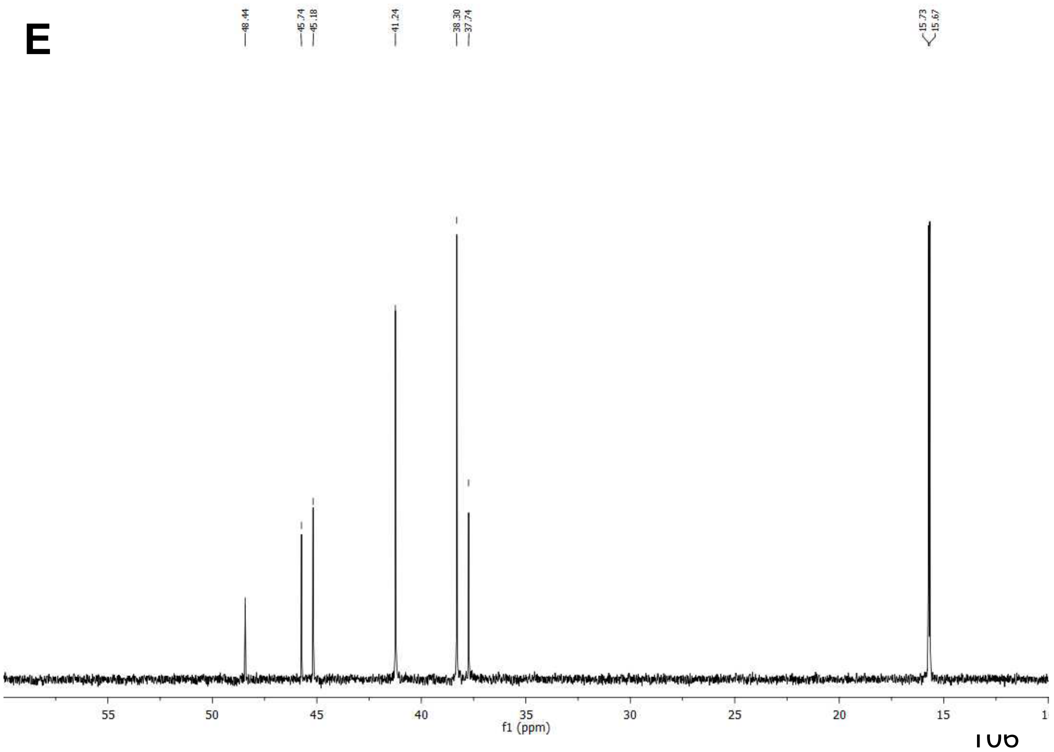
48.44
45.74
45.18
41.24
38.30
37.74
15.73
15.67
55
50
45
40
35
30
25
20
15
f1 (ppm)

F

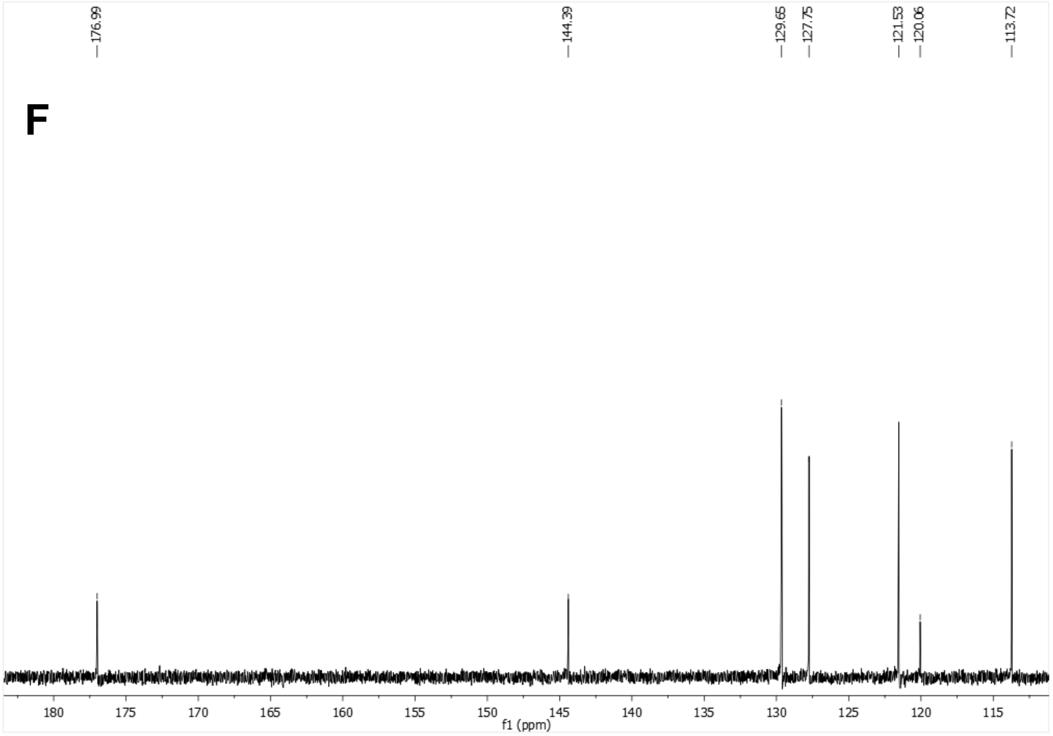
176.99
144.39
129.65
127.75
121.53
120.06
113.72
180
175
170
165
160
155
150
145
140
135
130
125
120
115
f1 (ppm)

112 HSQC ($CDCl_3$)

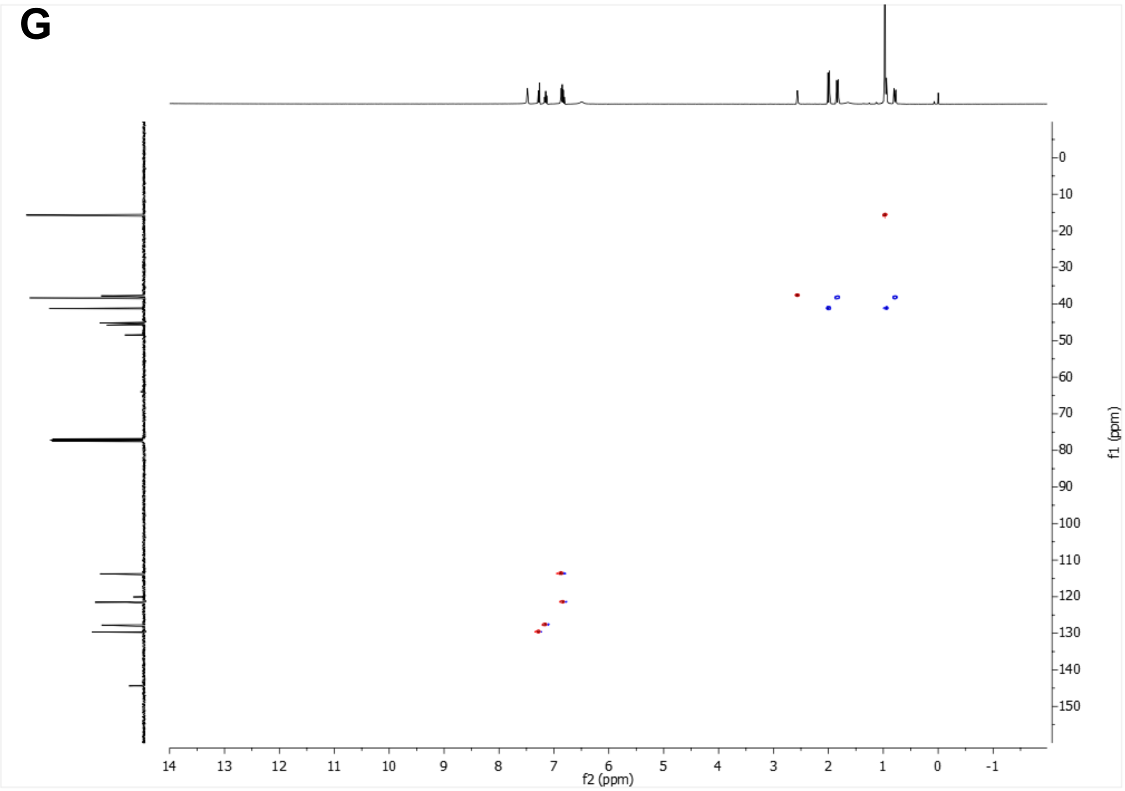


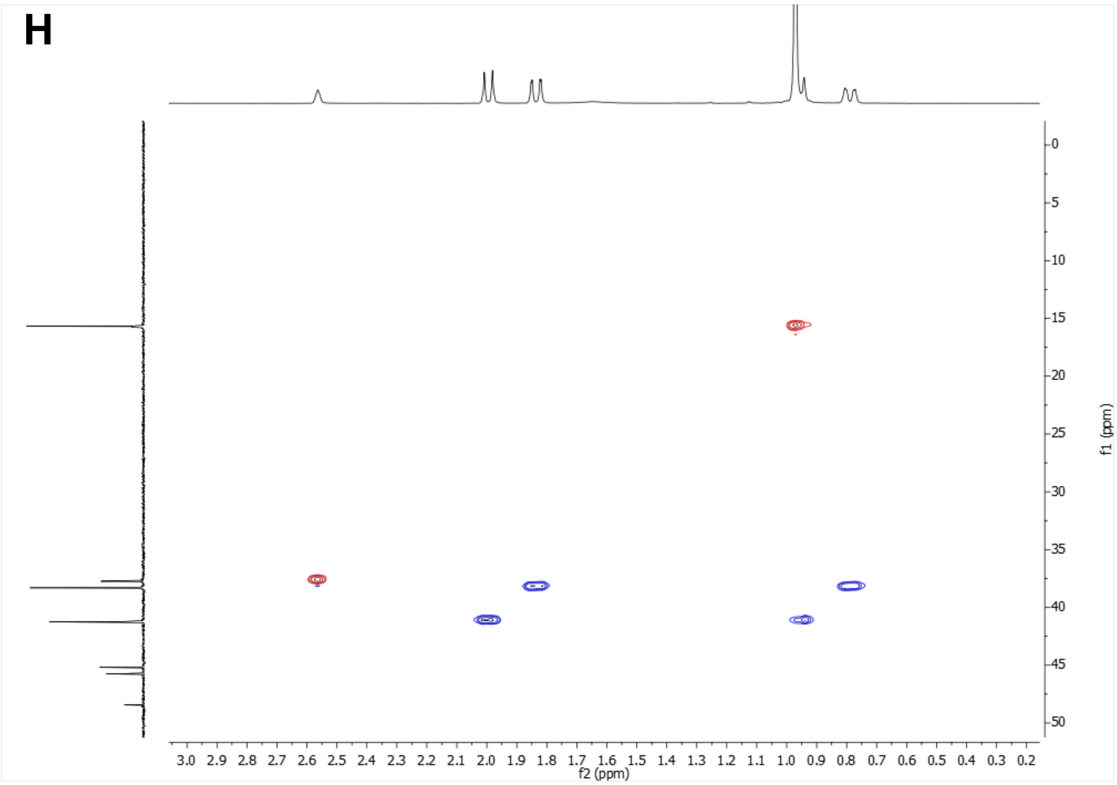

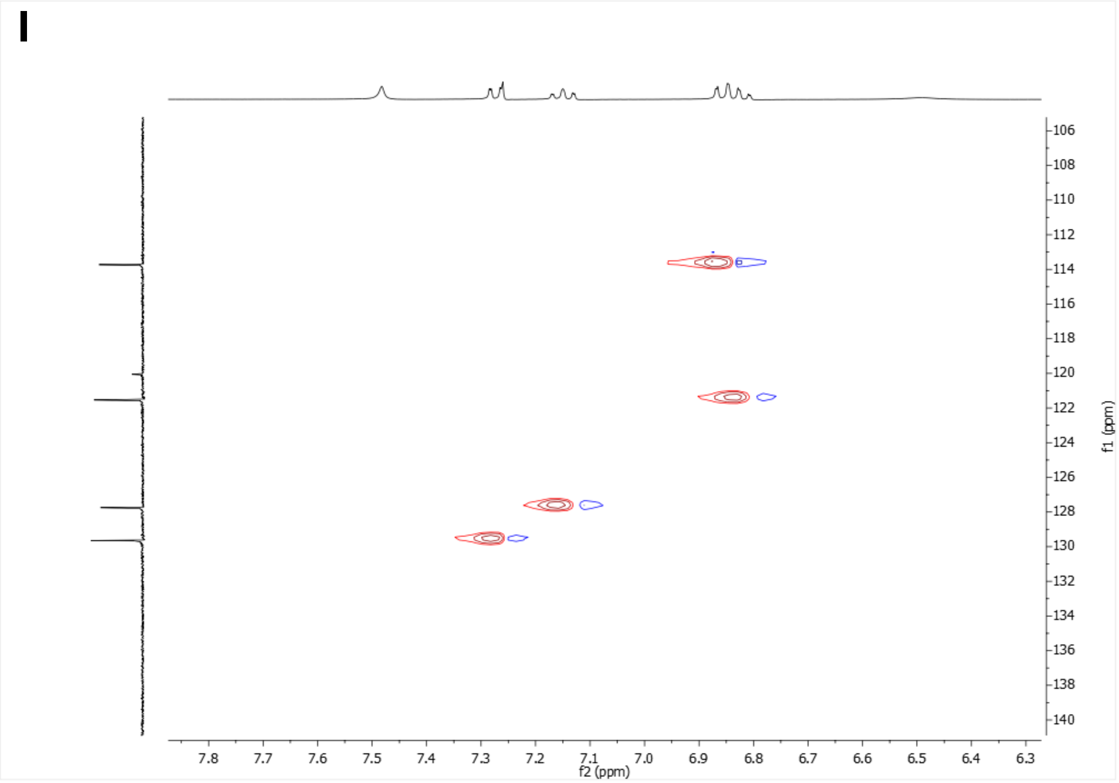


**Supplementary Figure 2: Characterization of *N'*-(2-chlorophenyl)-3,4,8,9-tetramethyltetracyclo[4.4.0.0$^{3,9}$.0$^{4,8}$]decane-1-carbohydrazide (UB-MBX-46).** (A-C) $^{1}$H NMR spectrum. (D-F) $^{13}$C-NMR spectrum. (G-I) HSQC spectrum.

174 $^1H$ NMR (400 MHz, $CDCl_3$)

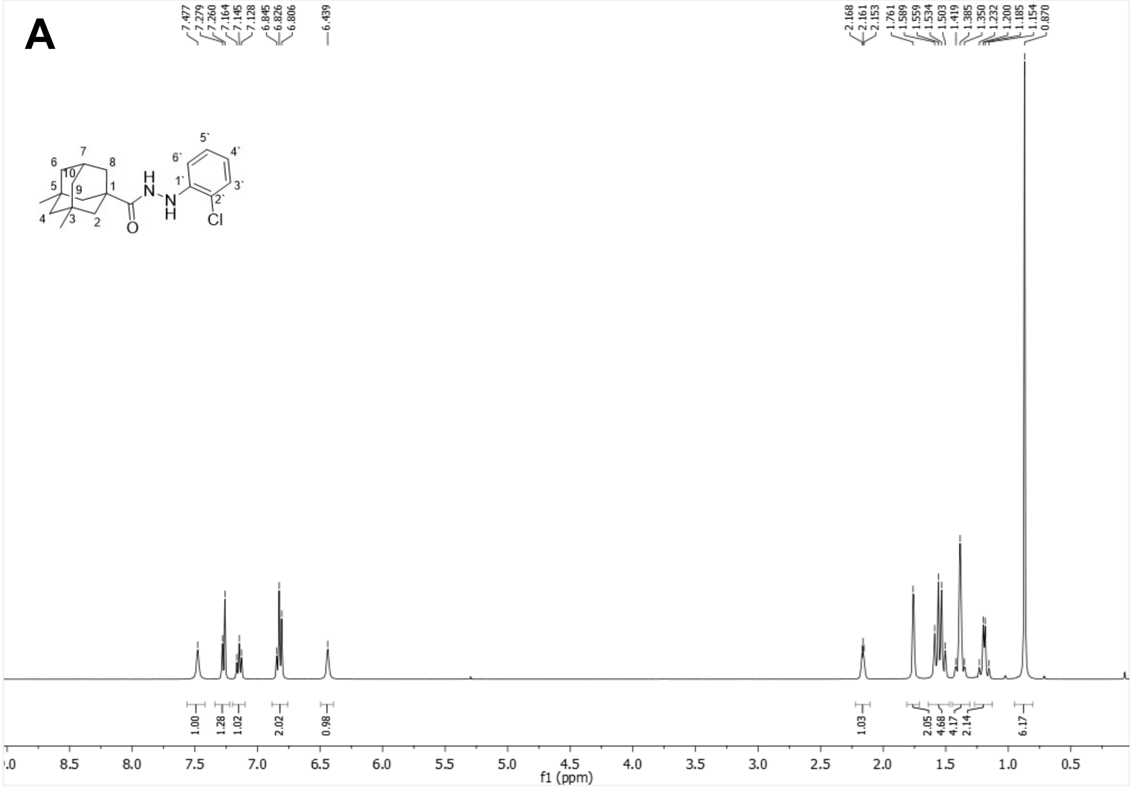


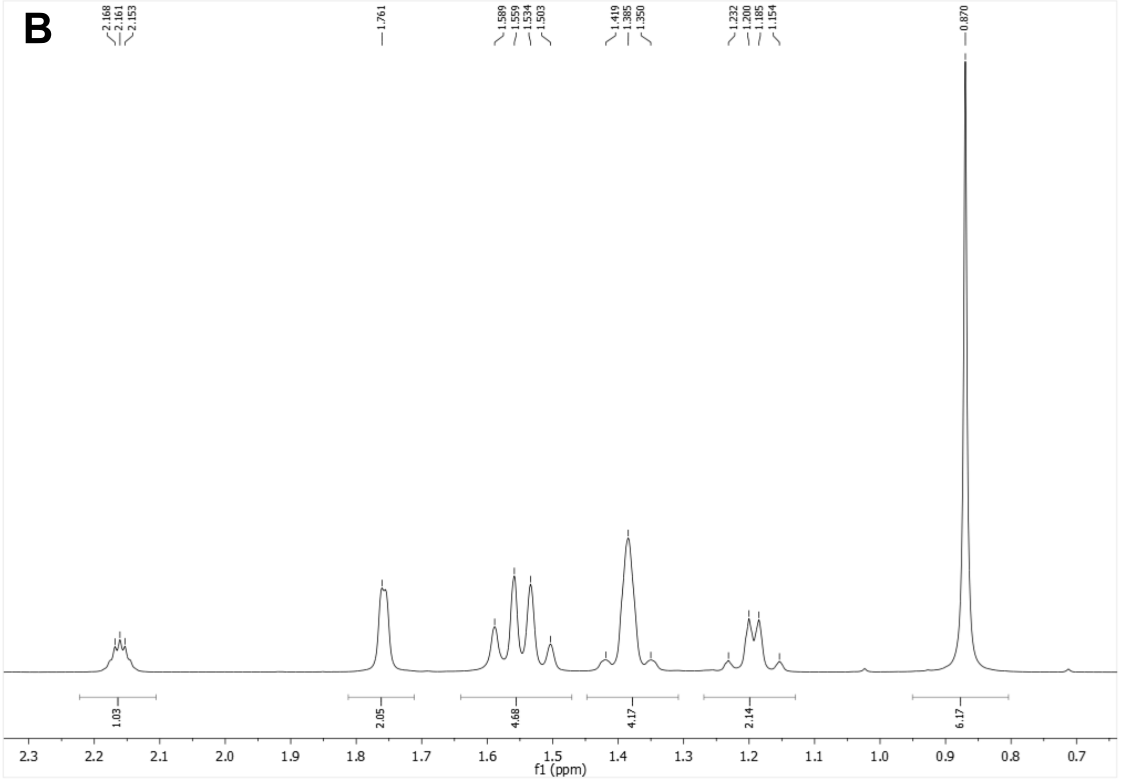

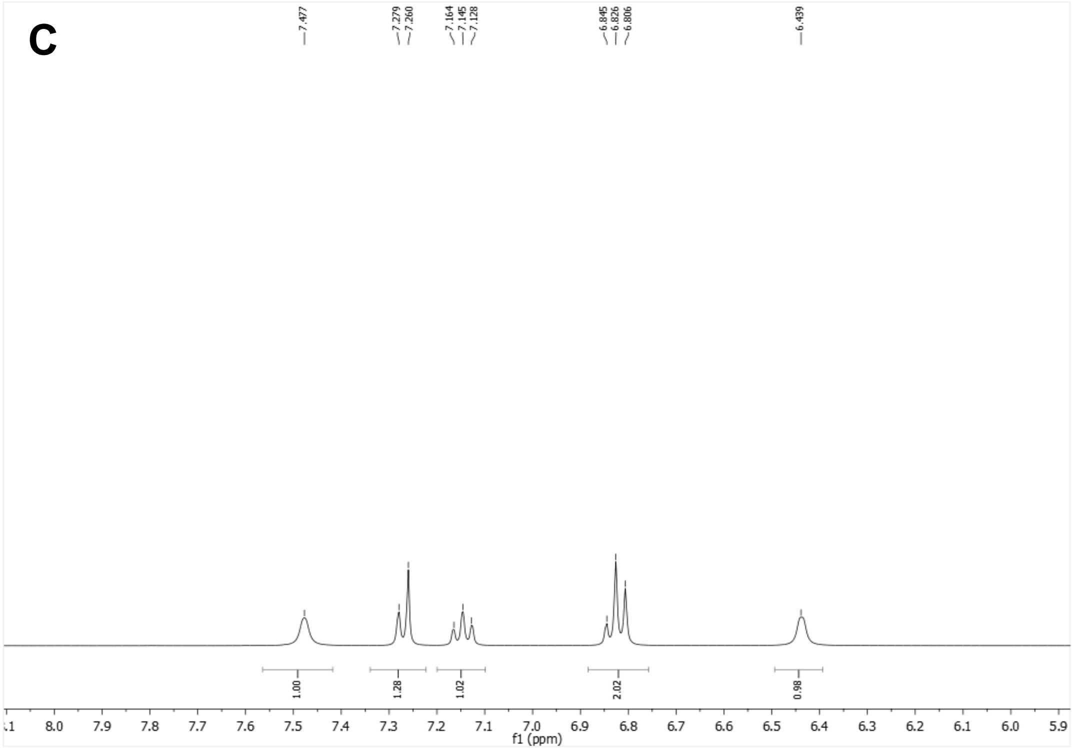


$^{13}C$-NMR (100.6 MHz, $CDCl_3$)

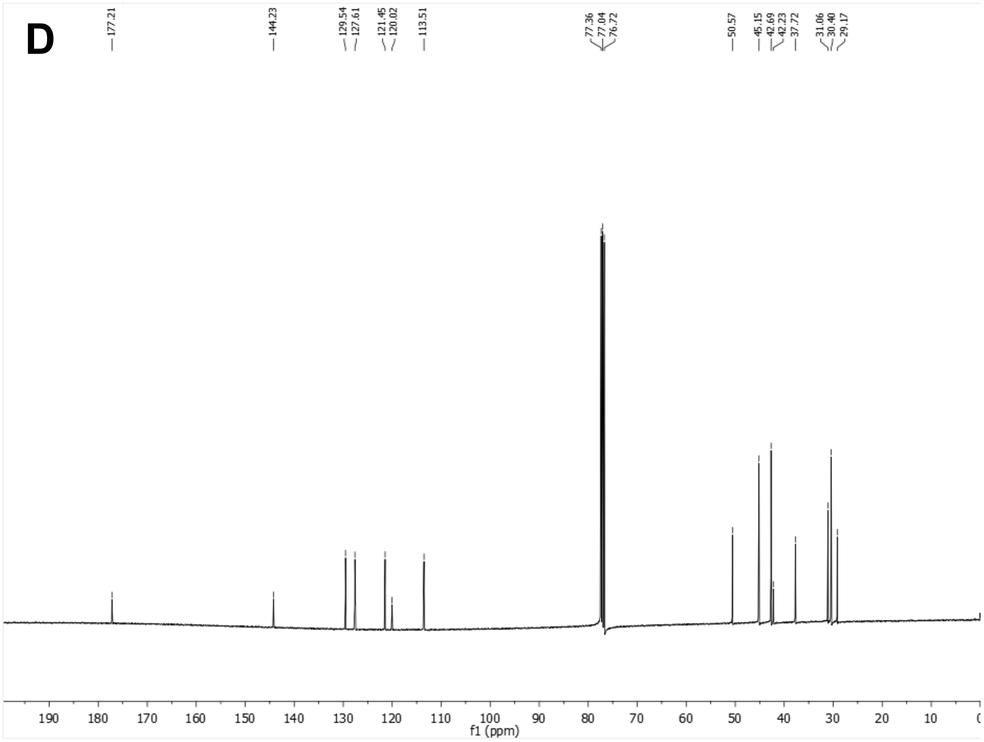

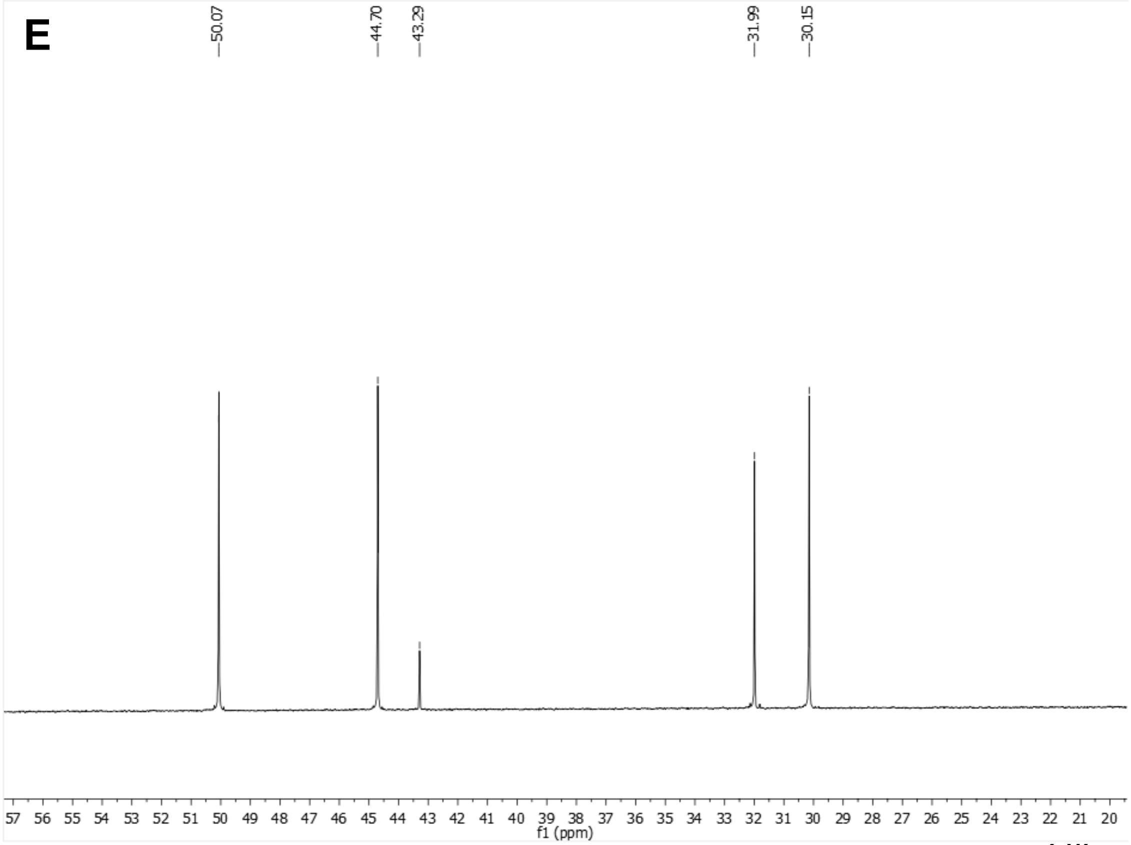
E
50.07
44.70
43.29
31.99
30.15
57 56 55 54 53 52 51 50 49 48 47 46 45 44 43 42 41 40 39 38 37 36 35 34 33 32 31 30 29 28 27 26 25 24 23 22 21 20
f1 (ppm)

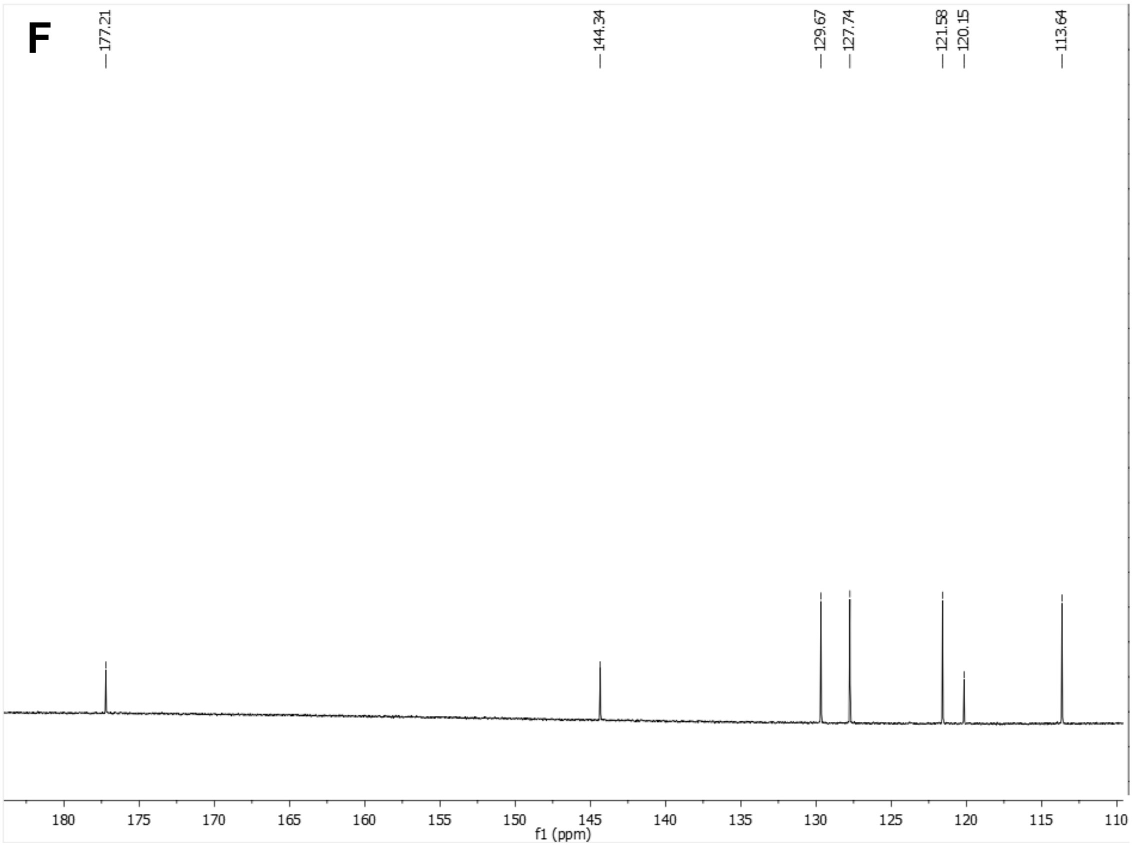
F
177.21
144.34
129.67
127.74
121.58
120.15
113.64
180 175 170 165 160 155 150 145 140 135 130 125 120 115 110
f1 (ppm)

HSQC ($CDCL_3$)

G

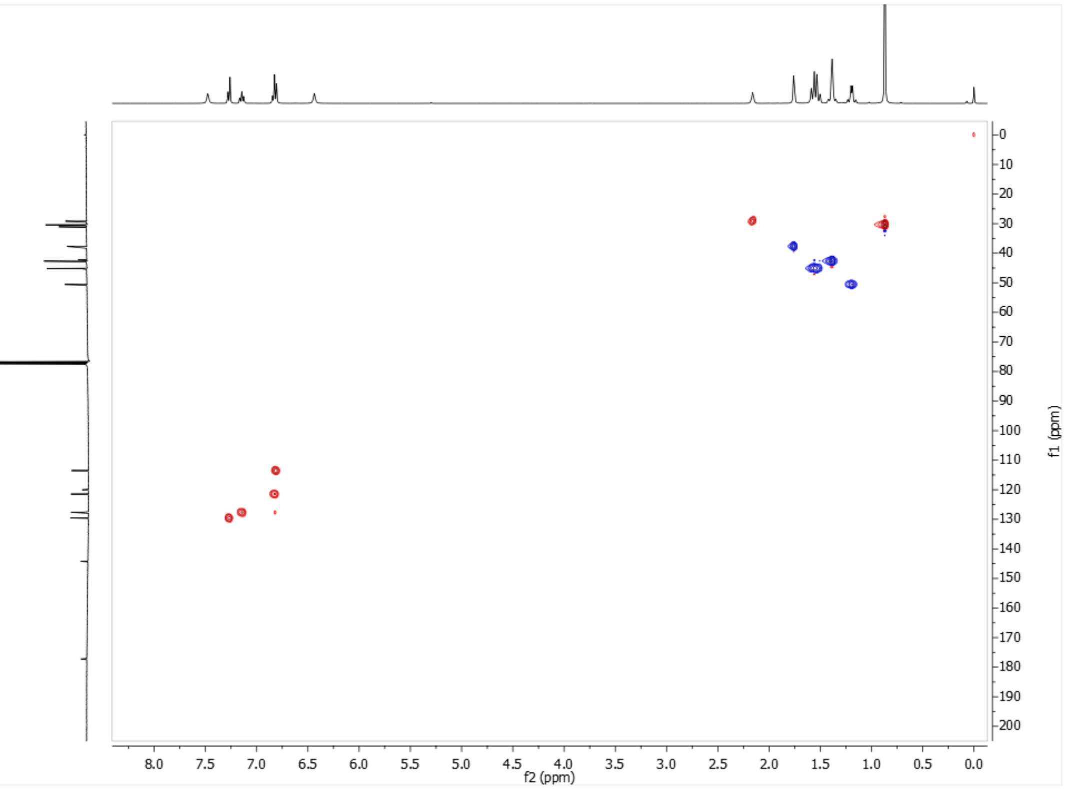


H

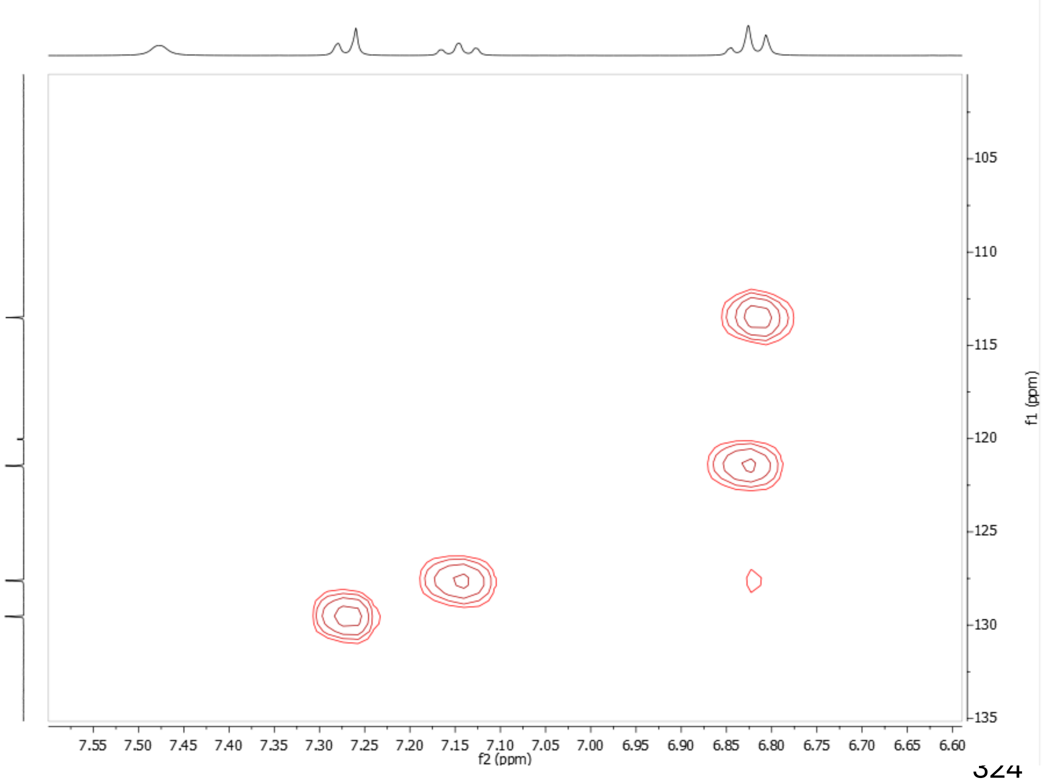

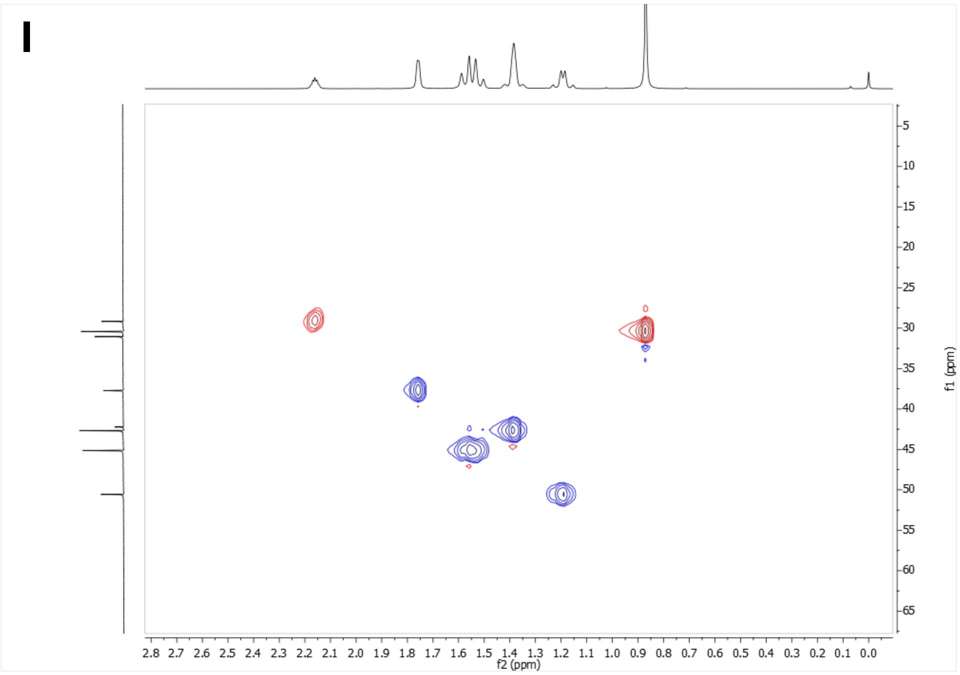


COSY ($CDCl_3$)

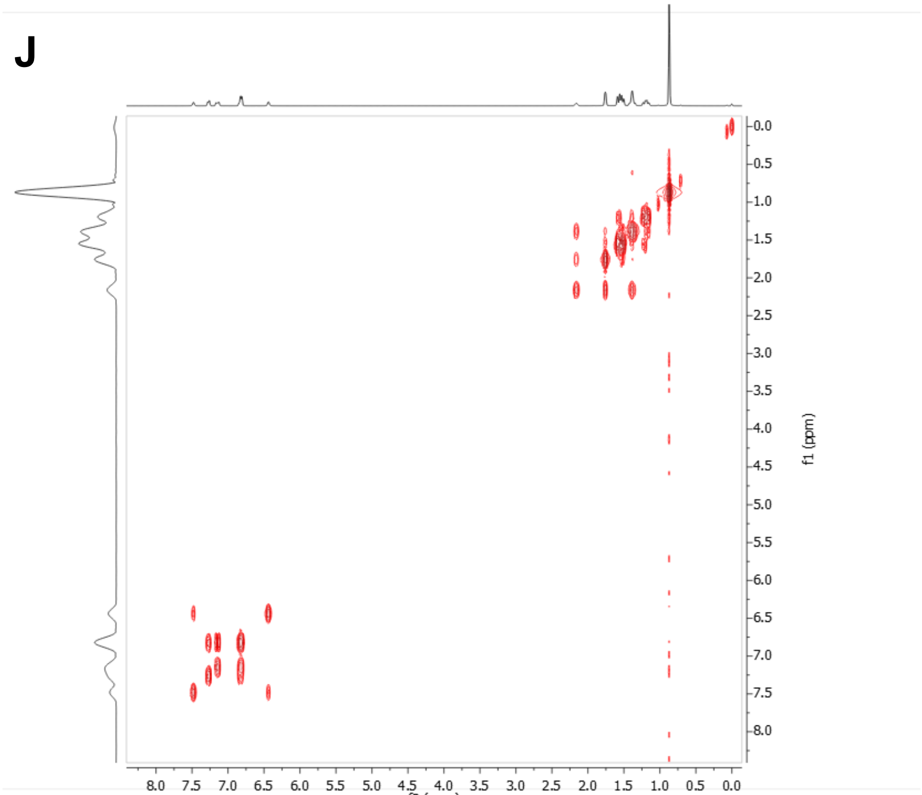


**Supplementary Figure 3: Characterization of *N'*-(2-chlorophenyl)-3,5-dimethyladamantane-1-carbohydrazide (UB-ALT-36).** (A-C) $^{1}H$ NMR spectrum. (D-F) $^{13}C$-NMR spectrum. (G-I) HSQC spectrum. (J) COSY spectrum.

$^1$H NMR (400 MHz, $CDCl_3$)

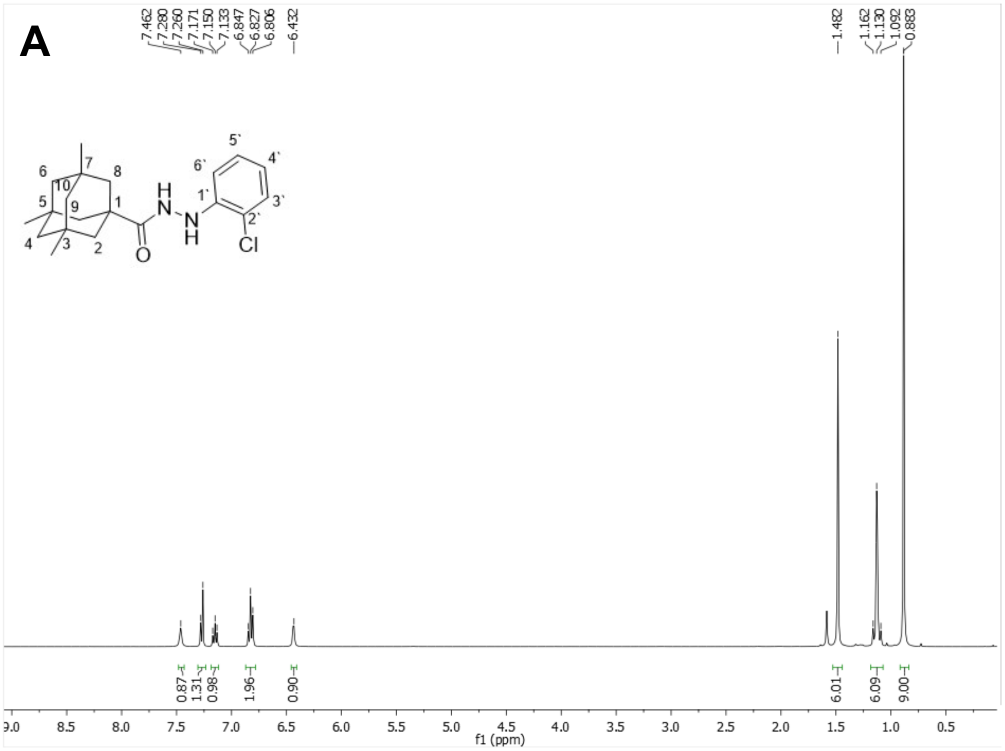


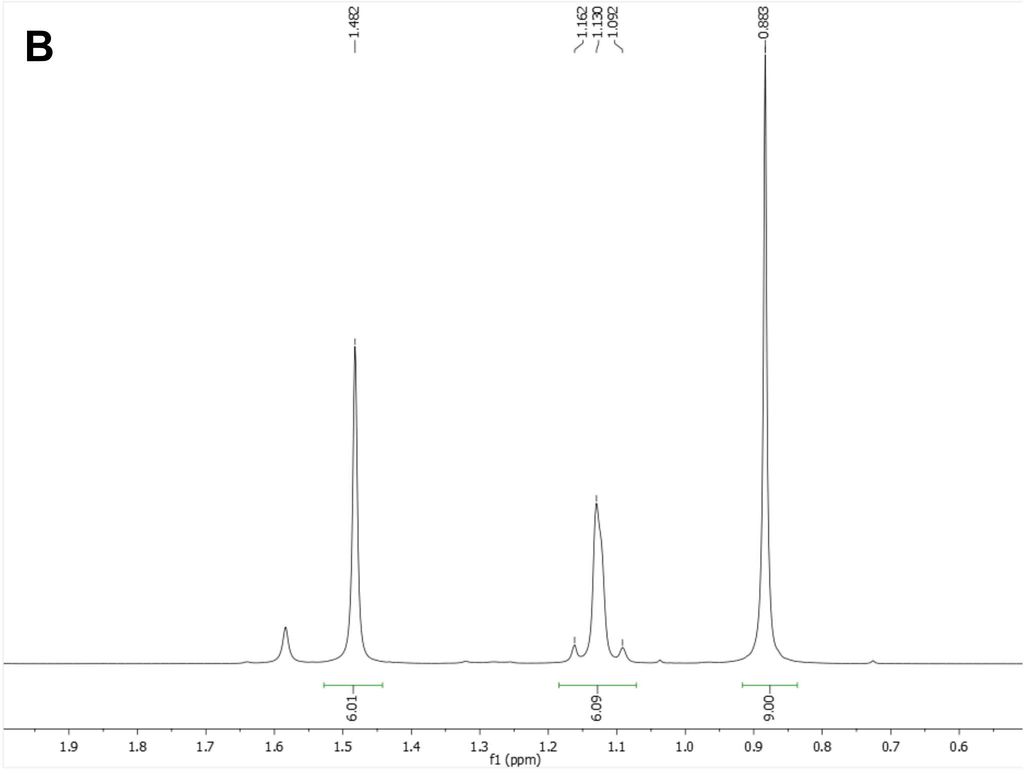

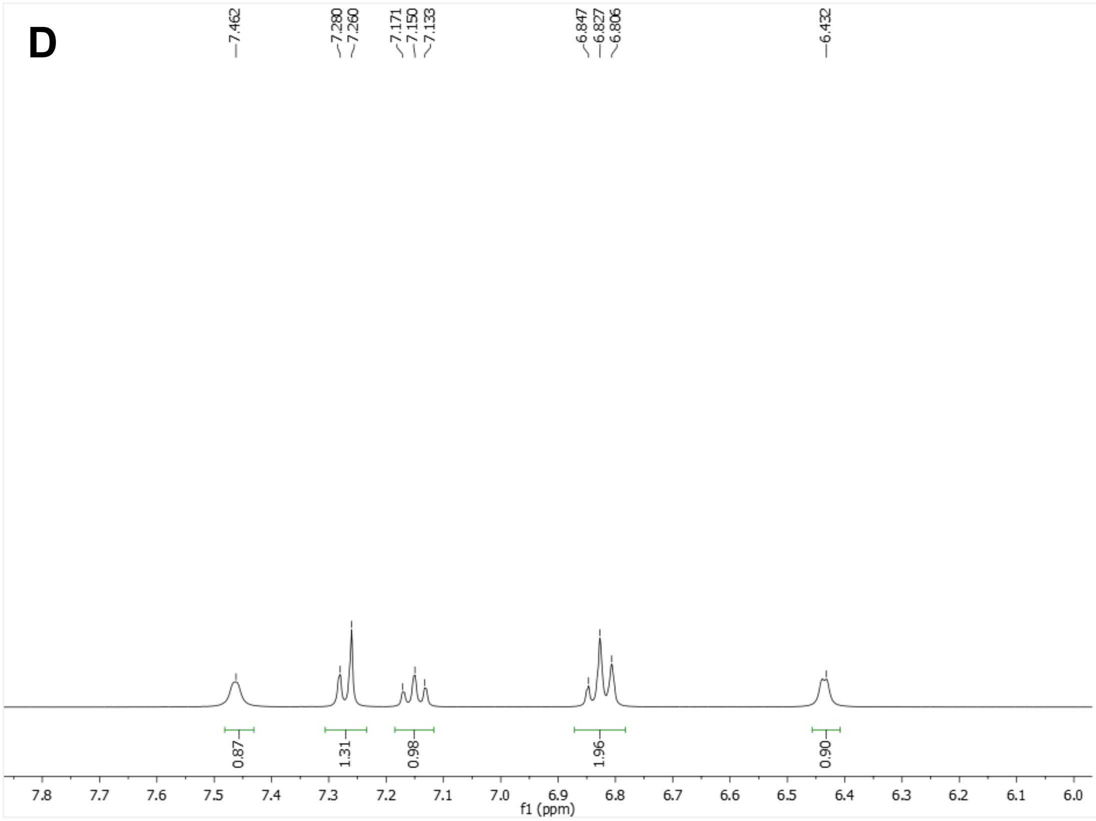


$^{13}$C-NMR (100.6 MHz, $CDCl_3$)

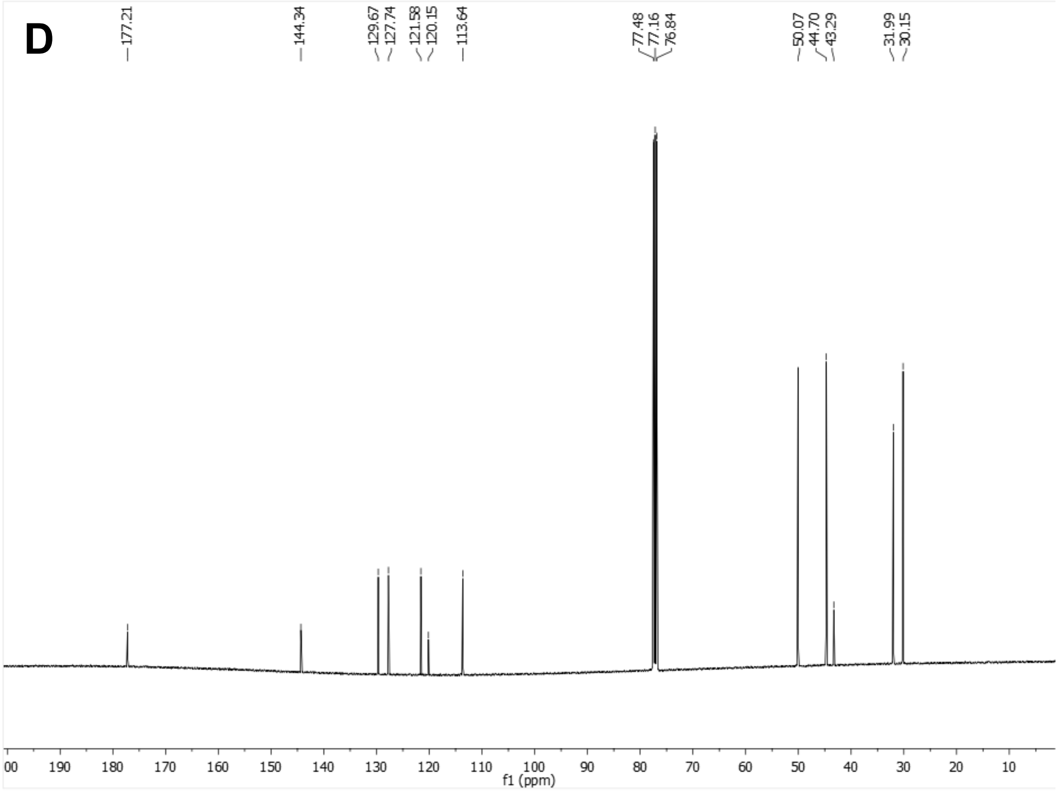

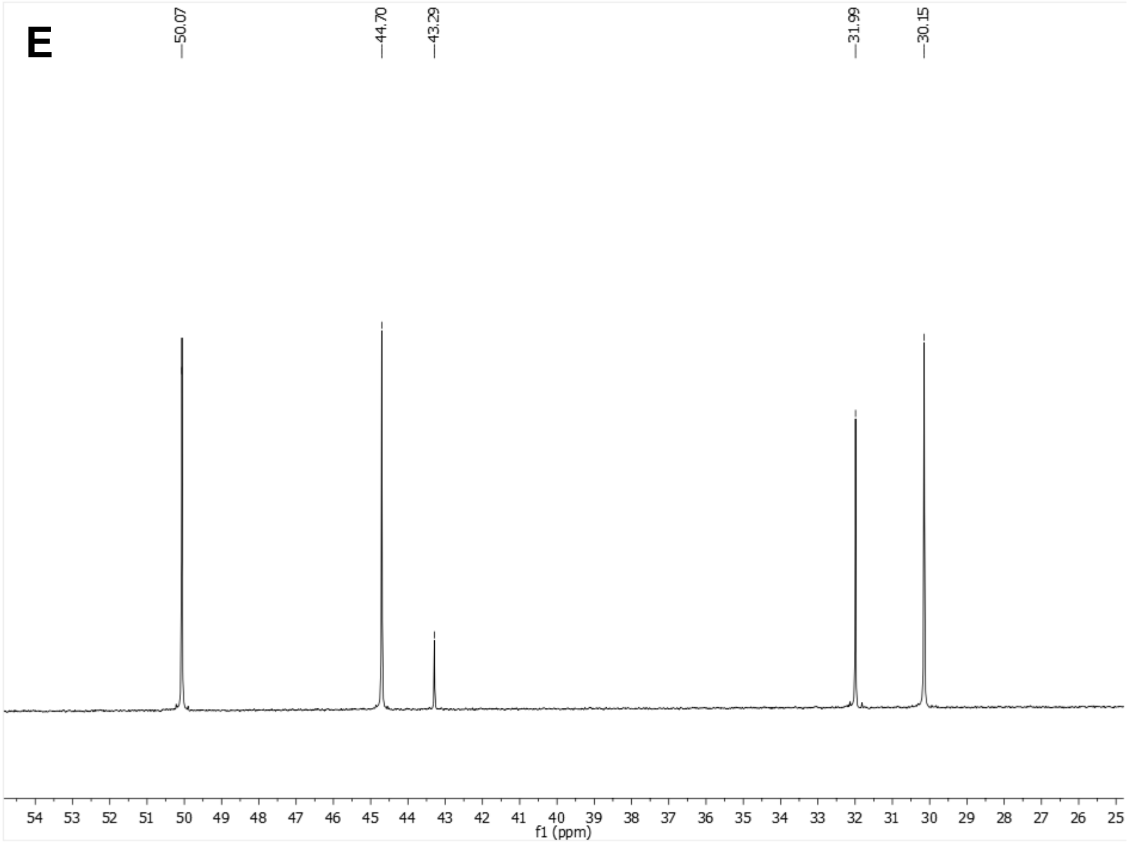
E
50.07
44.70
43.29
31.99
30.15
54 53 52 51 50 49 48 47 46 45 44 43 42 41 40 39 38 37 36 35 34 33 32 31 30 29 28 27 26 25
f1 (ppm)

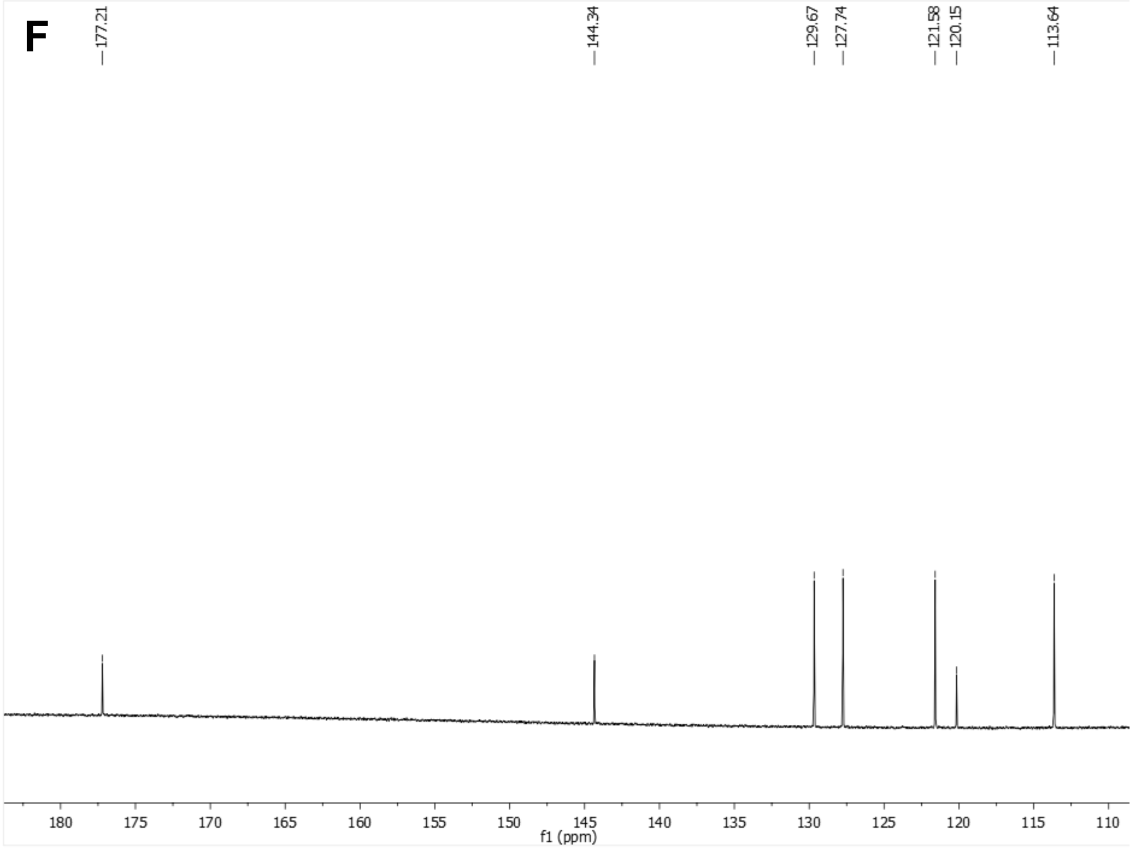
F
177.21
144.34
129.67
127.74
121.58
120.15
113.64
180 175 170 165 160 155 150 145 140 135 130 125 120 115 110
f1 (ppm)

HSQC ($CDCl_3$)

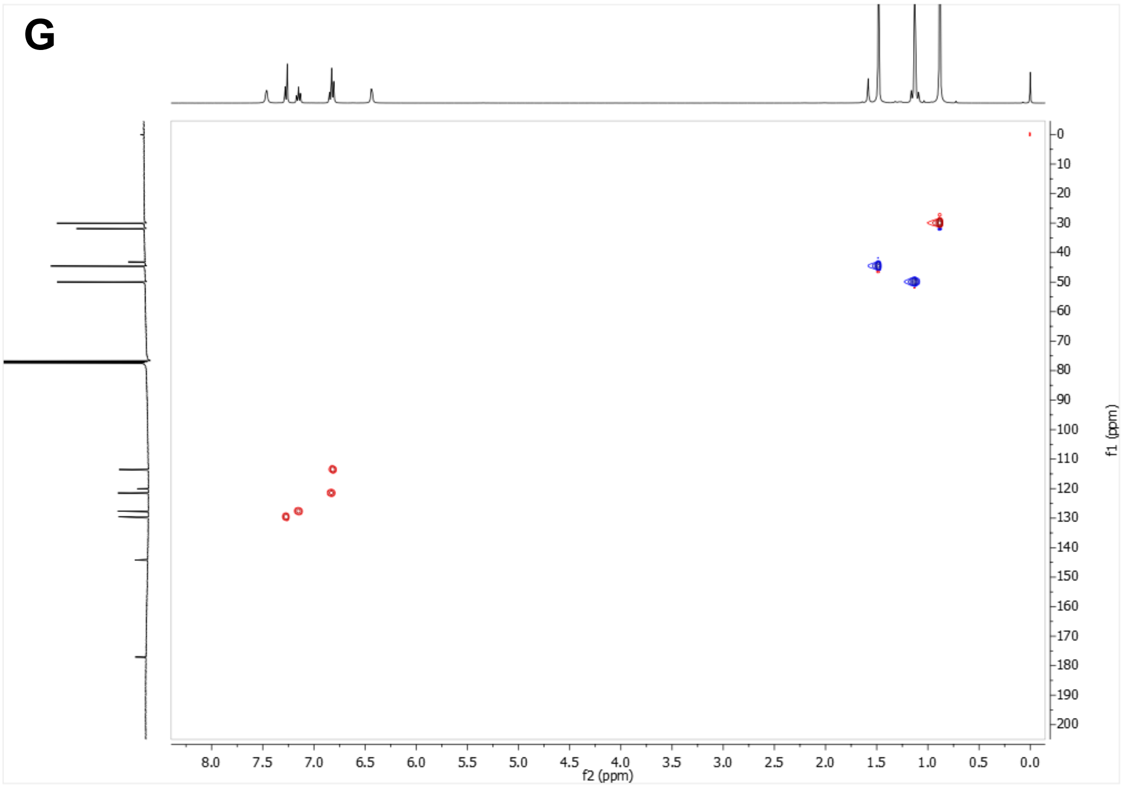


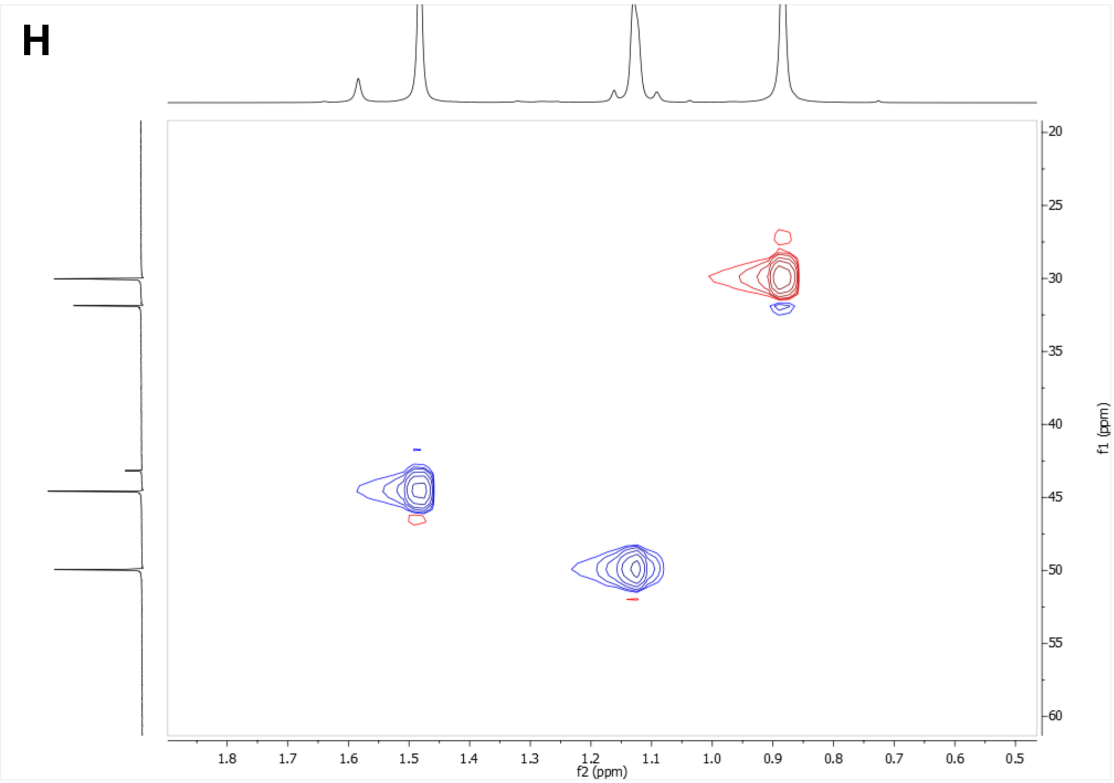

I

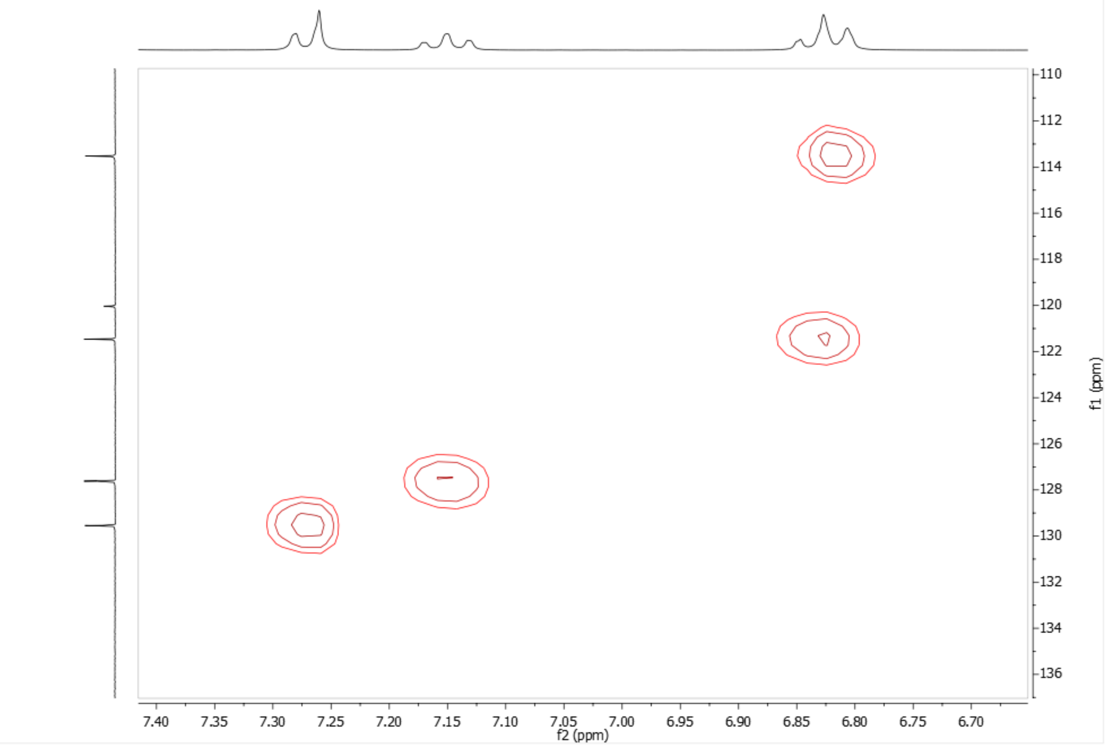


528 COSY ($CDCl_3$)

J

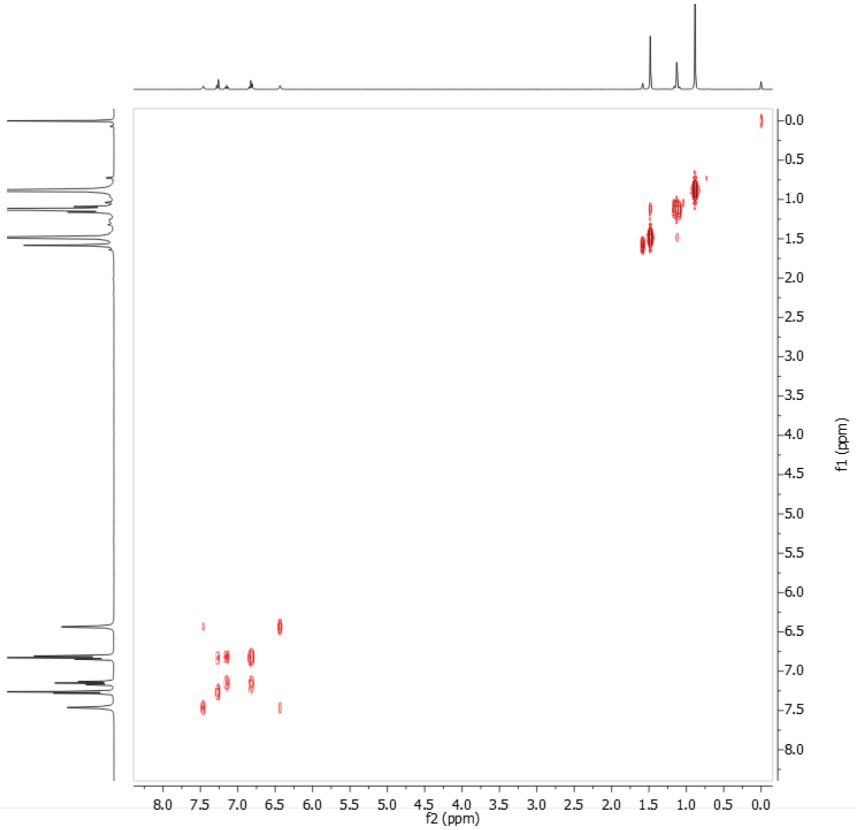

**Supplementary Figure 4: Characterization of *N'*-(2-chlorophenyl)-3,5,7-trimethyladamantane-1-carbohydrazide (UB-ALT-P37).** (A-C) $^{1}$H NMR spectrum. (D-F) $^{13}$C-NMR spectrum. (G-I) HSQC spectrum. (J) COSY spectrum.

$^{1}H$ NMR (400 MHz, $CDCl_3$

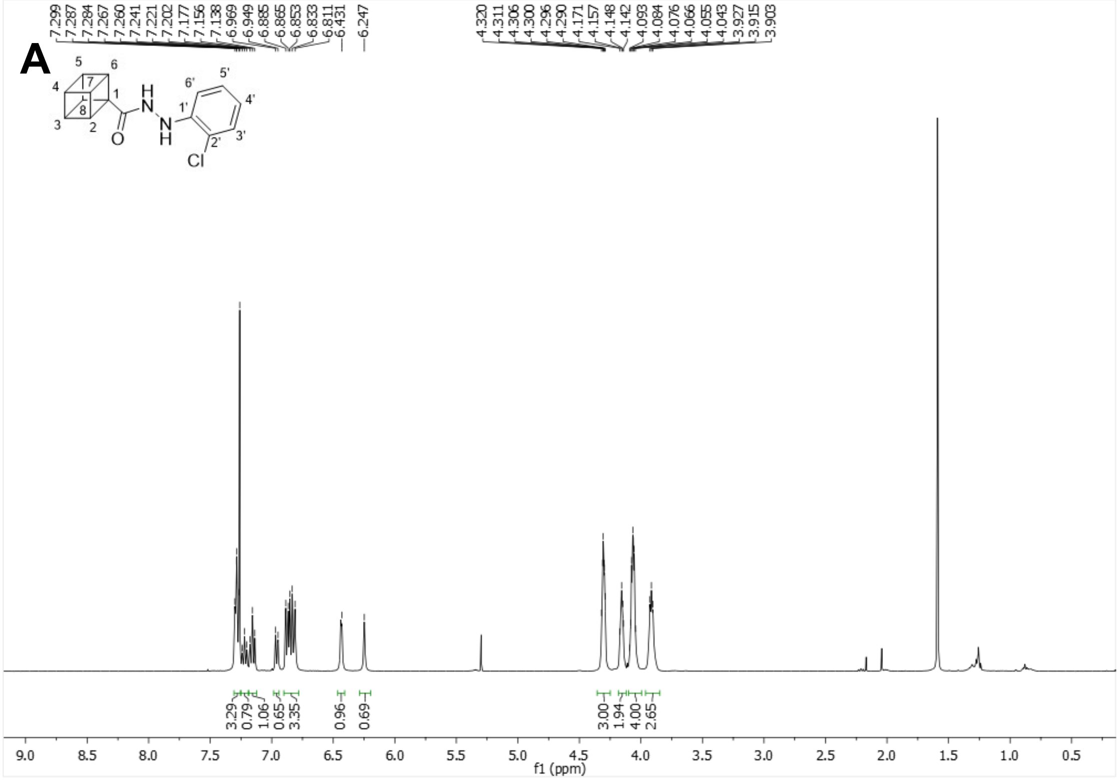


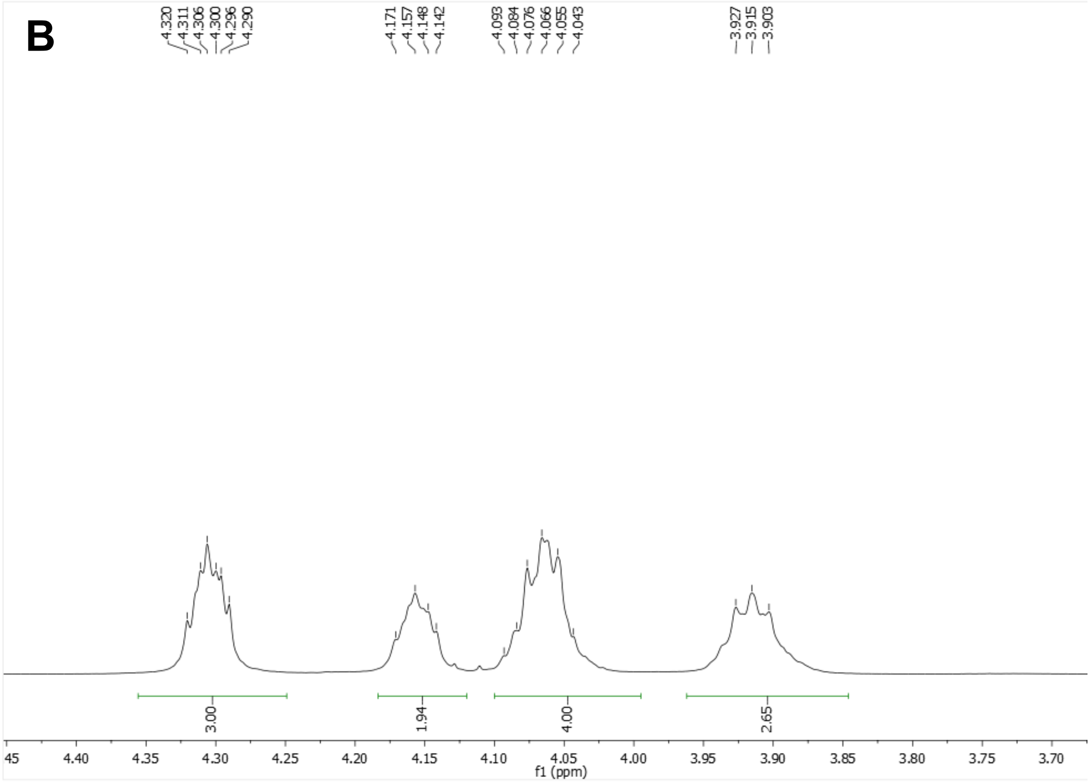

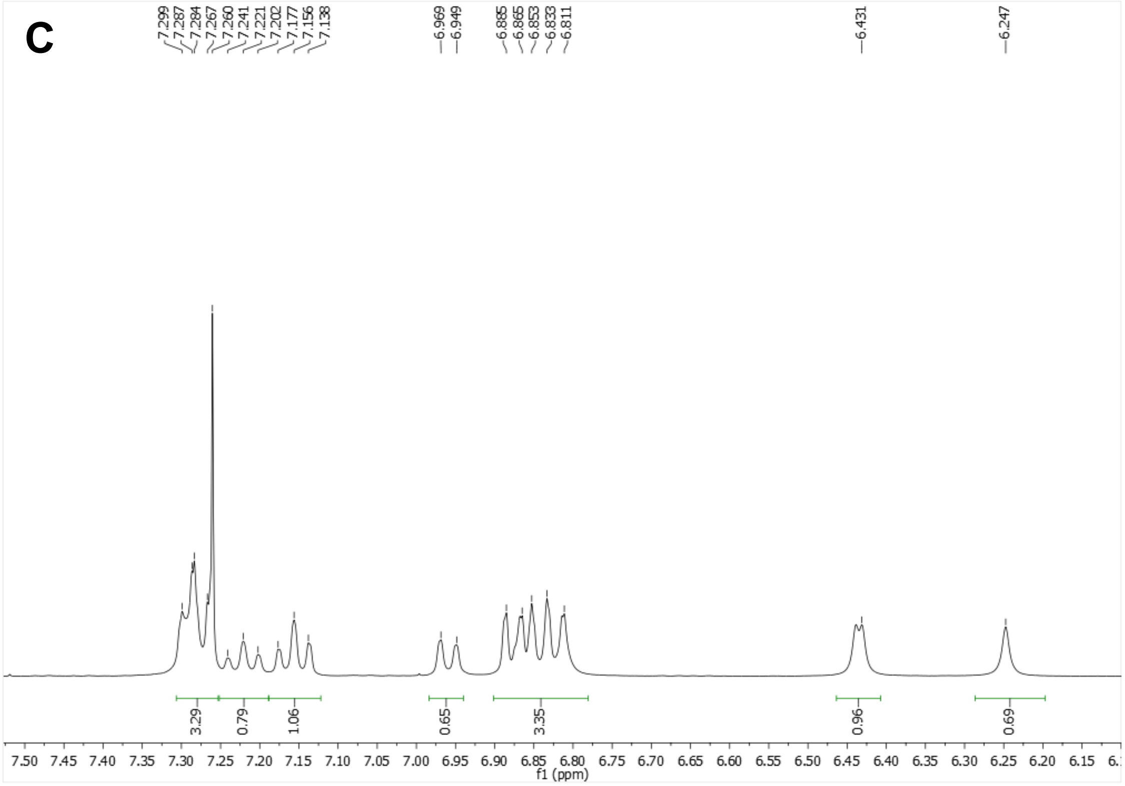


$^{13}$C-NMR (100.6 MHz, $CDCl_3$)

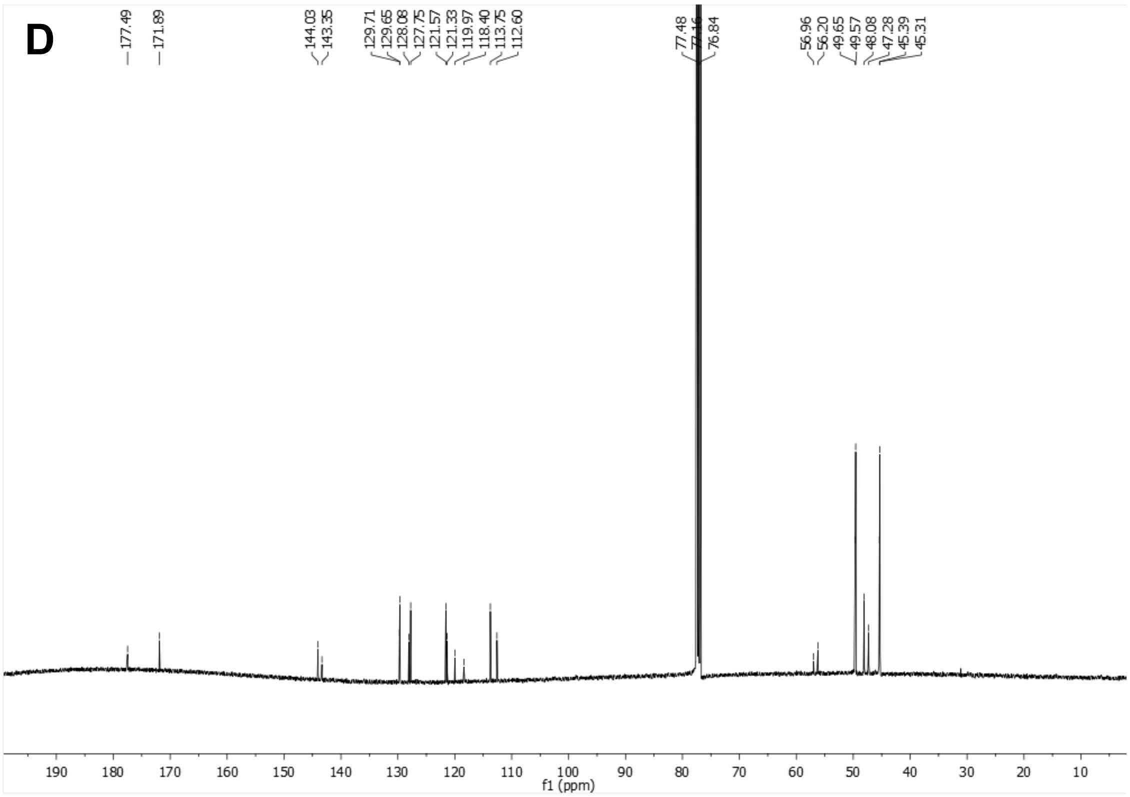

E

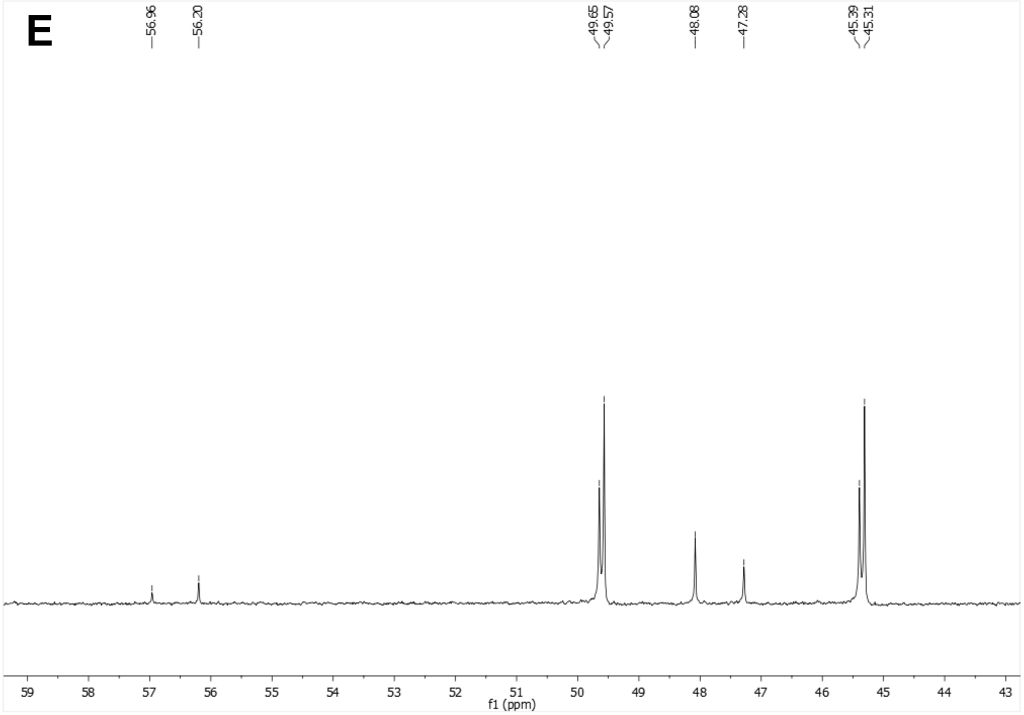
56.96
56.20
49.65
49.57
48.08
47.28
45.39
45.31
f1 (ppm)

F

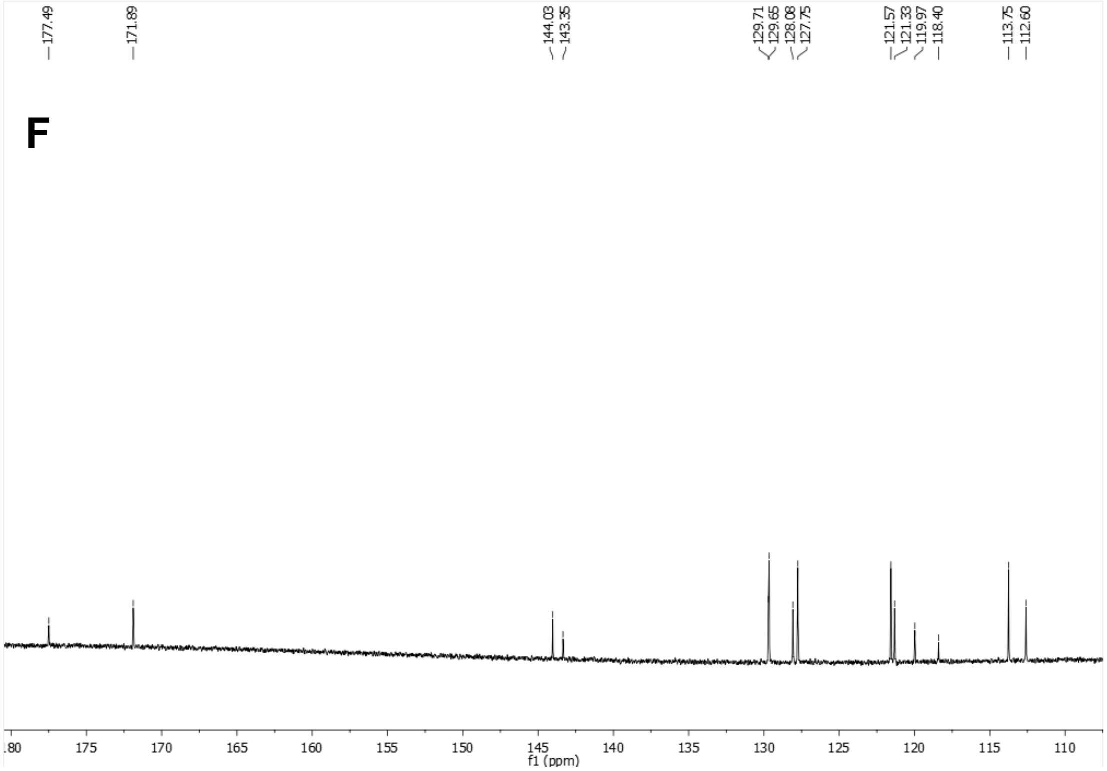
177.49
171.89
144.03
143.35
129.71
129.65
128.08
127.75
121.57
121.33
119.97
118.40
113.75
112.60
f1 (ppm)

HSQC ($CDCl_3$)

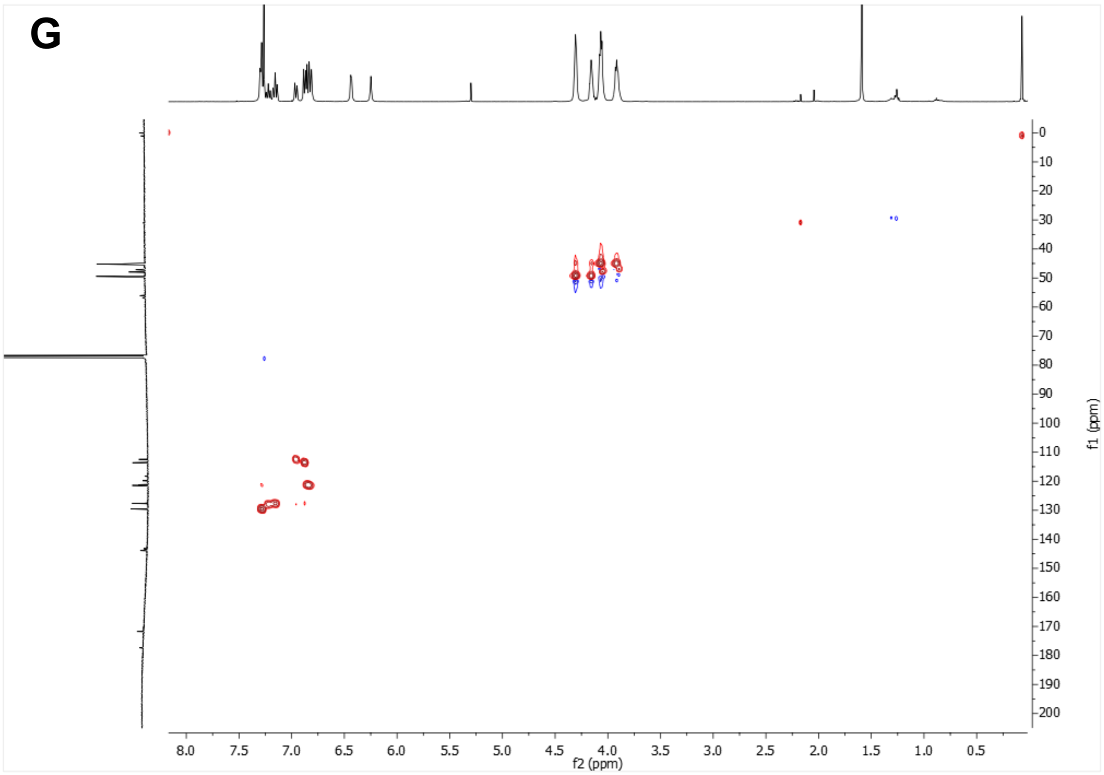


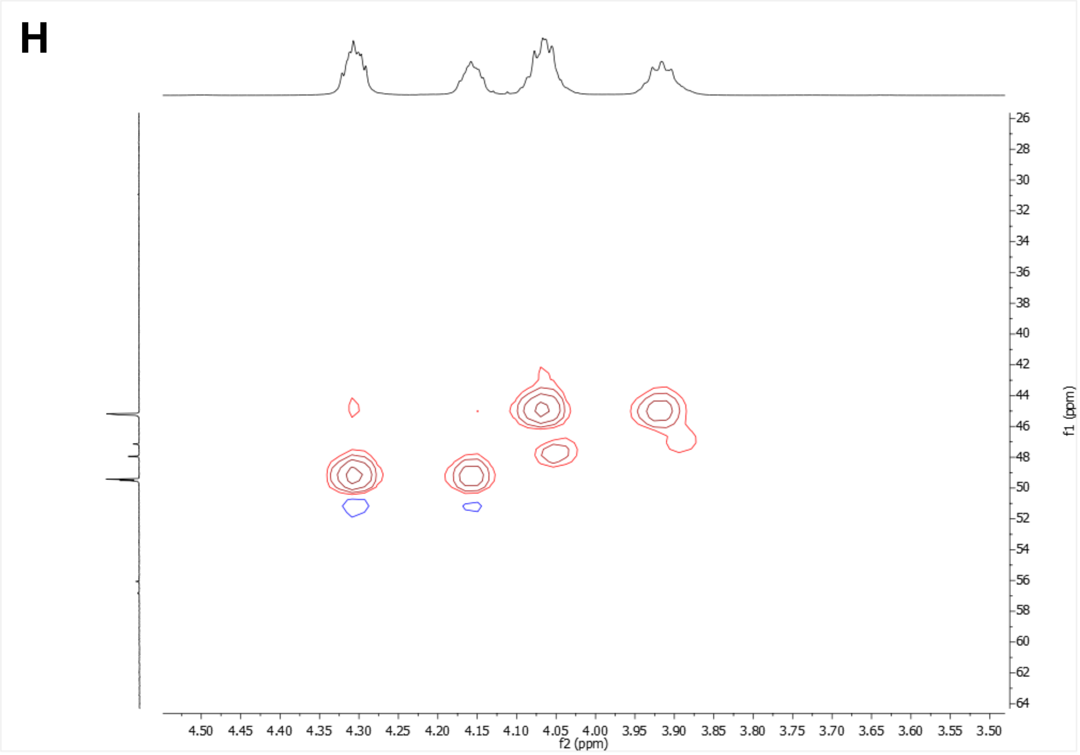

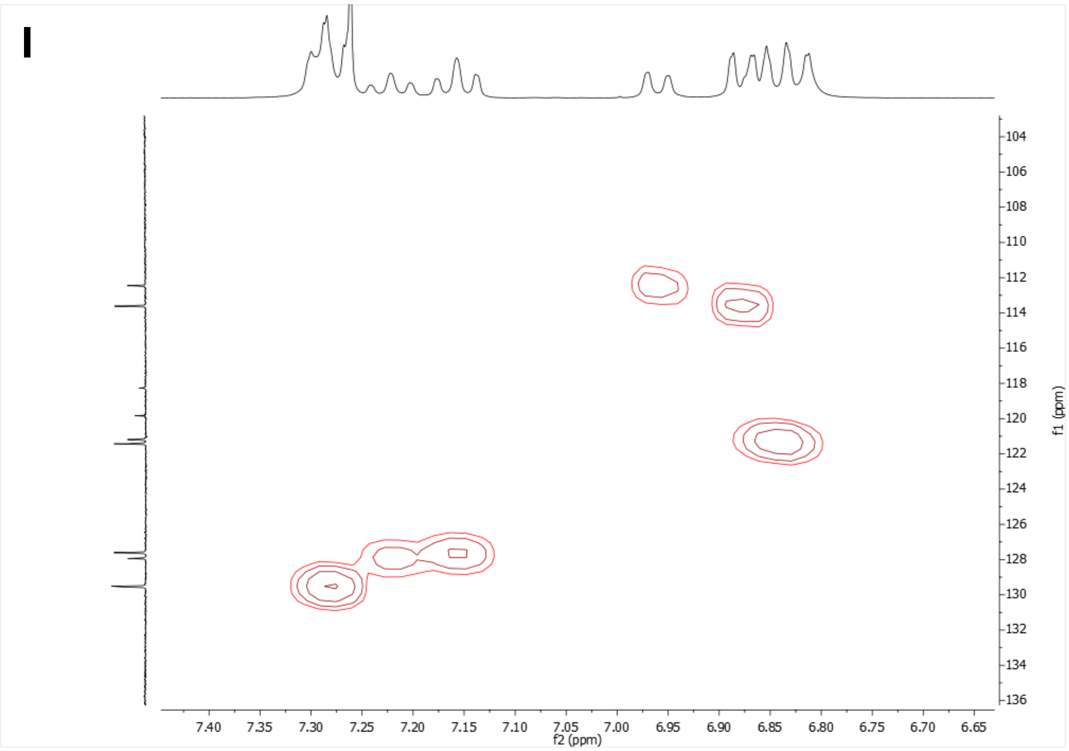


COSY ($CDCl_3$)

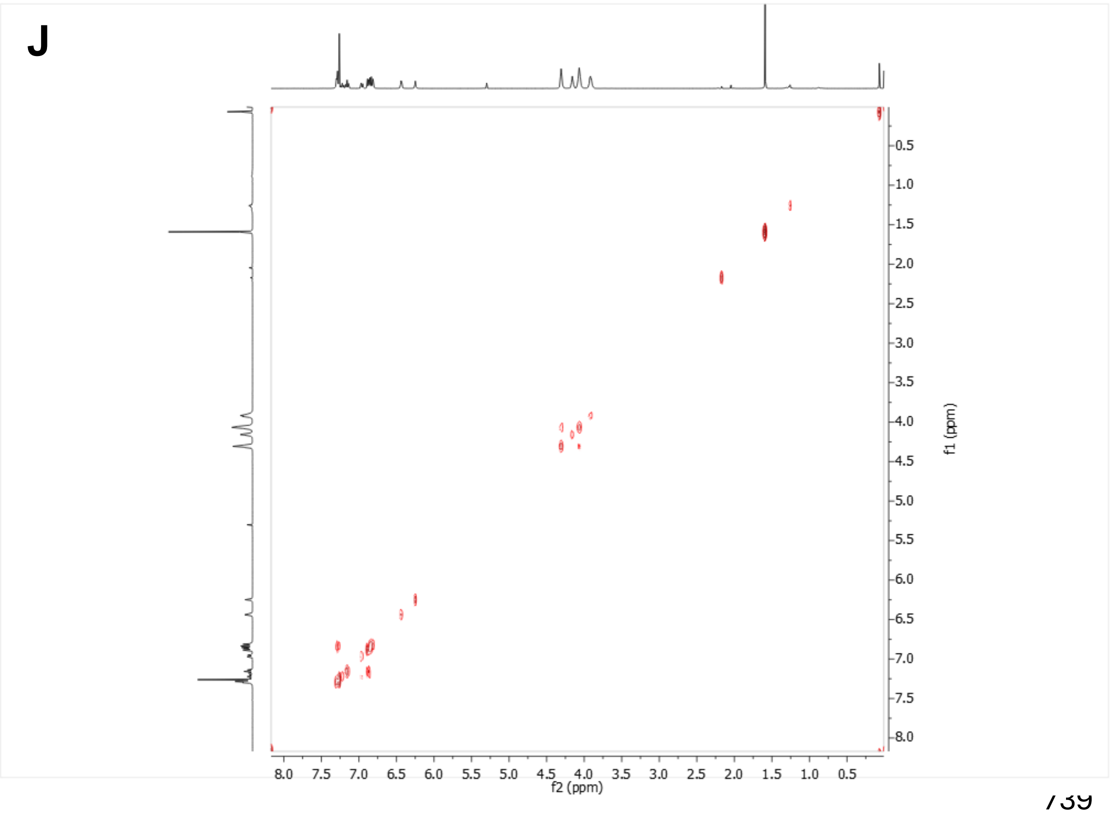

**K**

### HPLC-UV at 254 nm

**Blank**

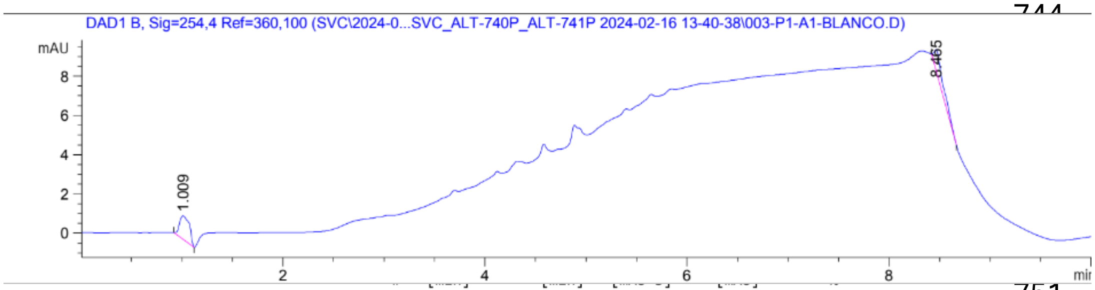


| | | | | | | |
|---|---|---|---|---|---|---|
| 1 | 1.009 | BB | 0.1022 | 8.31069 | 1.17909 | 53.3178 |
| 2 | 8.465 | BB | 0.0943 | 7.27639 | 1.05744 | 46.6822 |
| Totals : | | | | 15.58708 | 2.23653 | |

**UB-ALT-P38**

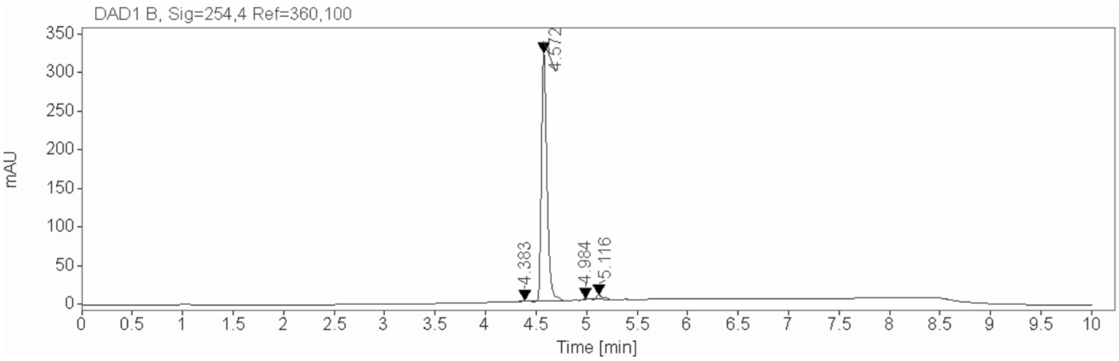


**Signal:** DAD1 B, Sig=254,4 Ref=360,100

| RT [min] | Type | Width [min] | Area | Height | Area% | Name |
|---|---|---|---|---|---|---|
| 4.383 | BB | 0.0579 | 7.7845 | 1.9803 | 0.6080 | |
| 4.572 | BB | 0.0584 | 1236.1952 | 324.8564 | 96.5513 | |
| 4.984 | BB | 0.0585 | 8.4846 | 2.2238 | 0.6627 | |
| 5.116 | BB | 0.0657 | 27.8861 | 6.0589 | 2.1780 | |
| | | Sum | 1280.3504 | | | |

**Supplementary Figure 5: Characterization of *N'*-(2-chlorophenyl)cubane-1-carbohydrazide (UB-ALT-P38).** (A-C) $^{1}$H NMR spectrum. (D-F) $^{13}$C-NMR spectrum. (G-I) HSQC spectrum. (J) COSY spectrum. (K) HPLC-UV at 254 nm.

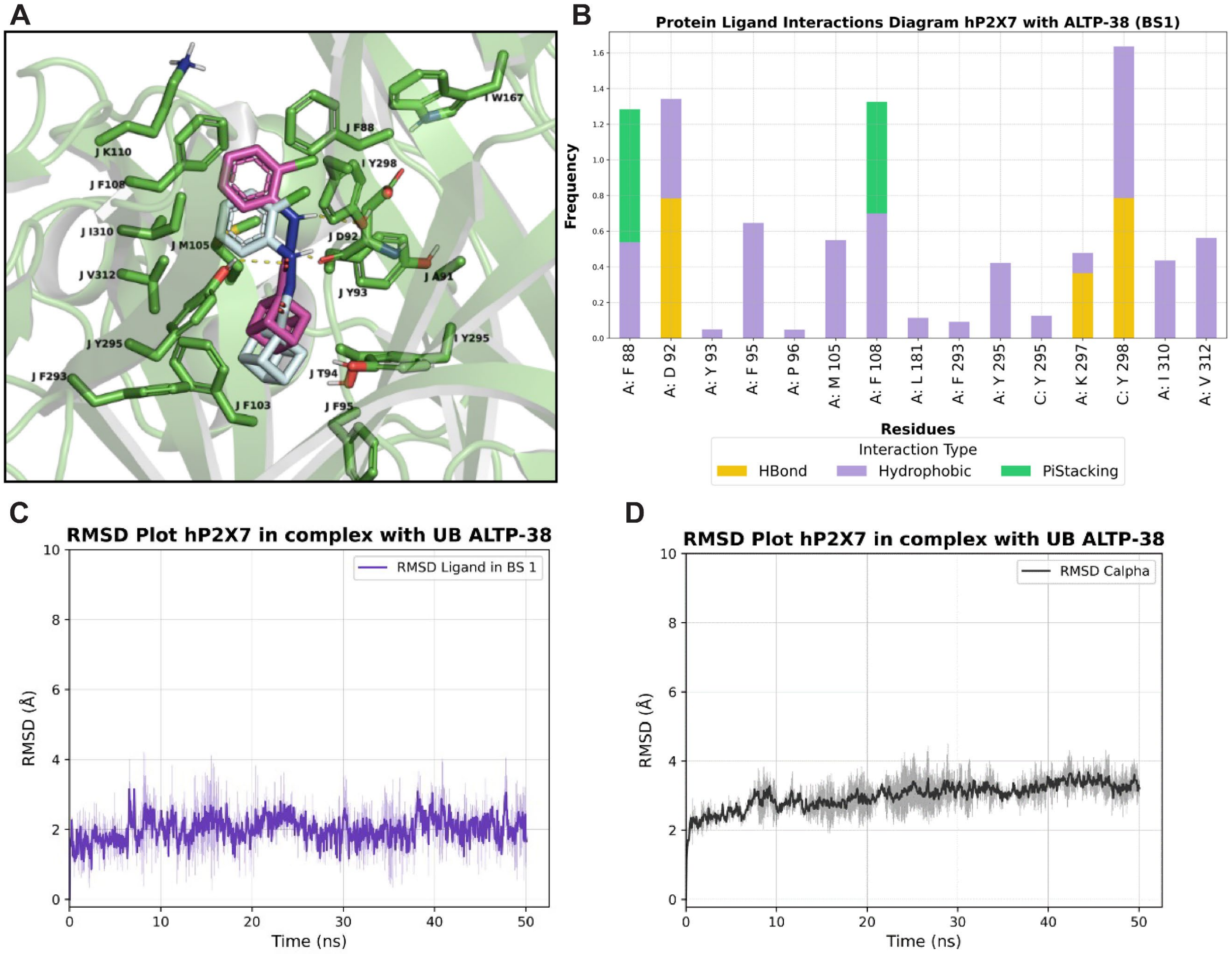


**Supplementary Fig. 6.** MD simulation results of UB-ALT-P38 bound to the hP2X7R in POPC bilayers using ff19sb to model the protein, lipid21force field to model the POPC lipids, and GAFF2 to model the ligand implemented within Amber22 program[3-8]. **(A)** Superposition of the last snapshot from 2 ns-TI/MD simulations with the last snapshot from 50 ns-TI/MD simulations of the hP2X7R in complex with UB-ALT-P38. **(B-D)** Ligand-receptor interactions and movements for one representative UB-ALT-P38 molecule bound to the hP2X7R during MD simulations. **(B)** Protein-ligand interaction frequencies for one UB-ALT-P38 bound to the hP2X7R. Color scheme in bar plots: hydrogen bonding interactions are shown with yellow bars; hydrophobic interactions are shown with magenta bars; cation-$\pi$ interactions are shown with green bars; water bridges with blue bars. Bars are plotted only for residues with interaction frequencies ≥ 0.2. **(C)** RMSD for ligand heavy atoms (RMSD lig) from 50 ns-MD simulations for one representative UB-ALT-P38 molecule bound to the hP2X7R. **(D)** RMSD for Cα carbons (excluding loops) from 50 ns-MD simulations of UB-ALT-P38 bound to the hP2X7R.

### Synthesis and characterization methods

*Chemistry. General methods*

400 MHz $^{1}H$ NMR and 100.6 MHz $^{13}C$ NMR spectra were recorded on a Varian Mercury 400 or a Bruker 400. The chemical shifts are reported in ppm (δ scale) relative to internal tetramethylsilane, or to solvent peak, and coupling constants are reported in Hertz (Hz). Assignments given for the NMR spectra of the new compounds have been carried out based on homocorrelation $^{1}H/^{1}H$ (COSY) and/or heterocorrelation $^{1}H/^{13}C$ (HSQC) experiments. The used abbreviations were: s, singlet; d, doublet; m, multiplet; cs, complex signal; broad s, broad singlet, or combinations thereof. IR spectra were run on a FTIR Perkin-Elmer Spectrum RX I or a Perkin-Elmer Spectrum TWO spectrophotometers, using sodium chloride (NaCl) pellets or attenuated total reflectance (ATR) techniques. Absorption values are expressed as wavenumbers ($cm^{-1}$); only significant absorption bands are given. Column chromatography was performed on silica gel 60 Å (Sigma Aldrich, 40 - 63 µm, 230-400 mesh) or with a CombiFlash Rf 150 Teledine ISCO provided with a UV-Vis detector. Thin Layer Chromatography (TLC) was performed with aluminium-backed sheets with silica gel 60 F254 (Merck, ref 1.05554), and spots were visualized with UV light, 1% aqueous solution of $KMnO_4$, iodine or ninhydrin. Melting points were determined in open capillary tubes with an MFB 59510M Gallenkamp melting point apparatus. Accurate mass spectra were recorded with ESI techniques on a Hewlett-Packard 5988a LC/MSD-TOF instrument or a Thermo LTQ Orbitrap Velos instrument at *Unitat d'Espectrometria de Masses de Caracterització Molecular dels Centres Científics i Tecnològics de la Universitat de Barcelona* (CCiTUB). The elemental analyses were carried out in a Flash 1112 series Thermofinnigan elemental microanalyzator (A5) to determine C, H, and N at the *Servei de Microanàlisis* of IIQAB (CSIC) of Barcelona. The analytical samples of all the new compounds possessed purity ≥95% as evidenced by their elemental analyses or their HPLC-UV. HPLC-UV were determined with a HPLC Agilent 1260 Infinity II LC/MSD coupled to a photodiode array and mass spectrometer. Samples (5 µL, 0.5 mg/mL) in a 1:1 mixture of water with 0.05% formic acid (A) and acetonitrile with 0.05% formic acid (B) were injected using an Agilent Poroshell 120 EC-C18 (2.7 µm, 50 mm × 4.6 mm) column at 40 °C. The mobile phase was a mixture of A and B, with a flow 0.6 mL/min, using the following gradients: from 95% A–5% B to 100% B in 3 min; 100% B for 3 min; from 100% B to 95% A–5% B in 1 min; and 95% A–5% B for 3 min. Purity is given as % of absorbance at 254 nm.

*General procedure A for the synthesis of the acyl chlorides*

To a solution of the carboxylic acid (0.50 mmol, 1 eq) in thionyl chloride (13.5 mmol, 27 eq) at room temperature a drop of DMF is added and the solution is refluxed for 2 h in a round bottom flask provided with a tube of anhydrous $CaCl_2$. Then, the excess of thionyl chloride is removed under vacuum. Toluene (5 mL) is added to the residue, and the remaining thionyl chloride is azeotropically removed under vacuum (twice), to give the acyl chloride as a waxy solid in quantitative yield.

*General procedure B for the synthesis of the carbohydrazides*

Under nitrogen atmosphere, 2-chlorophenylhydrazine (0.56 mmol, 1.1 eq) is solved in anhydrous THF (2 mL) at room temperature. Anhydrous triethylamine (1.03 mmol, 2 eq) is added and the mixture is stirred for 10 minutes. Next, the corresponding acyl chloride (0.5

mmol, 1 eq) in anh. THF (7 mL) is added and the mixture is left stirring overnight at room temperature. The purification procedure is specified below for each compound.

*UB-ALT-P30, N'-(2-chlorophenyl)adamantane-1-carbohydrazide*

It was synthesized as previously described[2]. The NMR spectra of the obtained sample was in agreement with the reported data[2].

*UB-MBX-46, N'-(2-chlorophenyl)-3,4,8,9-tetramethyltetracyclo[4.4.0.0$^{3,9}$.0$^{4,8}$]decane-1-carbohydrazide*

3,4,8,9-Tetramethyltetracyclo[4.4.0.0$^{3,9}$.0$^{4,8}$]decane-1-carbonyl chloride (128 mg, 0.51 mmol) was obtained following the general procedure **A** from 3,4,8,9-tetramethyltetracyclo[4.4.0.0$^{3,9}$.0$^{4,8}$]decane-1-carboxylic acid (synthesized as previously described)[9] and, without further purification, it was reacted with 2-chlorophenylhydrazine hydrochloride (100 mg, 0.56 mmol) in the presence of anh. triethylamine (143 µL, 1.02 mmol), following the general procedure **B**. The obtained precipitate was filtered off and the organic layer was concentrated *in vacuo* and subsequently washed with 2 mL of a mixture of DCM/hexane (1/9), yielding the desired product as a beige solid (110 mg, 61% yield), mp 217 °C. IR (ATR) ν: 3858, 3615, 3331, 2941, 2334, 2192, 2131, 2030, 1653, 1590, 1489, 1319, 1271, 1160, 1105, 1029, 748 cm$^{-1}$. $^{1}$H-NMR (400 MHz, $CDCl_3$) δ: 0.79 [dd, *J* = 11.6 Hz, *J'* = 2 Hz, 2H, 5(7)-$H_a$], 0.96 [d, *J* = 10.4 Hz, 2H, 2(10)-$H_a$], 0.968 [s, 6H, 3(9)-$CH_3$ or 4(8)-$CH_3$], 0.974 [s, 6H, 4(8)-$CH_3$ or 3(9)-$CH_3$], 1.84 [dd, *J* = 11.6 Hz, *J'* = 1.2 Hz, 2H, 5(7)-$H_b$] 1.99 [d, *J* = 11.2 Hz, 2H, 2(10)-$H_b$], 2.56 (broad s, 1H, 6-H), 6.49 (broad s, 1H, NH), 6.82 (m, 1H, 4'-H), 6.85 (dd, *J* = 8.0 Hz, *J'* = 1.2 Hz, 1H, 6'-H), 7.15 (m, 1H, 5'-H), 7.27 (dd, *J* = 8.0 Hz, *J'* = 1.6 Hz, 1H, 3'-H), 7.48 (s, 1H, NH). $^{13}$C-NMR (100.6 MHz, $CDCl_3$) δ: 15.67 [$CH_3$, C3(9)-$\underline{C}H_3$ or C4(8)-$\underline{C}H_3$], 15.73 [$CH_3$, C4(8)-$\underline{C}H_3$ or C3(9)-$\underline{C}H_3$], 37.7 (CH, C6), 38.3 [$CH_2$, C5(7)], 41.2 [$CH_2$, C2(10)], 45.2 [C, C3(9) or C4(8)], 45.7 [C, C4(8) or C3(9)], 48.4 (C, C1), 113.7 (CH, C6'), 120.1 (C, C2'), 121.5 (CH, C4'), 127.8 (CH, C5'), 129.7 (CH, C3'), 144.4 (C, C1'), 177.0 (C, CO). HRMS-ESI+ *m/z* $[M+H]^+$ calcd for $[C_{21}H_{28}ClN_2O]^+$: 359.1885, found: 359.1892. Elemental analysis: Calculated for $C_{21}H_{27}ClN_2O$: C 70.28, H 7.58, N 7.81. Found: C 70.16, H 7.62, N 7.80.

*UB-ALT-P36, N'-(2-chlorophenyl)-3,5-dimethyladamantane-1-carbohydrazide*

3,5-Dimethyladamantane-1-carbonyl chloride (217 mg, 0.96 mmol) was obtained following the general procedure **A** from commercially available 3,5-dimethyladamantane-1-carboxylic acid and, without further purification, it was reacted with 2-chlorophenylhydrazine hydrochloride (189 mg, 1.05 mmol) in the presence of anhydrous triethylamine (267 µL, 1.92 mmol), following the general procedure **B**. The solved was evaporated under vacuum and DCM (10 mL) was added. The organic phase was washed with 2 N aqueous HCl solution (2 x 5 mL), 2 N aqueous NaOH solution (2 x 5 mL) and brine (2 x 5 mL), dried over anhydrous $Na_2SO_4$, filtered and concentrated to dryness under vacuum. An analytical sample of the resulting solid was obtained by crystallization from DCM/Pentane (133 mg, 40 % yield), mp 170-171 °C. IR (ATR) ν: 3355, 3279, 2947, 2894, 2841, 1651, 1593, 1493, 1469, 1447, 1376, 1358, 1290, 1230, 1190, 1157, 1140, 1116, 1047, 1033, 918, 882, 842, 742, 704, 648, 630, 593, 555 cm$^{-1}$. $^{1}$H-NMR (400 MHz, $CDCl_3$) δ: 0.87 [s, 6H, 3(5)-$C\underline{H}_3$], 1.19 (m, 2H, 4-$H_2$), 1.38 [m, 4H, 6(10)-$H_2$], 1.54 [m, 4H, 2(9)-$H_2$], 1.76 (broad s, 2H, 8-$H_2$), 2.16 (m, 1H, 7-H), 6.44 (broad s, 1H, NH), 6.80-6.85 (cs, 2H, 4'-H and 6’-H), 7.14 (m, 1H, 5’-H), 7.27 (d, *J* = 7.6 Hz, 1H, 3’-H), 7.48 (s, 1H, NH). $^{13}$C-NMR (100.6 MHz, $CDCl_3$) δ: 29.3 (CH, C7), 30.5 [$CH_3$, C3(5)-$\underline{C}H_3$], 31.2 [C, C3(5)], 37.9 ($CH_2$, C8), 42.4 (C, C1), 42.8 [$CH_2$, C6(10)], 45.3 [$CH_2$, C2(9)], 50.7 ($CH_2$, C4), 113.6 (CH, C6’), 120.1 (C, C2’), 121.6 (CH, C4’), 127.7 (CH, C5’), 129.7 (CH, C3’), 144.3 (C, C1’), 177.3 (C, CO). HRMS-ESI+ *m/z* $[M+H]^+$ calcd for $[C_{19}H_{26}ClN_2O]^+$: 333.1728, found: 333.1736. Elemental analysis: Calculated for $C_{19}H_{25}ClN_2O$: C 68.56, H 7.57, N 8.42. Found: C 68.50, H 7.46, N 8.30.

*UB-ALT-P37, N'-(2-chlorophenyl)-3,5,7-trimethyladamantane-1-carbohydrazide*

3,5,7-Trimethyladamantane-1-carbonyl chloride (215 mg, 0.46 mmol) was obtained following the general procedure **A** from commercially available 3,5,7-trimethyladamantane-1-carboxylic acid and, without further purification, it was reacted with 2-chlorophenylhydrazine hydrochloride (94 mg, 0.52 mmol) in the presence of anhydrous triethylamine (133 µL, 0.96 mmol), following the general procedure **B**. The solved was evaporated under vacuum and DCM (10 mL) was added. The organic phase was washed with 2 N aqueous HCl solution (2 x 5 mL), 2 N aqueous NaOH solution (2 x 5 mL) and brine (2 x 5 mL), dried over anhydrous $Na_2SO_4$, filtered and concentrated to dryness under vacuum. An analytical sample of the isolated solid was obtained by crystallization from DCM/Pentane (112 mg, 67 % yield), mp 170-172 °C. IR (ATR) ν: 3277, 2943, 2890, 2836, 1661, 1597, 1544, 1475, 1454, 1410, 1373, 1355, 1292, 1230, 1208, 1156, 1139, 1116, 1099, 1067, 1044, 1035, 886, 851, 829, 774, 733, 667, 548 $cm^{-1}$. $^1$H-NMR (400 MHz, $CDCl_3$) δ: 0.88 [s, 9H, 3(5,7)-C$\underline{H}_3$], 1.13 [m, 6H, 4(6,10)-$H_2$] 1.48 [broad s, 6H, 2(8,9)-$H_2$], 6.43 (broad s, 1H, NH), 6.80-6.85 (cs, 2H, 4’-H and 6’-H), 7.15 (m, 1H, 5’-H), 7.27 (d, *J* = 8.0 Hz, 1H, 3’-H), 7.46 (s, 1H, NH). $^{13}$C-NMR (100.6 MHz, $CDCl_3$) δ: 30.2 [$CH_3$, C3(5,7)-$\underline{C}H_3$], 32.0 [C, C3(5,7)], 43.3 (C, C1), 44.7 [$CH_2$, C2(8,9)], 50.1 [$CH_2$, C4(6,10)], 113.6 (CH, C6’), 120.2 (C, C2’), 121.6 (CH, C4’), 127.7 (CH, C5’), 129.7 (CH, C3’), 144.3 (C, C1’), 177.2 (C, CO). HRMS-ESI+ *m/z* $[M+H]^+$ calcd for $[C_{20}H_{28}ClN_2O]^+$: 347.1885, found: 347.1894. Elemental analysis: Calculated for $C_{20}H_{27}ClN_2O$: C 69.25, H 7.85, N 8.08. Found: C 69.18, H 7.85, N 7.98.

*UB-ALT-P38, N'-(2-chlorophenyl)cubane-1-carbohydrazide*

Cubane-1-carbonyl chloride (55 mg, 0.33 mmol) was obtained following the general procedure **A** from commercially available cubane-1-carboxylic acid and, without further purification, it was reacted with 2-chlorophenylhydrazine hydrochloride (65 mg, 0.36 mmol) in the presence of anhydrous triethylamine (87 µL, 0.66 mmol), following the general procedure **B**. The solved was evaporated under vacuum and DCM (10 mL) was added. The organic phase was washed with 2 N aqueous HCl solution (2 x 5 mL), 2 N aqueous NaOH solution (2 x 5 mL) and brine (2 x 5 mL), dried over anhydrous $Na_2SO_4$, filtered and concentrated to dryness under vacuum. An analytical sample of the isolated solid was obtained by crystallization from DCM/Pentane (61 mg, 68 % yield), mp 189-190 °C. IR (ATR) ν: 3315, 3191, 3060, 2993, 2969, 1625, 1598, 1544, 1481, 1427, 1333, 1291, 1253, 1230, 1217, 1167, 1148, 1124, 1104, 1086, 1058, 1035, 972, 939, 885, 869, 839, 825, 743, 716, 693, 665, 645, 629, 583 $cm^{-1}$. $^1$H-NMR (400 MHz, $CDCl_3$) δ: mayor rotamer, 4.04-4.09 [cs, 4H, 3(5,7)-H and 4-H], 4.31 [m, 3H, 2(6,8)], 6.43 (broad s, 1H, NH), 6.80-6.90 (cs, 2H, 4’-H and 6’-H), 7.16 (m, 1H, 5’-H), 7.26-7.30 (cs, 2H, 3’-H and NH): minor rotamer, 3.92 [cs, 4H, 3(5,7)-H and 4-H], 4.16 [m, 3H, 2(6,8)], 6.25 (broad s, 1H, NH), 6.80-6.90 (m, 1H, 4’-H; overlapped with the other rotamer), 6.96 (d, *J* = 8.0 Hz, 1H, 6’-H), 7.22 (m, 1H, 5’-H), 7.26-7.30 (cs, 2H, 3’-H and NH; overlapped with the other rotamer). $^{13}$C-NMR

(100.6 MHz, $CDCl_3$) δ: mayor rotamer, 45.3 [CH, C3(5,7)], 48.1 (CH, C4), 49.6 [(CH, C2(6,8)], 56.2 (C, C1), 113.7 (CH, C6’), 120.0 (C, C2’), 121.6 (CH, C4’), 127.7 (CH, C5’), 129.6 (CH, C3’), 144.0 (C, C1’), 171.9 (C, CO); minor rotamer, 45.4 [CH, C3(5,7)], 47.3 (CH, C4), 49.7 [(CH, C2(6,8)], 57.0 (C, C1), 112.6 (CH, C6’), 118.3 (C, C2’), 121.3 (CH, C4’), 128.1 (CH, C5’), 129.7 (CH, C3’), 143.4 (C, C1’), 177.5 (C, CO). HRMS-ESI+ *m/z* $[M+H]^+$ calcd for $[C_{15}H_{14}ClN_2O]^+$: 273.0789, found: 273.0793. Elemental analysis: Calculated for $C_{15}H_{13}ClN_2O \cdot 0.05CH_2Cl_2$: C 65.26, H 4.77, N 10.11. Found: C 65.49, H 4.80, N 9.90. HPLC-UV purity: 96.6%

### Detailed molecular dynamics simulations methods

*Ligands Preparation*

Molecules UB-MBX-46, UB-ALT-P36, UB-ALT-P37, UB-ALT-P38, UB-MBX-P1, and UB-MBX-P2 were generated by building the corresponding cage alkyls on the coordinates of the UB-ALT-P30 using the Maestro interface (Schrödinger Release 2021-2: Maestro, Schrödinger, LLC, New York, NY, 2021). Subsequently, a minimization procedure was applied to all atoms of the protein complexes using the OPLS-2005 force field to ensure the stability of the complexes[10-12].

*Protein Preparation*

The cryo-EM structures of the f-hP2X7R in the apo closed state or the f-hP2X7R in complex with UB-ALT-P30 or UB-MBX-46 (with bound cholesterols) were utilized as starting models for MD simulations. In addition, models of the f-hP2X7R in complex with UB-ALT-P36, UB-ALT-P37, UB-ALT-P38, UB-MBX-P1, or UB-MBX-P2 were prepared from the experimental structure of UB-ALT-P30 bound to the f-hP2X7R (without bound cholesterols). The f-hP2X7R construct was generated from the cryo-EM structure of the hP2X7R in the apo closed state by deleting the cytoplasmic ballast. Both the N- and C-termini of the receptor were capped with acetyl and methylamino groups, respectively. The protein structure was optimized using the protein preparation module of Maestro program (Protein Preparation Wizard 2015-2; Schrödinger Release 2021-2: Maestro, Schrödinger, LLC, New York, NY, 2021)[13]. In this process, the bond orders and disulfide bonds were assigned, and missing hydrogen atoms and loops were added using Prime within Maestro[14,15]. As post preparation, all hydrogens in the protein complex were minimized employing the AMBER* force field via Maestro/Macromodel, with a distance-dependent dielectric constant set at 4.0[16,17]. The molecular mechanics minimizations were conducted using a conjugate gradient method, setting a convergence criterion threshold at 0.0001 kJ Å$^{-1}$ mol$^{-1}$. The ionization states of the compounds at pH 7.5 were confirmed based on the Epik program[18]. The protein complex was subjected in an all-atom minimization using the OPLS2005 force field with heavy atom RMSD values constrained to 0.30 Å[19].

*System setup for MD simulations*

Each model, prepared as mentioned above, was inserted in a pre-equilibrated hydrated POPC bilayer expanding 30 Å from the furthermost vertex of the protein to the edge of the simulation orthorhombic box in all axes. The protein was positioned with respect to the membrane plane (x,y plane) as suggested by the server "Orientations of Proteins in Membranes (OPM)" database using the System setup utility in Schrodinger Maestro software (Schrödinger Release 2020-2: Maestro, Schrödinger, LLC, New York, 2020) using the System Builder Wizard in Maestro software (Schrödinger Release 2020-4: Desmond Molecular Dynamics System, D. E. Shaw Research, New York, NY, 2021)[20]. Using the same utility, the TIP3P solvation model was applied while sodium and chloride ions were added randomly in the water phase to neutralize the system and reach the experimental salt concentration of 0.150 M NaCl[21]. The resulting lipid buffer contained approximately ~ 479,000 atoms, consisting of 584 POPC lipids and ~ 123,500 water molecules. The dimensions of the simulation box were 150×148×226 Å$^3$. Periodic boundary conditions were applied. We used the LEaP, the main program for preparing simulations in Amber Software, antechamber and Parmchk2 of AmberTools22 to assign the ff19sb parameters to model the protein, the lipid21 force field parameters to model the POPC lipids, the GAFF2 parameters to model the ligand, and the TIP3P model for waters and ions[5,6,8,22-24]. Partial charges for ligands were obtained using RESP fitting of the electrostatic potentials calculated with Gaussian03 at the Hartree-Fock (HF)/6-31G* level of theory and the antechamber of AmberTools22[4,25,26].

*Simulations protocol*

For each system, we executed an equilibration phase consisting of seven steps, starting with an energy minimization step. Systems were equilibrated by 5000 steps of energy minimization (1500 steps with the steepest descent algorithm and 3500 cycles with the conjugate gradient algorithm) in the presence of a harmonic restraint with a force constant of 10 kcal mol$^{-1}$ Å$^{-2}$ and 5 kcal mol$^{-1}$ Å$^{-2}$ on the heavy atoms of protein, ligand and lipid head groups. The next stage in MD simulation protocol is to allow the system to heat up from 0 K to 310 K using the Langevin thermostat (dynamics) for temperature control, as implemented in Amber22 program, employing a Langevin collision frequency of 2.0 ps and a friction coefficient constant at 1 ps$^{-1}$ [27,28]. In the next consecutive NVT and the three NPT simulation steps a gradual reduction in position restraints was implemented to maintain protein-ligand stability and ensure optimal lipid packing while heating was also applied. In the first step an 125 ps NVT simulation was applied in the presence of a harmonic restraint with a force constant of 10 kcal mol$^{-1}$ Å$^{-2}$ on all protein and ligand heavy atoms and 5 kcal mol$^{-1}$ Å$^{-2}$ on lipid head groups and in the second step an 125 ps NVT simulation was performed while the force constant was reduced to 5 and 2.5 kcal mol$^{-1}$ Å$^{-2}$, respectively.

In the third step, the temperature was raised to 310 K in a NPT$\gamma$ (with $\gamma$ = 10 dyn cm$^{1}$) simulation of 125 ps length in the presence of restraint with a force constant of 2.5 kcal mol$^{-1}$ Å$^{-2}$ on protein, ligand heavy atoms and 1.0 kcal mol$^{-1}$ Å$^{-2}$ on lipid head groups. For the remaining three 500 ps NPT simulation equilibration steps the applied restraints correspond to a force constant 1.0 and 0.5 kcal mol$^{-1}$ Å$^{-2}$ then 0.5 and 0.1 kcal mol$^{-1}$ Å$^{-2}$ and finally 0.1 and 0 kcal mol$^{-1}$ Å$^{-2}$, respectively. In the NPTγ simulations a surface tension 0 dyn cm$^{1}$ was implemented on *x*-*y* plane which gives pure semi-isotropic conditions. In each NPT$\gamma$ equilibration step for the pressure control the Berendsen barostat was used to adjust the density over the simulation time, with a target pressure of 1 bar and a 2 ps pressure relaxation time[29]. The temperature of 310 K was used in MD simulations to ensure that the membrane state is above the main phase transition temperature of 271 K for POPC bilayers[30].

Bonds involving hydrogen atoms were constrained by the SHAKE algorithm and a time step of 1 fs was used for the integration of the equations of motion for the first 2 NVT and the first NPT equilibration steps and for the rest of the NPT steps the time step was set at 2 fs with the leapfrog Verlet integrator[31,32]. Long range electrostatics were calculated using Particle-mesh Ewald summation (PME), with a 1 Å grid, and short-range non-bonding interactions were truncated at 12 Å with a continuum model long range correction applied for energy and pressure. The equilibration phase was followed by production MD simulation for 500 ns to 1 μs for the apo f-hP2X7R or its complexes with UB-ALT-P30 or the UB-MBX-46 using the same protocol as in the final equilibration step. Snapshots were recorded every 100 ps during the production phase. Within this simulation time, the RMSD (Cα) reached a plateau, and the systems were considered equilibrated and suitable for statistical analysis. Two MD simulation repeats were performed for each complex using the same starting structure and applying randomized velocities.

Short duration MD simulations (50 ns) were also performed for the complexes of UB-MBX-46, UB-ALT-P36, UB-ALT-P37, UB-ALT-P38, UB-MBX-P1, UB-MBX-P2 bound to the hP2X7R. These polycyclic hydrocarbons analogs of UB-ALT-P30 form complexes with the hP2X7R according to the MD simulations (see selected results in Supplementary Fig. 6 for the smallest analog, the cubyl derivative UB-ALT-P38).

Particle Mesh Ewald Molecular Dynamics (pmemd) is the primary engine for running MD simulations with AMBER22 software and the energy minimization step was performed using the Central Processing Unit (CPU) of the workstations by the implementation of pmemd[4]. The rest of the equilibration steps including the unrestraint production were run with AMBER22 software on RTX 4090 GPUs in lab workstations using pmemd.CUDA algorithm[4,33]. The pmemd.CUDA executable provides the ability to use NVIDIA GPUs to run the MD simulations.

*Analysis of MD simulations*

The visualization of the MD simulation trajectories was performed using VMD[34]. The analysis of all the MD simulations trajectories was performed by *ptraj* and *cpptraj* of AmberTools22[4,35]. For hydrogen bond interactions distance = 2.5 Å between donor and acceptor heavy atoms, and an angle ≥120$^o$ between donor-hydrogen-acceptor atoms and ≥ 90$^o$ between hydrogen-acceptor-bonded atoms were considered. Non-specific hydrophobic contacts were measured if the residue fell within 4.0 Å from a ligand's aromatic or aliphatic carbon, while π-π interactions were measured if two aromatic groups are stacked face-to-face or face-to-edge. Water-mediated interactions were measured if the distance between donor and acceptor atoms is 2.7 Å, the angle between donor-hydrogen-acceptor atoms is ≥ 110$^o$ and the angle between hydrogen-acceptor-bonded atoms is ≥ 80$^o$.

*Calculation of RBFEs*

For the TI/MD simulations, binding poses of UB-ALT-P36, UB-ALT-P37, UB-ALT-P38, UB-MBX-P1, UB-MBX-P2, UB-MBX-46 aligned with UB-ALT-P30 in complex with the hP2X7R from MD simulations were used as starting structures for the alchemical calculations described in Table 1. TI/MD calculations were also performed for the ligands in solution.

The setup procedure was the same as previously reported for the MD simulations with Amber22 program[36]. The bond constraint SHAKE algorithm was disabled for TI mutations in AMBER GPU-TI module pmemdGTI, and therefore a time step of 1 fs was used for all MD simulations[31,37]. Long range electrostatics were calculated using PME, with a 1 Å grid, and short-range non-bonding interactions were truncated at 12 Å with a continuum model long range correction applied for energy and pressure[38]. The ff19sb was used to model the protein, the lipid21 force field to model the POPC lipids, the GAFF2 to model the ligand, and the TIP3P model for waters and ions[5,6,8,22-24].

Thus, initial geometries were minimized using 20,000 steps of steepest descent minimization at $\lambda$ = 0.5. These minimized geometries were then used for simulations at all $\lambda$ values. Eleven $\lambda$ values were applied, equally spaced between 0.0 to 1.0. Each MD simulation was heated to 310 K for 500 ps using the Langevin thermostat (dynamics) for temperature control, as implemented in Amber22 program employing a Langevin collision frequency of 2.0 $ps^{1}$ in the presence of harmonic restraint with force constant 10 kcal $mol^{-1}$ $Å^{-2}$ on all membrane, protein, and ligand atoms[4]. The temperature of 310 K was used in MD simulations to ensure that the membrane state is above the main phase transition temperature of 271 K for POPC bilayers[30]. The Berendsen barostat was used to adjust the density over 500 ps at constant pressure (NPTγ) (with $\gamma$ = 10 dyn $cm^{1}$), with a target pressure of 1 bar and a 2 ps coupling time[29]. Then, the 500 ps of constant volume equilibration (NVT) was followed by 2 ns NVT production simulation without restraints. Energies were recorded every 1 ps, and coordinates were saved every 10 ps. Production simulations recalculated the potential energy at each $\lambda$ value every 1 ps for later analysis with MBAR[38,39].

For each alchemical calculation was applied dual topology and the 1-step protocol was performed which includes disappearing one ligand and appearing the other ligand simultaneously, and the electrostatic and van der Waals interactions are scaled simultaneously using softcore potentials from real atoms that are transformed into dummy atoms[3,40]. Two repeats were performed for the TI/MD calculation for each alchemical transformation and the resultant $\Delta\Delta G_{b,\ TI/MD}$ values are shown in **Table 1**.

The final states 0 and 1 of the alchemical calculations ligand 0 → ligand 1, i.e., structures of ligand 0-hP2X7R and ligand 1-hP2X7R complexes as resulted from the alchemical transformations were compared with these complexes structure resulted from converged 50 ns-MD simulations (2 repeats). This was performed to certify that the 2 ns MD simulation for each λ-state during the alchemical calculations was enough for the complexes 0— hP2X7R and 1— hP2X7R to converge to same structure with 50 ns-MD simulations (Supplementary Fig. 6).

Experimental relative binding free energies for the alchemical transformations from UB-ALTP-30 (ligand 0) to another ligand 1 were estimated using the experimental $pIC_{50,0}$ / $pIC_{50,1}$ values depicted in Table 1 to approximate $pK_{d,0}$ / $pK_{d,1}$ according to the following equation ($T$ = 298 K)

$$\Delta G_{b,0\rightarrow 1,exp} = -1.9872\, T\, (pK_{d,0} - pK_{d,1}) \sim \Delta G_{b,0\rightarrow 1,exp} = -1.9872\, T\, (pIC_{50,0} - pIC_{50,1}) \qquad (1)$$

*Calculation of RSFE*

To calculate the relative solvation free energy (RSFE) the TI/MD simulations for the alchemical calculation **UB-ALT-P30 → UB-MBX-46** was performed in the water and gas phase. The simulation box extends 15×15×17.5 $Å^3$ from the furthermost vertex of the ligand to the edge of the simulation orthorhombic box in all axes containing 3889 waters. The same parameters and simulation protocol were used as described previously for the alchemical transformation calculations.

### Supplemental References


1. Barniol-Xicota, M. et al. Escape from adamantane: Scaffold optimization of novel P2X7 antagonists featuring complex polycycles. *Bioorganic & Medicinal Chemistry Letters* **27**, 759-763 (2017).
2. Nelson, D.W. et al. Structure−Activity Relationship Studies on N'-Aryl Carbohydrazide P2X7 Antagonists. *Journal of Medicinal Chemistry* **51**, 3030-3034 (2008).
3. Song, L.F., Lee, T.-S., Zhu, C., York, D.M. & Merz, K.M., Jr. Using AMBER18 for Relative Free Energy Calculations. *Journal of Chemical Information and Modeling* **59**, 3128-3135 (2019).
4. Case, D.A. et al. AmberTools. *Journal of Chemical Information and Modeling* **63**, 6183-6191 (2023).
5. Tian, C. et al. ff19SB: Amino-Acid-Specific Protein Backbone Parameters Trained against Quantum Mechanics Energy Surfaces in Solution. *Journal of Chemical Theory and Computation* **16**, 528-552 (2020).
6. Dickson, C.J., Walker, R.C. & Gould, I.R. Lipid21: Complex Lipid Membrane Simulations with AMBER. *Journal of Chemical Theory and Computation* **18**, 1726-1736 (2022).
7. He, X. et al. Fast, Accurate, and Reliable Protocols for Routine Calculations of Protein–Ligand Binding Affinities in Drug Design Projects Using AMBER GPU-TI with ff14SB/GAFF. *ACS Omega* **5**, 4611-4619 (2020).
8. He, X., Man, V.H., Yang, W., Lee, T.-S. & Wang, J. A fast and high-quality charge model for the next generation general AMBER force field. *The Journal of Chemical Physics* **153**, 114502 (2020).
9. Leiva, R. et al. Pharmacological and Electrophysiological Characterization of Novel NMDA Receptor Antagonists. *ACS Chemical Neuroscience* **9**, 2722-2730 (2018).
10. Jorgensen, W.L., Maxwell, D.S. & Tirado-Rives, J. Development and Testing of the OPLS All-Atom Force Field on Conformational Energetics and Properties of Organic Liquids. *Journal of the American Chemical Society* **118**, 11225-11236 (1996).
11. Jorgensen, W.L. & Tirado-Rives, J. The OPLS [optimized potentials for liquid simulations] potential functions for proteins, energy minimizations for crystals of cyclic peptides and crambin. *Journal of the American Chemical Society* **110**, 1657-1666 (1988).
12. Shivakumar, D. et al. Prediction of Absolute Solvation Free Energies using Molecular Dynamics Free Energy Perturbation and the OPLS Force Field. *Journal of Chemical Theory and Computation* **6**, 1509-1519 (2010).
13. Madhavi Sastry, G., Adzhigirey, M., Day, T., Annabhimoju, R. & Sherman, W. Protein and ligand preparation: parameters, protocols, and influence on virtual screening enrichments. *Journal of Computer-Aided Molecular Design* **27**, 221-234 (2013).
14. Jacobson, M.P., Friesner, R.A., Xiang, Z. & Honig, B. On the Role of the Crystal Environment in Determining Protein Side-chain Conformations. *Journal of Molecular Biology* **320**, 597-608 (2002).

15. Jacobson, M.P. et al. A hierarchical approach to all-atom protein loop prediction. *Proteins: Structure, Function, and Bioinformatics* **55**, 351-367 (2004).
16. Mohamadi, F. et al. Macromodel—an integrated software system for modeling organic and bioorganic molecules using molecular mechanics. *Journal of Computational Chemistry* **11**, 440-467 (1990).
17. Watts, K.S., Dalal, P., Tebben, A.J., Cheney, D.L. & Shelley, J.C. Macrocycle Conformational Sampling with MacroModel. *Journal of Chemical Information and Modeling* **54**, 2680-2696 (2014).
18. Shelley, J.C. et al. Epik: a software program for pKaprediction and protonation state generation for drug-like molecules. *Journal of Computer-Aided Molecular Design* **21**, 681-691 (2007).
19. Kaminski, G.A., Friesner, R.A., Tirado-Rives, J. & Jorgensen, W.L. Evaluation and Reparametrization of the OPLS-AA Force Field for Proteins via Comparison with Accurate Quantum Chemical Calculations on Peptides. *The Journal of Physical Chemistry B* **105**, 6474-6487 (2001).
20. Lomize, M.A., Pogozheva, I.D., Joo, H., Mosberg, H.I. & Lomize, A.L. OPM database and PPM web server: resources for positioning of proteins in membranes. *Nucleic Acids Research* **40**, D370-D376 (2012).
21. Jorgensen, W.L., Chandrasekhar, J., Madura, J.D., Impey, R.W. & Klein, M.L. Comparison of simple potential functions for simulating liquid water. *The Journal of Chemical Physics* **79**, 926-935 (1983).
22. Joung, I.S. & Cheatham, T.E., III. Determination of Alkali and Halide Monovalent Ion Parameters for Use in Explicitly Solvated Biomolecular Simulations. *The Journal of Physical Chemistry B* **112**, 9020-9041 (2008).
23. Sengupta, A., Li, Z., Song, L.F., Li, P. & Merz, K.M., Jr. Parameterization of Monovalent Ions for the OPC3, OPC, TIP3P-FB, and TIP4P-FB Water Models. *Journal of Chemical Information and Modeling* **61**, 869-880 (2021).
24. Li, P., Song, L.F. & Merz, K.M., Jr. Systematic Parameterization of Monovalent Ions Employing the Nonbonded Model. *Journal of Chemical Theory and Computation* **11**, 1645-1657 (2015).
25. Bayly, C.I., Cieplak, P., Cornell, W. & Kollman, P.A. A well-behaved electrostatic potential based method using charge restraints for deriving atomic charges: the RESP model. *Journal of physical chemistry (1952)* **97**, 10269-10280 (1993).
26. Davidson, E.R. & Feller, D. Basis set selection for molecular calculations. *Chemical Reviews* **86**, 681-696 (1986).
27. Izaguirre, J.A., Reich, S. & Skeel, R.D. Longer time steps for molecular dynamics. *The Journal of Chemical Physics* **110**, 9853-9864 (1999).
28. Izaguirre, J.A., Catarello, D.P., Wozniak, J.M. & Skeel, R.D. Langevin stabilization of molecular dynamics. *The Journal of Chemical Physics* **114**, 2090-2098 (2001).
29. Berendsen, H.J.C., Postma, J.P.M., van Gunsteren, W.F., DiNola, A. & Haak, J.R. Molecular dynamics with coupling to an external bath. *The Journal of Chemical Physics* **81**, 3684-3690 (1984).

30. Koynova, R. & Caffrey, M. Phases and phase transitions of the phosphatidylcholines. *Biochimica et Biophysica Acta (BBA) - Reviews on Biomembranes* **1376**, 91-145 (1998).
31. Ryckaert, J.-P., Ciccotti, G. & Berendsen, H.J.C. Numerical integration of the cartesian equations of motion of a system with constraints: molecular dynamics of n-alkanes. *Journal of Computational Physics* **23**, 327-341 (1977).
32. Verlet, L. Computer "Experiments" on Classical Fluids. I. Thermodynamical Properties of Lennard-Jones Molecules. *Physical Review* **159**, 98-103 (1967).
33. Salomon-Ferrer, R., Götz, A.W., Poole, D., Le Grand, S. & Walker, R.C. Routine Microsecond Molecular Dynamics Simulations with AMBER on GPUs. 2. Explicit Solvent Particle Mesh Ewald. *Journal of Chemical Theory and Computation* **9**, 3878-3888 (2013).
34. Humphrey, W., Dalke, A. & Schulten, K. VMD: Visual molecular dynamics. *Journal of Molecular Graphics* **14**, 33-38 (1996).
35. Roe, D.R. & Cheatham, T.E., III. PTRAJ and CPPTRAJ: Software for Processing and Analysis of Molecular Dynamics Trajectory Data. *Journal of Chemical Theory and Computation* **9**, 3084-3095 (2013).
36. Case, D.A. et al. The Amber biomolecular simulation programs. *Journal of Computational Chemistry* **26**, 1668-1688 (2005).
37. Lee, T.-S., Hu, Y., Sherborne, B., Guo, Z. & York, D.M. Toward Fast and Accurate Binding Affinity Prediction with pmemdGTI: An Efficient Implementation of GPU-Accelerated Thermodynamic Integration. *Journal of Chemical Theory and Computation* **13**, 3077-3084 (2017).
38. Essmann, U. et al. A smooth particle mesh Ewald method. *The Journal of Chemical Physics* **103**, 8577-8593 (1995).
39. Shirts, M.R. & Pande, V.S. Comparison of efficiency and bias of free energies computed by exponential averaging, the Bennett acceptance ratio, and thermodynamic integration. *The Journal of Chemical Physics* **122**, 144107 (2005).
40. Mey, A.S.J.S. et al. Best Practices for Alchemical Free Energy Calculations [Article v1.0]. *Living Journal of Computational Molecular Science* **2**, 18378 (2020).
